# Co-evolution of a near-infrared aptamer:dye system for live-cell super-resolution RNA imaging

**DOI:** 10.1101/2025.09.18.677088

**Authors:** Simon Fürbacher, Pia Doll, Jingye Zhang, Laura Lange, Niklas van den Bergh, Yaqing Zhang, Erika Vitiello, Franziska Grün, Murat Sunbul, Andres Jäschke

## Abstract

Despite their advantageous properties for live-cell imaging and super-resolution microscopy, high-performance silicon rhodamine (SiR) near-infrared (NIR) probes are still rarely employed in RNA imaging via fluorescent light-up aptamers (FLAPs). Here, we developed the SiRiuS:SiR-5 system through a combined evolutionary approach: evolving the aptamer via fluorescence-activated cell sorting (FACS), along with targeted mutations, truncations, and rational design, and improvement of the dye by systematic chemical derivatization. This resulted in an aptamer:dye pair with high fluorogenicity and photostability, specifically optimized for visualization of RNAs in mammalian live cells. Our system demonstrates strong fluorescence enhancement in live-cell imaging, enabling time resolved imaging of dynamic processes such as stress-granule formation. Notably, we validate its application in STED super-resolution microscopy, establishing it as a powerful NIR imaging platform for RNA structures below the refraction limit. Its orthogonality to existing FLAPs operating in the yellow-orange spectrum further broadens its versatility for exploring complex RNA dynamics in live cells.

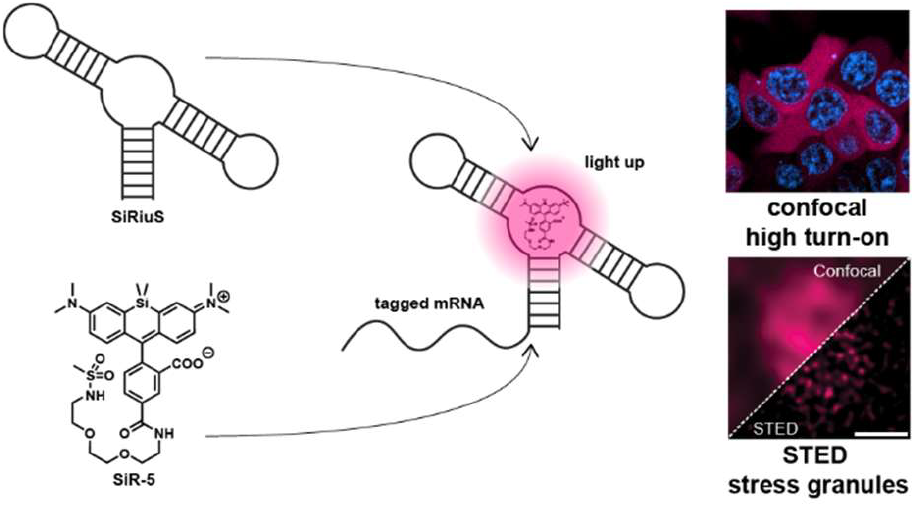

## INTRODUCTION

Fluorescence imaging has dramatically transformed our ability to observe and analyze complex cellular dynamics by providing detailed spatial and temporal information about biomolecules in living cells. This technology has been pivotal in advancing our understanding of critical processes such as gene expression, signal transduction, and cellular metabolism. Key to this progress was the development of fluorescent proteins that allow for the tagging of proteins across the visible spectrum, facilitating numerous groundbreaking techniques and discoveries. However, the lack of naturally fluorescent RNAs has hampered progress in RNA imaging and visualization^1^.

To overcome this obstacle, various strategies have been developed. Fluorescent in situ hybridization (FISH) utilizes complementary nucleic acid probes tagged with fluorophores, providing high specificity but necessitating cell fixation, which is unsuitable for live imaging^2,3^. Molecular Beacons—self-quenching probes—enable live-cell imaging but require microinjection or transfection^4,5^. The MS2-MCP-GFP system was the first widely used method for tracking RNA sequences in living cells using GFP-tagged RNA binding proteins^6,7^. However, this method’s use of large protein tags can disrupt RNA’s normal metabolism, localization, and interactions, and necessitates MCP co-expression, limiting applicability^8^.

In response to these limitations, fluorescent light-up aptamers (FLAPs) have emerged as a direct and genetically encodable approach for RNA imaging over the last 20 years. FLAPs are structured RNA strands that bind with high affinity and selectivity to fluorogenic dyes, thereby inducing dye fluorescence by different mechanisms^9^. Unlike indirect methods, FLAPs are genetically encodable tags and permit RNA imaging without fixation or the need for co-expressed proteins, thus simplifying experimentation, and reducing cellular metabolic burden. They also facilitate multicolor imaging and are particularly advantageous for live cell imaging due to their non-invasive attributes and broad coverage of the visible spectrum.

The first-generation FLAPs, Spinach^10^ and its successor Spinach2^11^, bind DMHBI, a mimic of the GFP chromophore, emitting green fluorescence. Subsequent developments, such as Broccoli^12,13^ offer robust aptamers with stable folding in cells, while Corn^14^, Squash^15^, RhoBAST^16^, Riboglow^17^ and Pepper^18^ and Clivia provide imaging of RNAs across the green to red spectrum with enhanced photostability and brightness. Notably, Pepper and its HBC dyes achieve exceptional brightness in their respective spectra^19^.

In contrast to the extensive visible light FLAP repertoire, there are only very few systems for RNA imaging in the near-infrared (NIR) range of the spectrum. NIR fluorescence (650-750 nm) is advantageous for minimizing cellular autofluorescence, reducing phototoxicity and bleaching, and enhancing tissue penetration^20–22^. It also supports multi-channel imaging by minimizing spectral overlap with other FLAPs or fluorescent proteins. While the DIR2^23^ and MGA^24^ aptamers have shown NIR fluorescence turn-on, their application in mammalian live-cell RNA imaging remains to be demonstrated. Riboglow permitted NIR imaging of tagged mRNAs^17^ in live mammalian cells, however only at conventional resolution.

In 2019, our lab introduced SiRA^26^, an aptamer that binds the silicon rhodamine derivative SiR-NH_2_ (Figure 1a)^27^, a near-infrared fluorophore characterized by exceptional photostability, brightness, and cell permeability. The system’s fluorogenicity is based on spirolactonization (Figure 1b), favoring the fluorescent form upon binding. It demonstrated impressive *in vitro* performance, enabling significant fluorescence enhancement, nanomolar binding, and successful NIR imaging in live *E. coli* with minimal photobleaching and phototoxicity. SiRA also performed well in STED microscopy, making the SiRA system the first FLAP cpable of super-resolution imaging^26^. However, when applied in mammalian live-cells, we found the SiRA:SiR-NH_2_ system to show minimal fluorescence turn-on, even with overexpressed circular RNA^28^, and problematic dye aggregation (Figure S1a). These observed properties have therefore severely limited the usability of SiRA-SiR-NH_2_ as a NIR FLAP system for RNA imaging in living eukaryotic cells.

**Figure 1.**
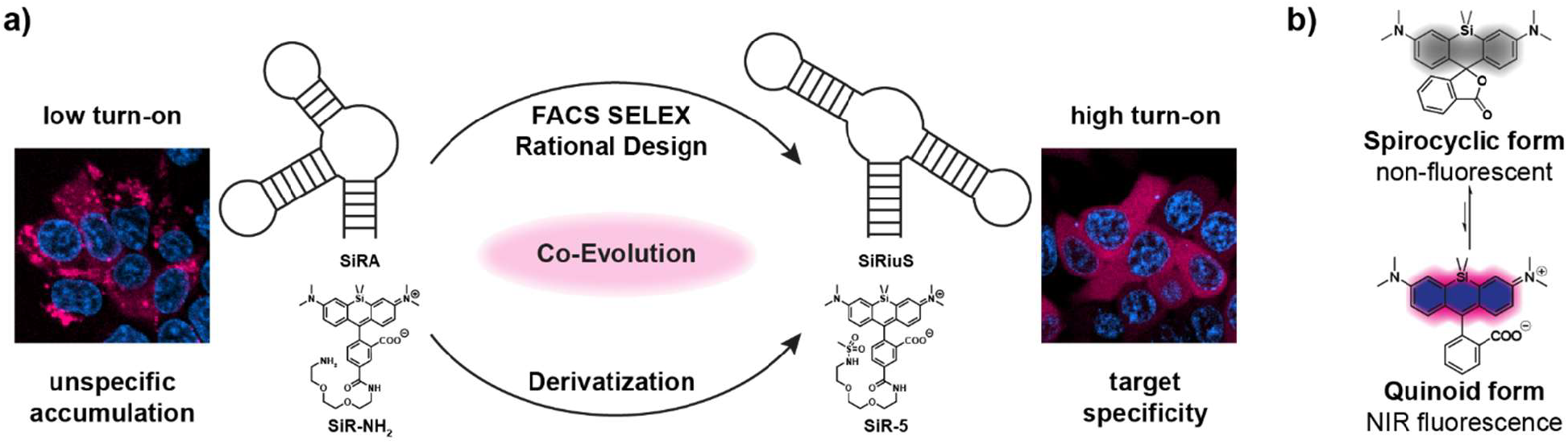
Co-evolution of an improved SiR-binding aptamer and its cognate dye. a) Schematic representation of the SiRA Aptamer and its cognate dye^26^(left) and the newly co-evolved SiRiuS:SiR-5 system and their performance in cells. b) Intramolecular spirolactonization of silicon rhodamines.

Here, we report the development of a novel FLAP:dye pair for NIR imaging in mammalian live cells. Addressing the limitations of conventional affinity-based SELEX, we utilized fluorescence-activated cell sorting (FACS) to enrich directly for fluorescence^12,29^, followed by screening in mammalian cells, chemical probing, mutation, and rational design, resulting in the SiRiuS (Superior infrared RNA imager using Silicon Rhodamine) aptamer. Simultaneously, we tackled the problem of dye accumulation through derivatization without compromising binding properties. The SiRiuS:SiR-5 aptamer:dye combination exhibited uniform cytosolic distribution and high fluorescence enhancement in live mammalian cells, facilitating live-cell mRNA imaging across four different mammalian cell lines and confirming its utility in dynamic cellular contexts. Leveraging this capability, we were able to visualize the time-dependent formation of stress granules in living cells, further highlighting the system’s potential for studying transient RNA-associated processes. Additionally, we demonstrate its application in STED super-resolution microscopy, achieving a resolution of about 40 nm.

## RESULTS AND DISCUSSION

### Utilizing FACS for fluorescence-based selection of SiR binding aptamers

In our pursuit to enhance the performance of the SiRA aptamer, we recognized the limitations of relying solely on high-affinity binding as determined by standard SELEX, which initially evolved the aptamer from a synthetic combinatorial RNA library. This library, encompassing approximately 10^14^ different sequences, provided a robust foundation; however, high-affinity binding alone does not guarantee significant fluorescence enhancement, as evidenced by our previous research and that of others^30,31^. Consequently, we sought alternative strategies that prioritize direct selection based on fluorescence intensity. Fluorescence-Activated Cell Sorting (FACS) emerged as a promising choice, having been successfully utilized for selecting aptamers against protein targets^29^, as well as the Broccoli FLAP through fluorescence guidance ^12^. This approach not only allows for the enrichment of fluorophore-activating aptamers but also addresses issues related to aptamer misfolding, poor thermostability, and ion dependence, which are common in constructs identified solely through *in vitro* methods^11^. Therefore, we implemented FACS to isolate silicon rhodamine binders directly expressed in cells rather than relying on *in vitro* production.

We selected *E. coli* BL21(DE3) cells as the host organism because they are easily manipulated genetically and allow high-level expression of aptamers. DNA pools from rounds 7 (L1) and 14 (L2) of the original SiRA SELEX were used as input due to their strong enrichment in SiR-binding sequences. To protect the aptamers from nucleolytic degradation, the library was embedded in a tRNA^Lys^ scaffold^32^, and the resulting tRNA^Lys^–library construct was cloned downstream of the T7 promoter in a pET28 vector (Table S1). In this expression system, high levels of aptamers were obtained upon IPTG induction.

The plasmid libraries were first transformed into *E. coli*, yielding approximately 10^5^ plasmid-containing bacteria. Transformed cells were then cultivated, and aptamer transcription was induced with 1 mM IPTG. However, *E. coli* carrying pET28 vectors showed poor survival and growth after IPTG treatment (Figure S2 b,c), raising concerns about the feasibility of this approach for iterative aptamer selection, as candidate sequences might be lost. Since the original SiRA aptamer performed reliably in both live and fixed *E. coli*, we incorporated a fixation step with 1% paraformaldehyde (PFA) prior to sorting. After fixation, bacteria expressing the aptamer libraries were incubated with 1 µM fluorogenic SiR-NH_2_ dye and sorted based on their fluorescence signal. The sorting gate was set above the fluorescence of a negative control population and adjusted each round to fine-tune selection pressure. From ~10,000 positively sorted cells, aptamer sequences were recovered by PCR to generate an enriched DNA pool for the next round. The final FACS-based selection cycle (Figure 2a) thus comprised four steps: cloning, transformation, sorting, and PCR amplification.

**Figure 2.**
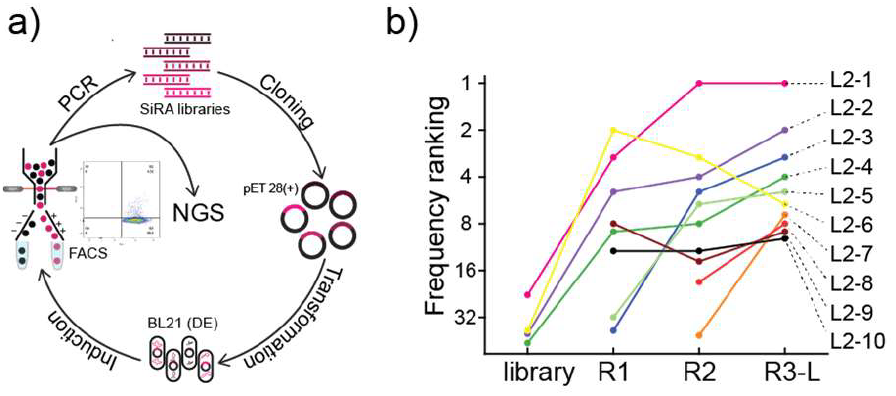
FACS SELEX principle and workflow. a) The DNA from rounds 7 and 14 of the original SiRA selection was cloned into pET 28(+) vectors and transformed into *E. coli*. After induction, fixed cells were sorted by FACS and plasmids from positive cells were recovered. PCR-amplified inserts started the next cycle. b) Top ten candidates after R3-L were analyzed in respect to their copy number enrichment per round and further characterized for affinity and turn-on.

FACS SELEX was carried out over three iterative rounds (R1-R3). Additionally, an alternative third round (R3-L) was performed with a five-fold reduction in dye concentration (200 nM) to further increase selection pressure. A notable increase in the percentage of positive cells (those within the sorting gate) was observed over the selection process, as figures rose from 1.3% to 3.7% for L1, and from 0.4% to 4.5% for L2. These results signified a successful enrichment of turn-on inducing sequences through the FACS SELEX process. As expected, the reduced dye concentration in R3-L led to a decrease in positive cell counts (Figure S3).

Following FACS SELEX, all four selection rounds and the initial pools for L1 and L2 were subjected to Illumina sequencing, to assess sequence abundance and enrichment (Figures 2b and S4). The top ten most abundant sequences emerging from the final round (R3-L) of FACS SELEX in both libraries were chosen for further characterization. Six sequences appeared in the top 10 group of both libraries. Therefore, 14 individual sequences were scrutinized. These selected sequences not only demonstrated high abundance but also showed enrichment, suggesting their competitive performance during the FACS-based selections.

The individual aptamers were synthesized through *in vitro* transcription, and their dissociation constants (K_D_) and fluorescence turn-on values were measured. Remarkably, eleven aptamer candidates exhibited nanomolar affinity for SiR-NH_2_, with three showcasing K_D_ values below 100 nM. Additionally, twelve candidates demonstrated considerable *in vitro* turn-on, with the highest value reaching an 8.2-fold increase (Table S2). From these candidates, the four with the highest affinity were designated for evaluation of their *in cellulo* imaging properties in HEK293T cells: L2-1 (K_D_ = 25 nM), L2-7 (K_D_ = 85 nM), L2-8 (K_D_ = 53 nM), and L1-8 (K_D_ = 105 nM, Table S2). These aptamers, along with RhoBAST as a negative control, were overexpressed in circularized form using the Tornado expression system. HEK293T cells were then imaged in the presence of 1 µM SiR-NH_2_ dye (Figure S5). Unexpectedly, the candidate with the highest *in vitro* affinity, L2-1, displayed the lowest fluorescence intensities *in cellulo*. In contrast, aptamer L2-8 showed a marked improvement in *in cellulo* signal-to-noise ratio and was thus chosen for further exploration and optimization in subsequent studies.

### Optimization of selected aptamer L2-8 for *in cellulo* imaging

The secondary structure prediction of aptamer L2-8, within the tRNA_Lys_ scaffold using the Zuker algorithm33, revealed a four-way junction configuration featuring four helices, as well as internal and apical loops, creating a total length of 115 nucleotides (Figure 3a). Helix H1 originates from the tRNA scaffold, and the prediction suggests a kissing loop interaction between two complementary 7-nucleotide sequences in loops L2 and L4 (Figure 3a). Notably, the constant SELEX primers A and B (Figure 3d in blue and Table S1) play critical roles in the formation of secondary structures, which impose limitations on truncation possibilities, unlike the original SiRA aptamer.

**Figure 3.**
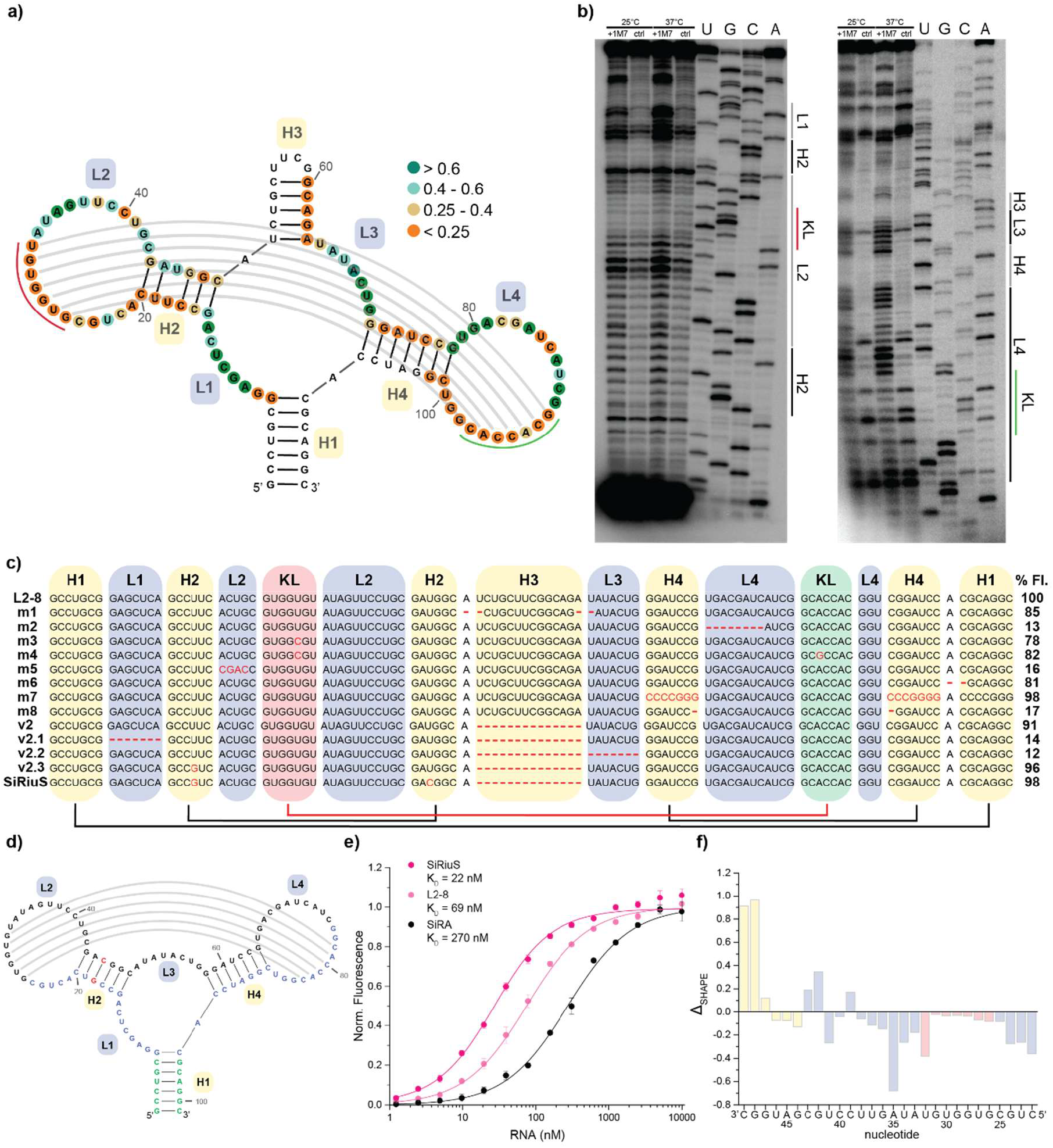
Development of the optimized SiR-binding aptamer SiRiuS. a) SHAPE reactivities plotted to aptamer structure. Analysis revealed two constricted sequences in L2 and L4 perfectly complementing each other. With the nucleotides of these complements being completely restricted according to SHAPE reactivity, we suspect a kissing loop interaction between them (marked in red and green). b) SHAPE probing of the candidate L2-8. Due to the length of the sequence, two primers have been designed, one to bind at the 3’ end of the aptamer, one in the tetraloop (for sequences, see Table S1. SHAPE gels show 1M7 reactivities with alignment to a full nucleotide resolved sequencing ladder. Aptamers were freshly folded for analysis and subsequently incubated at 25°C or 37°C with 1M7. After reverse transcription with radioactively labeled primer, cDNA was run for 4 h on a 60 cm 20% PAGE gel. c) Mutation variants were designed based on these findings. Fluorescent values are given in respective value to L2-8. d) Structure of the final aptamer version SiRiuS. Fixed mismatch in H2 is marked in red. Nucleotides originating from the SELEX primers are marked in light blue. Nucleotides that were added as a base stem due to the open structure of L2-8 are marked in green e) Affinity curves of the original aptamer L2-8 and final version SiRiuS compared to SiRA. f) Differences of SHAPE reactivities in the suspected binding pocket with and without ligand added to the probing reaction.

To test these predictions, SHAPE probing^34^ of L2-8 was conducted, assessing nucleotide accessibility to the chemical acylation reagent 1-methyl-7-nitroisatoic anhydride (1M7)^35^. This method provided insight into the aptamer’s structural nuances, confirming the predicted helices through low reactivity in the stem regions. An especially low SHAPE reactivity was observed in nucleotides implicated in the kissing loop interaction (Figure 3a,b). Additionally, most predicted loop nucleotides showed high reactivity with 1M7, indicating notable flexibility. Interestingly, the unreactive state of nucleotides between H2 and the kissing loop (5’-ACUGC-3’) suggests potential tertiary interactions or steric hindrance, hinting at a dye binding pocket. This is further supported by the decrease of SHAPE reactivity of said nucleotides when repeating the experiment in presence of 2 µM SiR-NH_2_ (Figures 3f and S6). The consistent SHAPE patterns at both 25°C and 37°C (Figure 3b) underscore the aptamer’s stable folding at physiological temperatures, indicating its potential suitability for live-cell imaging applications.

Further analysis involved designing mutated and truncated variants of L2-8, measuring their *in vitro* fluorescence turn-on values. Unexpected resilience in fluorescence turn-on was observed with truncation (Figure 3c, mutant m1) of nucleotides in the constant tetraloop stem, part of the original SELEX constant region. Shortening parts of the open loop L4 not involved in the kissing loop led to almost complete loss of turn-on (m2). The mutation of a nucleotide in the kissing loop (m3) significantly reduced turn-on, although restoring complementarity (m4) slightly improved fluorescence. Permutating the nucleotides connecting the kissing loop with helix H2 (m5) led to loss of over 80% of turn on. This led us to believe, these nucleotides to be involved in binding. Deleting the connecting A between H4 and H1 also decreased performance (m6). While H4 could be completely replaced by a CCCCGGG stem while retaining turn-on (m7), shortening this helix by just one base pair led to almost complete loss of fluorescent turn on(m8), confirming our findings through SHAPE. Building on the findings of m1, we removed H3 entirely (v2), with the truncated version exhibiting a comparable K_D_ to the initial aptamer (Figure S7). In our strive to find a minimal version of the aptamer, we based subsequent mutants on this second version of the initial aptamer (version 2). Removing sections of loops L1 (v2.1) and L3 (v2.2) led to a complete loss of fluorescence turn-on. Introducing a base pair by altering a U-U mismatch in stem 2 (v2.3) retained turn-on. Enhancements continued with the replacement of a weak G-U base pair with a G-C base pair, boosting affinity significantly (Figure 3e).The results emphasize the critical role of the kissing loops in preserving the aptamer’s functional structure. Without further truncation or permutation possibilities, the optimized aptamer, dubbed SiRiuS (Superior infrared RNA imager using Silicon Rhodamine), was finalized (Figure 3d). SiRiuS exhibits a K_D_ of 22 nM towards SiR-NH_2_ and an *in vitro* fluorescence turn-on of 8.4-fold, representing significant improvements over SiRA and L2-8 (Figure 3e).

To facilitate RNA tagging, which necessitates multiple repeats of the aptamer, we engineered three synonymous versions of SiRiuS, altering stem regions to prevent crossfolding while preserving the overall structure. The GC content remained consistent across versions, and *in silico* analyses confirmed that they folded into identical structures with similar Gibbs free energies (Figure S8).

### Dye evolution to facilitate accumulation-free SiR signal in mammalian live cells

In addition to exhibiting low fluorescence turn-on (Figure S1a,b), the original SiRA:SiR-NH_2_ system demonstrated the formation of brightly fluorescent particles throughout live HEK293T cells (Figures 4a and S1a). Similar results were obtained when SiR-NH_2_ dye was administered to cells expressing orthogonal aptamers such as RhoBAST (Figures 4b and S1a). This observation highlighted the necessity for structural optimization of the dye to enable effective use of these aptamer systems in mammalian live cells.

**Figure 4.**
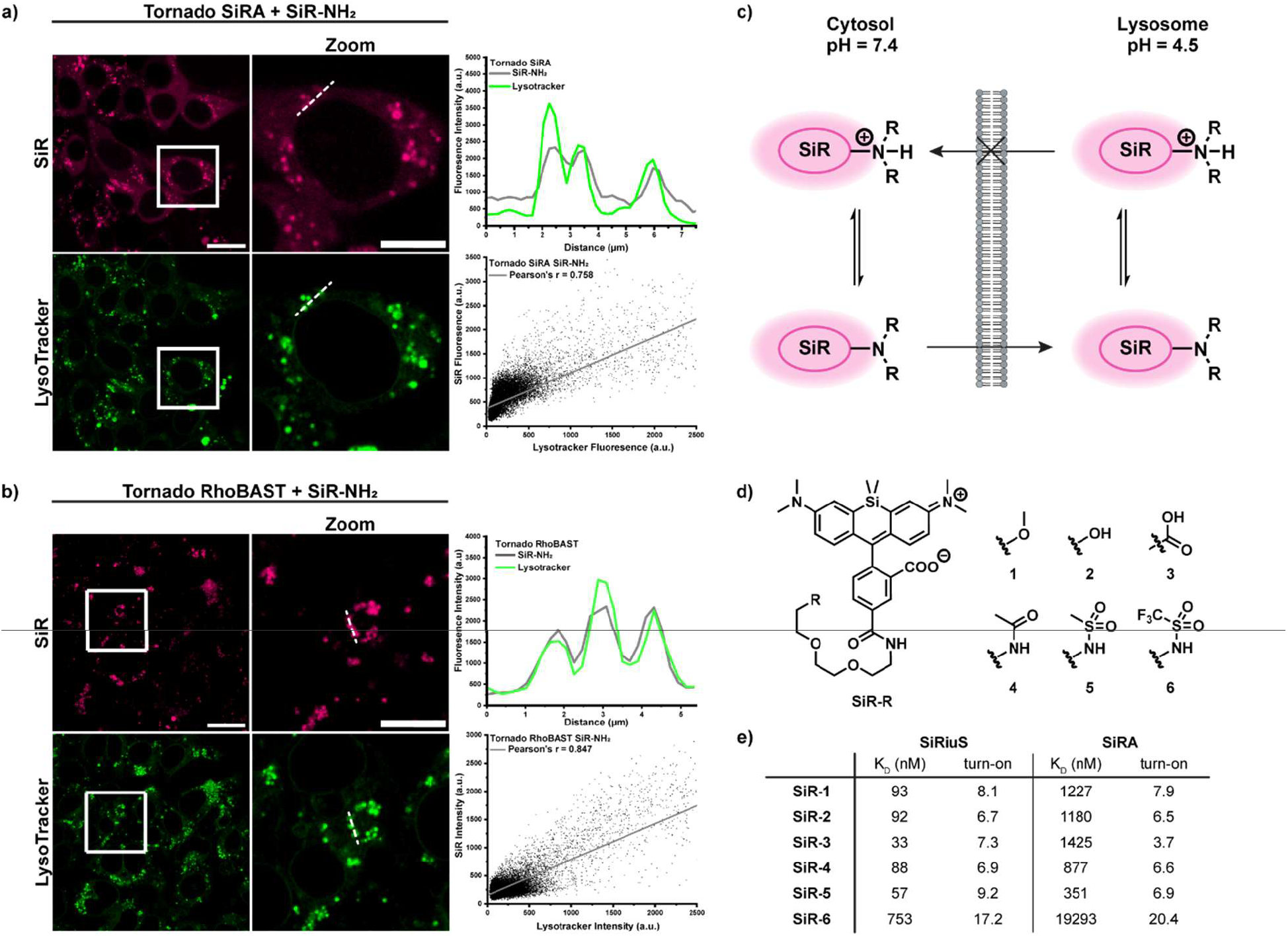
Lysosomal accumulation of SiR-NH_2_ in live cells. a) Colocalization experiment in live HEK293T cells expressing circular Tornado SiRA. Cells were incubated with SiR-NH_2_ (500 µM) and LysoTracker Green (60 µM). Intensity line plots and determination of Pearson’s coefficient show high correlation between the SiR and LysoTracker signal. b) Colocalization experiment in live HEK293T cells expressing circular Tornado-RhoBAST as negative aptamer control. Cells were incubated with SiR-NH_2_ (500 µM) and LysoTracker Green (60 µM). Intensity line plots and determination of Pearson’s coefficient show high correlation between the SiR and LysoTracker signal. c) Concept of ion trapping of basic terminal amino groups in the lysosome due to the acidic pH. d) Synthetic concept of designing NIR dyes suitable for FLAP imaging with SiRiuS and SiRA. e) *In vitro* properties of SiR-R derivates when binding to SiRiuS or SiRA.

During the development of SiRA, the (PEG)_2_-NH_2_ linker moiety (Figure 1a), initially employed to immobilize the target during SELEX, was found to be crucial for binding, reducing the K_D_ by sixfold to 430 nM compared to 2.6 µM for 5-carboxy-SiR (5C-SiR)^26^. For SiRiuS, the effect is even more pronounced, as 5C-SiR yielded a K_D_ of 6.25 µM, in stark contrast to 22 nM for SiR-NH_2_ (Figure S9, Tables S4 and S5). Notably, structurally closely related dyes, such as SiR-HaloTag7 ligand^36^ or the avidity probe SiR_2_^37^, have not shown similar accumulation problems, suggesting that the polar and basic nature of the terminal amino group might contribute to the formation of SiR-NH_2_ accumulations. Colocalization studies using LysoTracker Green in live HEK293T cells, expressing either circular SiRA (Figure 4a) or the orthogonal RhoBAST aptamer as a negative control (Figure 4b), demonstrated a strong correlation between the SiR-NH_2_ signal and green LysoTracker signal, with Pearson’s coefficients of r = 0.758 for SiRA-expressing cells and r = 0.847 for RhoBAST. This indicates the occurrence of lysosomal dye accumulation largely independent of the aptamer expression status.

We hypothesized that the observed lysosomal accumulation was due to ion trapping of SiR-NH_2_ in the acidic pH of lysosomes, as observed for other amines (Figure 4c)^38,39^. The uncharged spirocyclic SiR-NH_2_ can easily traverse the lysosomal membrane; however, the protonation of the basic terminal amino group in the acidic environment results in the formation of membrane-impermeable ammonium cations.

To mitigate lysosomal accumulation while preserving high binding affinity to SiRiuS and SiRA, we synthesized minimally modified SiR-R derivatives by substituting the terminal amino group with non-basic moieties, including, methoxy (SiR-1), hydroxy (SiR-2), carboxy (SiR-3), acetamide (SiR-4), methanesulfonamide (SiR-5) and trifluoromethanesulfonamide (SiR-6) groups (Figure 4d). Synthesis involved attaching the modified NH_2_-(PEG)_2_-R linker to 5C-SiR via amide coupling. Our goal was to engineer a reliable NIR aptamer:dye system suitable for live mammalian cells by varying the electronic and hydrogen-bond donor-acceptor properties, as well as the steric demand of the newly introduced terminal groups. The complete *in vitro* characterization of the SiR-R derivates can be found Figures S9-S11 and Tables S3-S5.

Probing the *in vitro* and *in cellulo* performance of the novel SiR-R dyes in conjunction with SiRiuS and SiRA identified SiR-1, SiR-4, and SiR-5 as the three most promising candidates. *In* vitro, these dyes exhibited high binding affinities in the sub-hundred nanomolar range and high turn-on, upon binding to SiRiuS and moderate binding affinity to SiRA (Figure 4e). Furthermore, these derivates produced a homogeneous cytosolic signal in live cells expressing either circular SiRiuS or SiRA (Figures 5c-e, S12, S14). SiR-6 showed drastically reduced binding affinity to either aptamer and accumulated in mitochondria (Figure S13). Conversely, SiR-2 and SiR-3 did bind to SiRiuS, however, SiR-2 generated lower fluorescence turn-on *in vitro* (Figure 4e and Table S5) and live-cell imaging of cells expressing circular SiRiuS with SiR-3 (100 nM) resulted in up to 7-fold lower mean fluorescence intensity, compared to the other derivates (Figure S15c).

**Figure 5.**
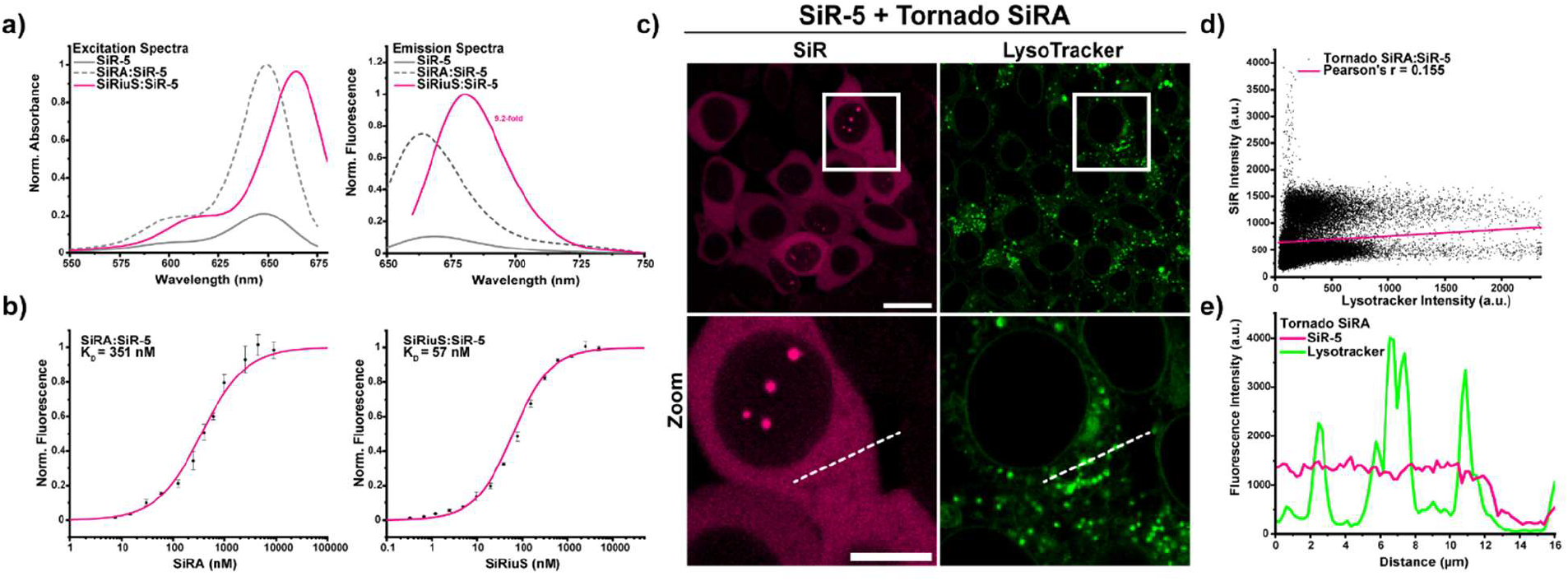
*In vitro* characteristics and *in cellulo* behavior of SiR-5. a) Excitation and emission spectra of SiR-5 (free dye), SiRA:SiR-5 and SiRiuS:SiR-5 complexes. b) Normalized K_D_ curves of SiR-5 binding to SiRA and SiRiuS. c) Colocalization of SiR-5 and LysoTracker Green in live HEK293T cells expressing circular SiRA in the cytosol. The bright spherical formations in the nucleus are previously reported Tornado punctae due to high concentration of circular RNA in the nucleus leading to phase separation of RNA^16,28^. d) Pearson correlation plot of the SiR-5 and LysoTracker signals shown in c). e) Line plots of SiR-5 and LysoTracker in c). Scale bars: 20 µm.

Further assessments centered on the spectral characteristics of the new SiR-R derivatives, revealing excitation in the NIR region at 648 nm and emission at 668 nm for all free dyes in buffered aqueous solutions (Figures 5a, S10, and Table S3). Upon binding to the SiRA aptamer, these dyes demonstrated similar excitation and emission properties as in the unbound state, however, when bound to SiRiuS, a bathochromic shift by 15 nm and 20 nm in excitation and emission maxima, respectively, is observed. A similar red shift in excitation and emission has previously been reported in the DFHBDI chromophore binding to the Spinach aptamer and could be attributed to long range electrostatic interactions and the negatively charged RNA backbone, stacking interactions with the nucleosides and H-bond formation^40^. Practically, this red shift enhances fluorescence turn-on at the emission maximum, while minimizing background fluorescence of the free dye. It is important to note that the standard laser box of the confocal microscope used for these experiments is equipped with a fixed 640 nm excitation laser, which is better suited for the unbound dye (excitation maximum at 648 nm) than for the SiRiuS-bound dye (664 nm). Consequently, the *in cellulo* fluorescence turn-on may be improved by utilizing a full-spectrum laser with adjustable excitation wavelengths, thereby capitalizing on the bathochromic shift observed in the SiRiuS system.

To identify the best-performing dye derivative, we conducted titration experiments in live HEK293T cells expressing circular SiRiuS incubated with varying dye concentrations from 50 nM to 1 µM, and the fluorescence signal to background ratio under each condition was determined. The highest signal to background ratio of SiR-4 was reached at a concentration of 500 nM, resulting in a 8.9-fold fluorescence turn-on, while SiR-1reached its highest turn-on of 9.4-fold at 250 nM. Notably, SiR-5 consistently produced an impressive 11.4-fold increase in fluorescence signal at a concentration of 100 nM in cells transfected with circular SiRiuS (Figure S14). Given that the sulfonamide derivative SiR-5 combined favorable *in vitro* characteristics, including high binding affinity to SiRiuS, high brightness, and red-shifted excitation/emission properties, alongside robust fluorescence turn-on at low concentrations in live cells, it was selected as the optimal candidate for the SiRiuS FLAP system.

Having established the SiRiuS:SiR-5 combination as the leading candidate, we aimed to benchmark its performance against its predecessor, SiRA. Consequently, we transfected live HEK293T cells with both circular SiRiuS and SiRA, evaluating the fluorescence turn-on following incubation of both samples with 100 nM SiR-5. Under optimized microscope settings, the SiRiuS:SiR-5 system yielded an impressive 14.6-fold increase in fluorescence, while SiRA exhibited a modest 2.7-fold fluorescence turn-on (Figures 6a,b and S15). To assess the applicability of SiRiuS:SiR-5, we tested it in different commonly used cell lines, namely HeLa, COS-7 and U2OS. Cells were transfected to express circular SiRiuS and were incubated with SiR-5 (100 nM) for 1 h. Similar to our studies in live HEK293T cells, transfected cells exhibited a uniform and bright fluorescence signal in the SiR-channel with little background (Figure S16). These findings confirm the enhanced performance and broad utility of the SiRiuS:SiR-5 system in live-cell imaging.

**Figure 6.**
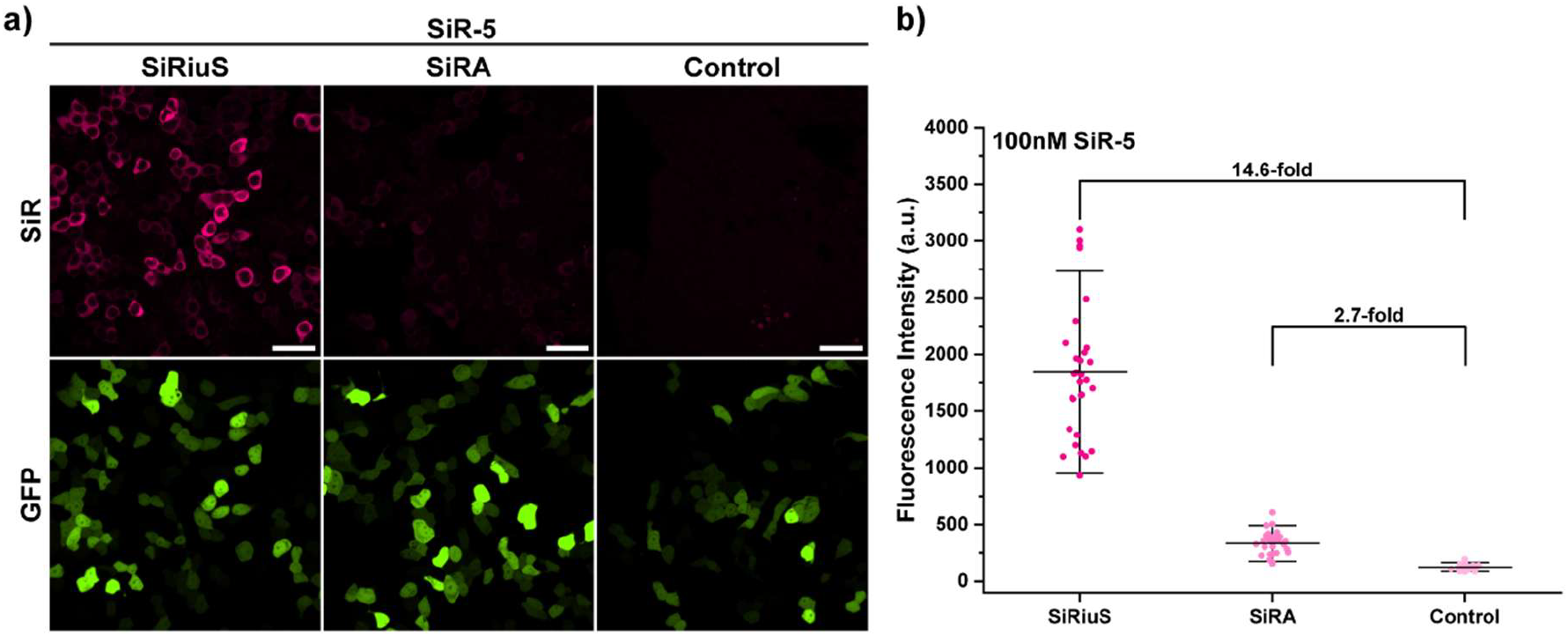
Improved performance of SiRiuS:SiR-5 in comparison to SiRA:SiR-5. a)Comparison of the performance of SiR-5 in live HEK293T cells expressing Tornado SiRiuS, Tornado SiRA or Tornado RhoBAST (negative control). b) Quantification of fluorescence intensity of positively transfected cells shown in a). Number of quantified cells per sample: *n*= 30. Scalebars: 20 µm. This experiment was performed in triplicates yielding similar results.

### Confocal imaging of mRNA in live mammalian cells

With the optimized SiRiuS:SiR-5 combination in hand, we set out to characterize the essential parameters for imaging in live mammalian cells using our systems. We could not observe dependence on monovalent cations (Figure S17a). Given that living cells are commonly imaged at 37°C within a live-cell chamber, we investigated the temperature dependence of the aptamer across a range of 25-75°C. While we observed a steady decline in fluorescence with increasing temperature, at 37°C, SiRiuS maintained 74% of its maximum fluorescence, consistent with results obtained for the original SiRA(Figure S17b). Furthermore, SiRiuS exhibited markedly reduced dependence on Mg^2+^ ions compared to SiRA; in fact, while the relative fluorescence of SiRA decreased to 70% at a 0.25 mM Mg^2+^ concentration^26^, SiRiuS remained above 90% relative fluorescence, even at 1 µM. In the physiological range for mammalian cells (0.25-1 mM Mg^2+ 16,26^), SiRiuS sustained over 95% of its maximum fluorescence (Figure S17c,d).Additionally, in a confocal setting, the SiR-5 dye was rapidly taken up by the cells and exhibited minimal bleaching during continuous imaging (Figure S18), demonstrating high photostability. For mRNA imaging with the SiRiuS aptamer, we cloned a 4x repeat cassette into the 3’ UTRs of various RNAs of interest (ROI). To benchmark achievable fluorescence turn-on levels for mRNA imaging, we imaged mEGFP mRNA tagged with 4, 8, and 16 repeats of the aptamer (Figure 7a). Selecting a gene that self-reports its expression, such as mEGFP, allows for optimal evaluation of aptamer performance with a non-circular RNA susceptible to degradation. We observed a linear increase in fluorescence from 4 to 8 repeats; however, minimal increases were noted when doubling from 8 to 16 repeats (Figure 7b). To minimize impact on RNA metabolism and cellular behavior while ensuring adequate fluorescence turn-on, we decided to utilize the 8x repeat cassette for tagging mRNAs occurring natively in the cell.

**Figure 7.**
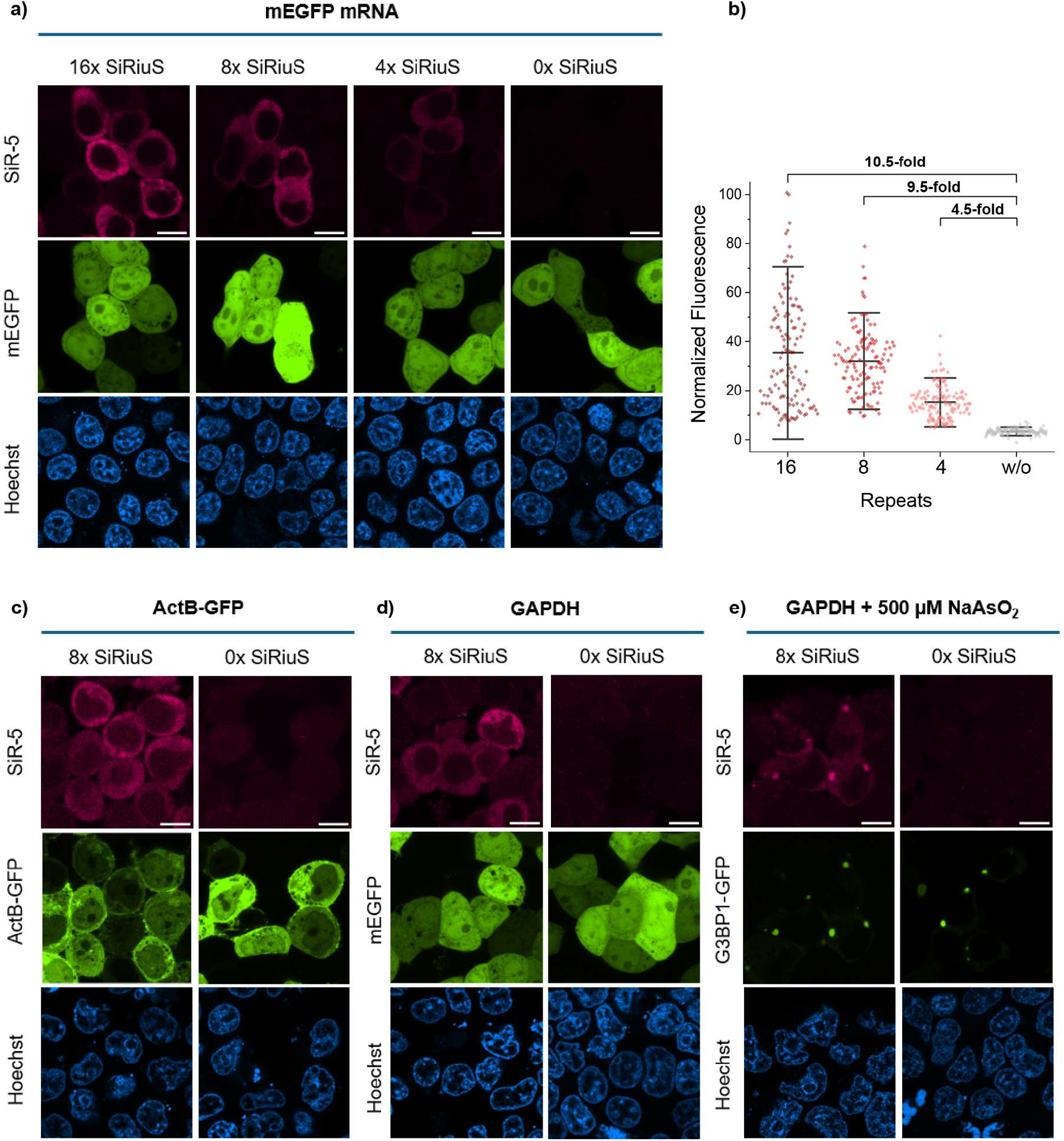
mRNA imaging in live HEK293T cells. a) Imaging mEGFP mRNA using different cassettes of the SiRiuS aptamer. HEK293T cells expressing mEGFP mRNA constructs with 0, 4, 8 and 16 SiRiuS repeats. b) Quantification of fluorescence in cells expressing mEGFP and turn-on compared to untagged constructs (*n*=50) and normalized to the highest measured signal. SiRiuS:SiR-5 was also used in tagging of well characterized mRNAs (c-e). A clear signal of the mRNAs is visible in each case, GAPDH accumulation when under stress colocalizes with stress granule marker G3BP1^41^ (e). Scale bars: 20 µm. Experiments were repeated three times with similar results.

To demonstrate the utility of SiRiuS for tagging mRNAs that are endogenous to the cell, we employed the 8x cassette to recombinantly tag GAPDH and actin-B mRNA (Figure 7c,d). These targets were selected due to their characterization and utilization as a standard in various studies^18,37,42,43^. The cassette was inserted into the 3′ UTR of GAPDH and actin-B-GFP mRNA on their respective plasmids as detailed in the methods section. GAPDH mRNA transcription was driven by the CMV promoter, while actin-B was expressed from its endogenous promoter. For GAPDH, cells were co-transfected with an mEGFP transfection control plasmid, while actin-B was tagged with GFP. Following a 24-hour incubation post-transfection, significant fluorescence was observed in cells transfected with either GAPDH or actin-B-GFP, demonstrating the capacity of SiRiuS to image mRNAs in mammalian cells. These mRNAs were found in uniform distribution throughout the cytoplasm consistent with previous reports using the same transcripts^18,37,42,43^.

To further showcase the utility of the SiRiuS system in a dynamic biological context, we aimed to demonstrate mRNA localization in subcellular structures. We focused on well-characterized stress granules, which are phase-separated condensates that form in response to stressors such as heat shock, oxidative stress, viral infection, UV light, or nutrient deficiencies^41,44^. In order to observe mRNA recruitment in stress granules, we decided to select the GAPDH 8x SiRiuS system due to GAPDH’s extensively studied behavior in stress response and its high expression level (Figure 7e)^45–47^. Cells were co-transfected with a plasmid encoding GFP tagged G3BP1, a stress granule-associated protein, and a plasmid encoding GAPDH 8x SiRiuS mRNA. Sodium arsenite (NaAsO_2_) was used as an oxidative stressor; cells were incubated with 500 µM NaAsO_2_ for 15 min prior to imaging to induce stress granule formation 24 h after transfection. We successfully observed the accumulation of GAPDH mRNA within the stress granules, colocalizing with G3BP1-GFP signals, while also retaining fluorescence in the cytosol, as described in the literature^48^. Although only a limited fraction of GAPDH mRNA accumulates within granules^47^, we were able to visualize these granules after 15 min(Figure 8). in a time-resolved manner. We performed time-lapse imaging of stress granule assembly post-stress induction, capturing dynamic changes in granule formation over time (Figure 8 a). Detailed temporal analysis revealed specific formation kinetics and structural changes in the granules, providing insight into the mRNA localization mechanism under stress conditions.

**Figure 8.**
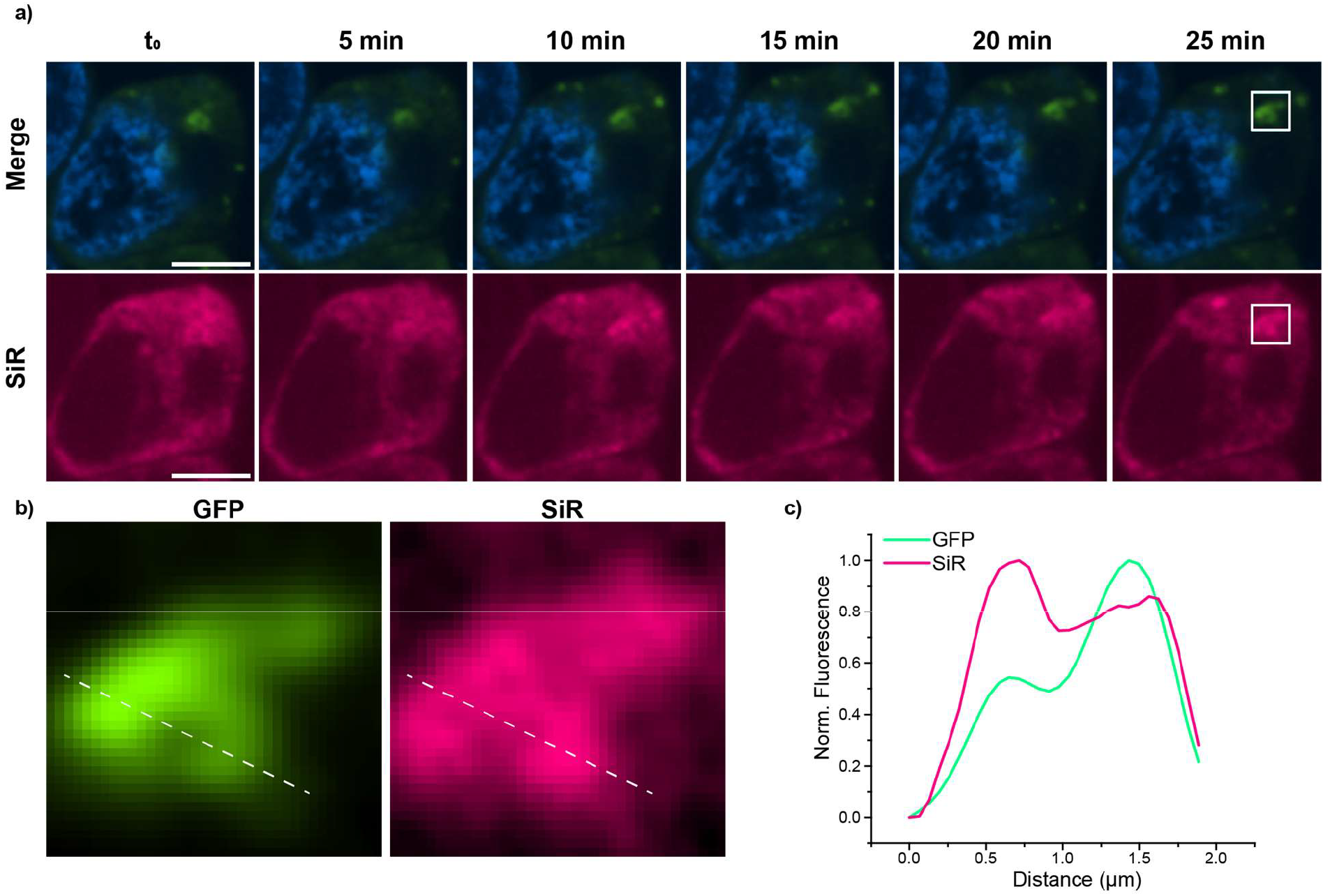
Time series of the formation of stress granules. a) Merge shows Hoechst and GFP channel. Larger G3BP1-GFP granules start forming after 10-15 min and GAPDH 8x SiRiuS signal could be observed after 15 min accumulating in the stress granules. Time stamps in min after NaAsO_2_ addition. Scale bars 5 µm. b) Zoom in (white boxes in a) of the colocalizing granules with G3BP1-GFP protein and GAPDH 8x SiRiuS mRNA. Scale bars 2 µm. c) Corresponding Intensity line graphs to cross section in b).

### Super-resolution imaging in mammalian cells using stimulated emission depletion (STED)

Next, we aimed to leverage the SiRiuS:SiR-5 pair for super-resolution NIR STED imaging, a capability that until now had only been demonstrated with SiRA in bacterial cells. The unique bathochromic shift associated with the SiRiuS:SiR-5 combination is anticipated to enhance the signal-to-background ratio in STED microscopy, particularly since the system employed is equipped with a fully adjustable laser. To optimize our imaging efforts, we focused on the above-described stress granule system that provides a localized target amidst the diffuse presence of mRNAs in the cytosol. We imaged live HEK293T cells expressing GAPDH mRNA directly 15 min after NaAsO^2^ treatment and successfully visualized stress granules at super-resolution using STED microscopy (Figure 9a,b). To assess whether SiRiuS is also compatible with fixed-cell imaging, we treated GAPDH 8x SiRiuS expressing cells with NaAsO_2_, fixed them with paraformaldehyde and imaged using SiR-5. Stress granules were likewise detected in fixed cells (Figure 9a.c) with a remarkable resolution of 43 nm.

**Figure 9.**
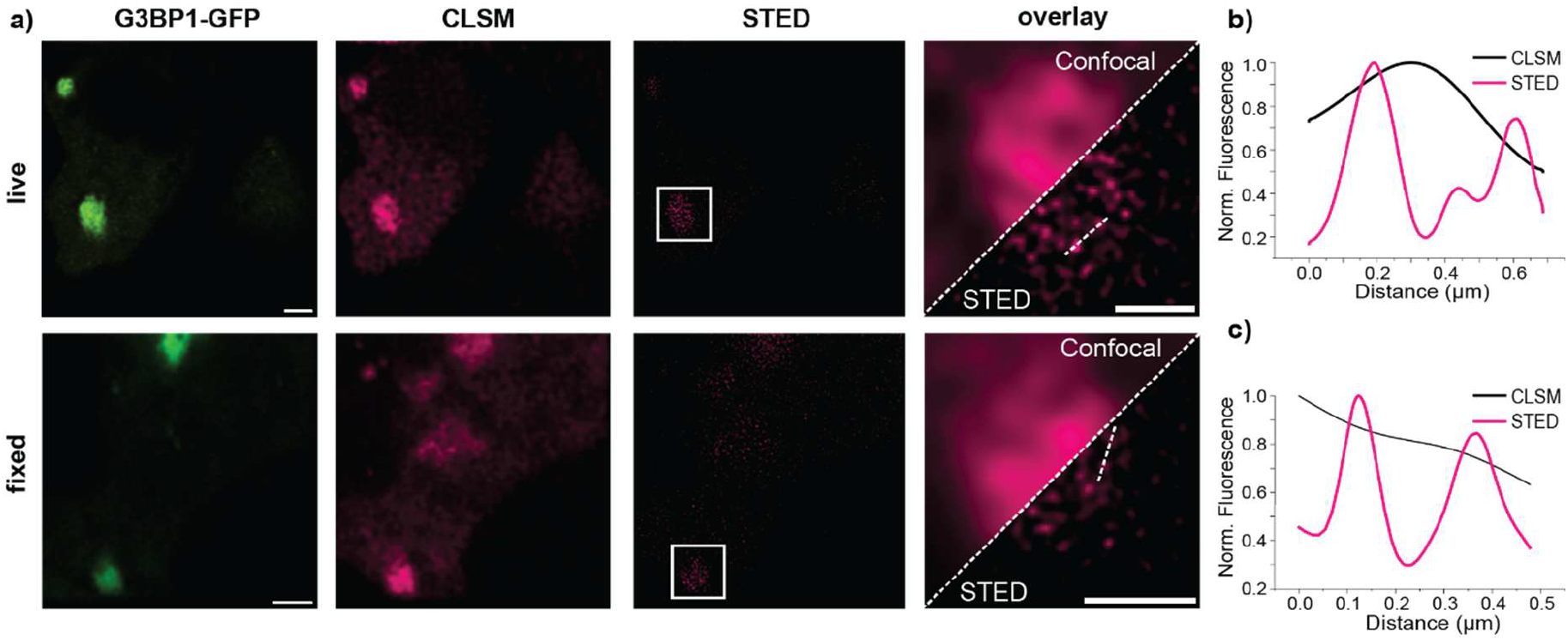
STED microscopy of GAPDH mRNA accumulation in phase-separated stress granules. a) Cells were transfected with the GAPDH 8xSiRiuS mRNA and G3BP1-GFP plasmids and stress was induced 15 min before imaging. For live cells, images were taken for no longer than 5 min after 15 min induction. Fixed cells were fixed at precisely 15 min and could subsequently be imaged indefinitely. Images show colocalization in confocal images between G3BP1 and SiR fluorescence. For fixed cells, 43 nm resolution was achieved compared to 311 nm for the respective confocal image, for live cells 94 nm compared to 419 nm for the respective confocal image. Resolution was calculated using the Image Decorrelation Analysis plugin for FIJI. Experiments were performed 3 times with similar results. Scale bars: 2 µm. b,c) Intensity line plot for cross section of b) live cells and c) fixed cells. Black line shows intensity of the cross section for confocal pictures, magenta for STED pictures c).

While the Okra and Pepper aptamers have been used in SIM microscopy, including the imaging of stress granules and other aggregates achieving a resolution of 155 nm^43,49^, the only other FLAP system demonstrated for STED imaging in mammalian cells is the RhoBAST:SpyRho system, also developed in our lab^50^. This FLAP operates at shorter wavelengths and is orthogonal to the SiRiuS system (Figure S19). To assess the potential for multicolor STED imaging using both systems in tandem, we tagged mAzurite mRNA (which was previously shown to also accumulate in stress granules after oxidative stress^37^) with eight repeats of RhoBAST and co-transfected the cells with GAPDH 8x SiRiuS and mAzurite RhoBAST plasmids. Cells expressing both plasmids were then imaged to evaluate the feasibility of dual-color STED imaging. Notably, we observed distinct and clearly resolved signals for both the SiRiuS and SpyRho systems (Figure 10), demonstrating the potential for multicolor applications in super-resolution imaging. Interestingly, we observed that GAPDH and mAzurite mRNA aggregates did not perfectly colocalize, constistent with observations reported by other researchers^16^. These results highlight the versatility of the SiRiuS system in combination with other FLAP systems, paving the way for future explorations of complex RNA dynamics within live mammalian cells.

**Figure 10.**
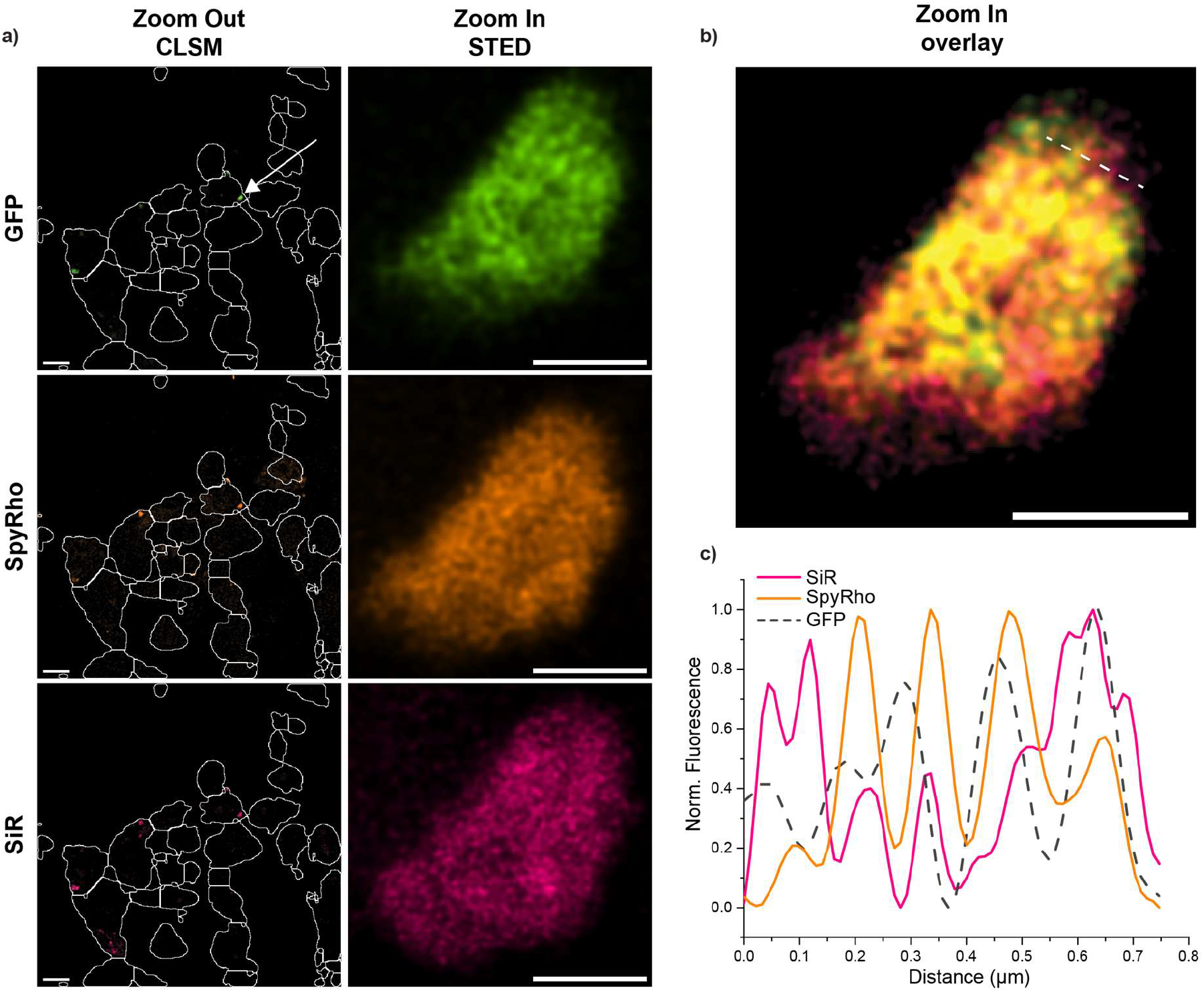
Dual-color STED microscopy of mRNA accumulation in phase separated stress granules. a) Cells were cotransfected with the GAPDH 8xSiRiuS mRNA plasmid, mAzurite 8xRhoBAST mRNA plasmid and G3BP1-GFP plasmid for colocalization. Oxidative Stress was induced 15 min before fixation. Images show stress granule formation for GFP, SpyRho and SIR fluorescence in multiple cells (left) and zoomed in STED image of the stress granule marked with the white arrow. Scale bar 20 µm for Zoom Out, 1 µm for Zoom-In pictures. Experiments were performed 3 times with similar results. b) Merge of STED images of GFP, SpyRho and SiR. Line,c) Line plots of SpyRho/SiR-5 STED images (orange/magenta) of cross-sections in a) compared with confocal profiles. Fluorescence on y axis was normalized to each curves cognate maximum.

## Conclusion and Outlook

In this work, we have demonstrated the exceptional capabilities of the SiRiuS system for mRNA imaging in both live and fixed mammalian cells within the near-infrared (NIR) region, utilizing confocal and STED microscopy techniques. By re-selecting an existing SELEX library with a focus on fluorescence turn-on, we successfully identified an aptamer that exhibits enhanced affinity for SiR and optimized turn-on characteristics specifically in mammalian cell environments.

Through careful structural elucidation, rational design, and strategic truncation of the aptamer, we developed a modular structure that supports stable repeat folding while achieving maximal fluorescence turn-on with as few as eight repeats. This innovation signifies a substantial advancement in aptamer engineering, as it minimizes the influence on RNA metabolism and cellular behavior during imaging.

Furthermore, we modified SiR-NH_2_, initially the highest-affinity dye for SiRA, to render it suitable for live imaging in mammalian cells. This alteration effectively mitigated ion trapping, which commonly occurs in lysosomal environments, while still capitalizing on the favorable effects of the linker on binding affinity. Remarkably, these modifications resulted in only a minimal decrease in binding affinity and fluorescence turn-on in live cells, while completely preventing accumulation problems.

In summary, we have developed an aptamer-dye system that boasts the highest excitation wavelength currently available within the FLAP toolbox. Additionally, it is the first system to facilitate imaging in the NIR spectrum and to be employed in STED microscopy applications in mammalian cells. We envision that the SiRiuS system will catalyze major advancements in RNA cell biology, particularly when used in conjunction with the RhoBAST:SpyRho system for multicolor STED imaging. This synergy has the potential to enable exploration into the organization of subcellular RNA structures as well as investigations into RNA interactions.

While our study primarily focused on mRNA imaging, the sensitivity and versatility of the SiRiuS system suggest that it could be effectively adapted for the visualization of non-coding RNAs, such as long non-coding RNAs (lncRNAs). Future research could thus expand the applicability of the SiRiuS system, enabling comprehensive insights into the functional roles of various RNA species and their interactions within the intricate landscape of cellular environments. Ultimately, this work lays the groundwork for innovative imaging strategies that will deepen our understanding of RNA dynamics and functionality in living systems.

## Supporting information

Experimental methods, Supplementary data, Supplementary tables

## ASSOCIATED CONTENT

### Supporting Information

The Supporting Information is available free of charge at:

Full experimental details, Supplementary Figures and Tables.

## AUTHOR INFORMATION

### NOTES

The authors declare no competing financial interest.

## ACKNOWLEDGEMENTS

This work was supported by the Deutsche Forschungsgemeinschaft (Grant No. Ja 794/11). We thank the Nikon imaging Center for access to their facilities and technical support with the microscopes. Additionally, we want to acknowledge Dr. Holger Lorentz and the ZMBH imaging facility for access to their STED microscope and help with image acquisition and quality assessment, and the ZMBH Flow Cytometry & FACS Core Facility (FFCF). We also thank Nils Materna for experimental assistance, Heiko Rudy for mass spectrometry and Tobias Timmermann for NMR measurements.

