## Supplementary material for "Co-evolution of a near-infrared aptamer:dye system for live-cell super-resolution RNA imaging": Experimental methods, Supplementary data, Supplementary tables

#### General

Reagents and solvents involved in the synthesis were purchased from commercial suppliers including Sigma-Aldrich, ABCR, Acros, TCI, Alfa Aesar and Carbolution or from the reprocessing plant of Heidelberg University. If not stated otherwise, all chemicals were used without further purification. Reactions requiring an inert atmosphere were carried out using standard Schlenk techniques under argon atmosphere.

Manual silica gel column chromatography was performed using technical grade silica gel (60 Å, 230-400 mesh, 40-63 µm) purchased from Sigma Aldrich. Automated normal phase column chromatography was performed IntelliFlash 310 by Varian and Puriflash XS520Plus by Interchim. Silica flash columns were purchased from Telos, Interchim and Büchi. All fluorophores were purified *via* reverse phase High Performance Liquid Chromatography (HPLC) prior to their usage in all *in vivo* and *in vitro* experiments. HPLC runs were performed on an AGILENT 1100 Series HPLC system equipped with a multi wavelength detector and a fraction collector using a Phenomenex Luna 5 µm C-18(2) 100 Å (250 x 21.2 mm) column. Purification runs were carried out at a constant flow rate of 5 mL/min using water and acetonitrile (each containing 0.1% of TFA as additive) as eluents. Product fractions were freeze-dried using a lyophilizer Alpha 2-4 LDplus by Christ.

All NMR spectra were recorded on the following NMR spectrometers: Varian Mercury Plus 300 (300 MHz) and Varian Mercury Plus 500 (500 MHz). If not stated differently, all spectra were recorded at room temperature (298 K). In $^{1}H$ and $^{13}C$ spectra, chemical shifts (δ) are listed in ppm using tetramethylsilane as an internal standard, while coupling constants (*J*) are listed in Hz. All recorded $^{1}H$ and $^{13}C$ spectra were referenced to the protio impurity or the $^{13}C$ signal of the deuterated solvent. $^{13}C$ measurements were recorded as APT spectra. Deuterated solvents were purchased from Euriso-Top.

High resolution mass spectra (HR-MS) were measured on a Bruker microTOF QII-ESI system using sodium formate as internal calibrant. The reported mass to charge ratios (m/z) refer to the isotopic peak with the highest intensity.

All general cloning procedures were performed with enzymes purchased from Thermo Fisher Scientific. *E. coli.* DH5α and BL21 Star (DE3) strains were purchased from Thermo Fisher Scientific. Bacterial cells were transformed with heat shock (30 sec at 42 °C) and grown with LB medium or plates (Carl Roth) with appropriate antibiotic. Fluorescence *in vitro* data was obtained on a FP-8500 Spectrofluorometer (Jasco). Excitation and emission filters were placed at ±5 nm of desired value. Absorbance spectra were recorded on Cary 100 Bio UV/Vis spectrometer (Varian) and a multimode microplate reader (TECAN Spark). PCR was performed using a T100 Thermal Cycler (BioRad). PCRs were performed using Q5 Hot Start® DNA Polymerase (NEB) or Taq polymerase produced in house. Oligonucleotides were purchased from IDT. Plasmids were obtained from Addgene. Agarose gels were stained with ethidium bromide and read with the Azure 300Q imager (Azure Biosystems) using its ethidium bromide setting. SHAPE gels and RNA purification gels were cast using 25% Rotiphorese® Sequencing Gel (Carl Roth) diluted to 20% and 10% respectively. SHAPE gel results were read on the Typhoon FLA 9500 using the

Phosphor imaging preset. Nucleic acid concentrations were measured with the NanoDrop One (Thermo Fisher Scientific). dNTPs were obtained from Jena Biosciences. 1-Methyl-7-nitroisatoic anhydride (1M7) was obtained from Tocris Bioscience. $^{32}$P-ATP was ordered from Hartmann Analytics.

#### *In vitro* transcription

*In vitro* transcription was performed in 500 µL reactions with 10 µg of DNA template (φ 6.5 T7 promotor at the 5'-end of the sequence, Table S1), T7 polymerase (1 µM) produced in house, 10 mM DTT, transcription buffer (40 mM Tris-HCl pH 8.1, 2 mM spermidine, 22 mM $MgCl_2$, 0.01%Triton X100) and 2 mM ATP\UTP\CTP\GTP. Reactions were incubated for 4 h at 37 °C and subsequently digested with 20U DNAse I for 30 min at 37 °C.

RNA was purified via gel electrophoresis on a denaturing 10% PAGE gel and excised using UV shadowing. RNA was eluted with the crush and soak method using 0.3 M NaOAc and subsequently precipitated with isopropanol (1 volume) and washed two times with 75% cold ethanol and dissolved in MilliQ® water. Concentrations were measured using the Nanodrop. To obtain accurate values, the aptamers were digested with NaOH into single nucleotides, and their concentrations were calculated based on their sequence and the extinction coefficients of the individual nucleotides.

#### FACS SELEX

For insertion into pET 28(+) plasmids, 1 µg of enriched DNA pools (7 and 14) from the SiRA SELEX (kindly provided by Dr. Regina Wirth) were PCR amplified in 10 cycles using Primer A and B with SalI and AatII site overhangs. Next, the PCR product was digested with SalI and AatII, and ligated to the pET28-tRNA$^{Lys}$ plasmid which had been digested with SalI and AatII.

For performing FACS on bacterial cells, BL21 Star (DE3) *E. coli* strain was transformed with the 200 ng of plasmid libraries L1 and L2, and the control plasmid pET-tRNA$^{Lys}$ using a standard electroporation procedure on a MicroPulser Electroporator (BioRad). After a 1 h rescue period in LB medium, bacteria were grown in 15 mL LB medium with 30 µg/mL kanamycin until an $OD_{600}$ of 0.4 was reached. Cells were grown for 3 h after addition of a final concentration of 1 mM IPTG. Cells were fixed with 1% of PFA for 10 min at room temperature. Cells were pelleted and resuspended in M9 medium. SiR-$NH_2$ was added to a final concentration of 1 µM for rounds 1-3 and 200 nM for round 3-L and incubated at 37 °C for 30 min before starting FACS.

The FACS experiments were performed on a FACS Aria III in the FFCF Facility at ZMBH, Heidelberg University. The quad sorting gate was placed in a way that it included <0.1% of the negative control (pET-tRNA$^{Lys}$) in the selection gate but was still as close as possible to the population in order to catch low fluorescence binders for round 1 and 2. For round 3 and 3-L only the top 1% or 0.5% of the positive cells were collected. For each library, $10^4$ events were collected with a flow rate of 3000 cells/s. Collected cells were washed with MilliQ® water once and used as templates for the next library. DNA from each round was additionally subjected to Illumina sequencing to analyze sequence frequencies throughout the rounds. Briefly, cDNAs derived from each round were attached with offset primers set 1 and index barcodes using offset and barcode PCR, respectively[1]. The resulting multiplexed libraries were sequenced on a NextSeq 550 platform with custom PhiX spike in controls. Data processing, read quality control, length filtering, pre-hairpin loop identification and abundance counting were performed following the previously reported pipeline[1].

#### SHAPE probing

Constructs of L2-8 for SHAPE probing were designed according to the original method[2]. Stable tetraloop structures in the 5’ and 3’-parts of the structure were selected to be probed in order to enhance stable folding (Table S5). Primers for reverse transcription reactions were radioactively labelled in a Polynucleotide Kinase (NEB) reaction according to the manufacturer’s instructions. Primers were designed to bind the 3’ end and middle of the L2-8 sequence to ensure single-nucleotide resolution throughout the entire sequence.

The RNA template for SHAPE was prepared by *in vitro* transcription. For adduct formation, 1-Methyl-7-nitroisatoic anhydride (1M7) was utilized[3]. The modification step was performed by adding 1/10 volume of 100 µM 1M7 to freshly folded RNA (8 pmol) and incubating at 25°C or 37°C respectively. RNA was recovered by ethanol precipitation.

For reverse transcription, 1 pmol of RNA was incubated with 0.2 µM radioactively labelled primer and 500 nM of each dNTP. For sanger sequencing ladder reactions, 1 mM of the respective ddNTP was added to the reaction. Annealing was done by heating to 75°C for 10 min and snap cooling on ice for 1 min. Following incubation, 1/10 v of 100 µM DTT was added together with 5x Superscript IV buffer and 100 U of Superscript IV (Thermo Fisher scientific). Reverse transcription (RT) was incubated at 55 °C for 30 min. To denature RNA in the sample, 1/5 V of 1 M NaOH was added and incubated at 90 °C for 5 min.

Reactions were analyzed on 60 cm 20% PAGE run for 4 h and visualized using a phosphoscreen and subsequent phosphorescence readout on a Typhoon FLA9500 instrument (GE Healthcare).

SHAPE reactivities were calculated using the following formula:

$$R_{SHAPE} = \frac{I_{1M7} - I_{ctrl}}{(I_{1M7} - I_{ctrl})_{max}},$$

with $R_{SHAPE}$ being SHAPE reactivity and I being the band intensity (quantified with FIJI software) of the respective lane.

#### *In vitro* characterization

For every measurement, aptamer was freshly folded by incubating the final volume of aptamer and water at 75 °C for 2 min, followed by cooling to 25 °C at a rate of 1 °C per 10 s. 6X Aptamer Selection Buffer with Tween (ASB-T, 120 mM HEPES, 750 mM KCl, 30 mM $MgCl_2$, 0.5% Tween20) was added to a final concentration of 1X.

Turn-on values for mutation studies were determined by incubating 5 µM of freshly folded aptamer and 5 nM of dye. Fluorescence was determined at the emission maximum of the aptamer:dye complex. Turn-on is calculated as the quotient of the fluorescence signal intensity of complex and dye only.

Equilibration dissociation coefficients ($K_D$ values) were determined upon measuring the change in fluorescence of each dye upon binding to the respective folded aptamer as a function of the aptamer concentration at a fixed dye concentration (10 nM) in 1X ASB buffer. $K_D$ values were obtained by fitting the measured intensities to the previously reported equation[4], using OriginPro (2015):

$$F = F_0 + (F_\infty - F_0)\frac{(K_D + P_0 + [Apt] - \sqrt{([Apt] - P_0)^2 + K_D \cdot (K_D + 2[Apt] + 2P_0)}}{2P_0},$$

with $F$ describing the measured fluorescence intensity, $F_0$ the fluorescence of the free dye, $F_\infty$ the fluorescence upon saturation, $P_0$ the initial dye concentration and *[Apt]* the concentration of the aptamer. The fluorescence intensity $F$ in the $K_D$ curves was normalized to $F_{norm}$: using the following formula

$$F_{norm} = \frac{(F - F_{min})}{(F_{max} - F_{min})},$$

with $F_{min}$ and $F_{max}$ representing the asymptotic minimal and maximal fluorescence intensities, respectively, as calculated by the above-mentioned fit.

Extinction coefficients $\varepsilon$ of free dyes were obtained by recording absorbance spectra of the respective dye at different concentrations (5 µM, 10 µM, 20 µM, 30 µM, 40 µM) and fitting the absorbance intensities at the absorbance maxima to Lambert-Beer's law. To obtain $\varepsilon$ values for aptamer:dye complexes, absorbance spectra of the respective dye (1 µM) were recorded in the presence of an access of aptamer (5 µM) in 1x ASB-T buffer after incubation at room temperature for 1 h.

#### Quantum yields

Absolute fluorescence quantum yields were measured on the Quantaurus-QY Absolute PL quantum yield spectrometer C11347 series by Hamamatsu, using a 1 mL cuvette at a wavelength of 648 nm. Quantum yields of free SiR-R dyes were determined at a concentration of 0.14 µM in ASB buffer. For the analysis of quantum yields of the aptamer:dye complexes, a 0.14 µM solution of each SiR-R dye was incubated with an access (>90% bound dye to aptamer, according to the respective $K_D$) of folded SiRA or SiRiuS aptamer.

#### $D_{50}$-Values

Absorbance spectra of each dye (20 nM) were recorded in different solvent compositions of dioxane and water (0%. 10%, 20%, 30%, 40%, 50%, 60%, 70%, 80%, 90%, 100% v/v). Each measurement of every composition was carried out in independent triplicates. The absorbance was normalized to the corresponding absorbance maxima of each dye and plotted against the dielectric constant of the solvent mixtures[5]. $D_{50}$ values were obtained by determining the dielectric constant of each dye at half-maximum absorbance.

#### Cloning of plasmids

Plasmids were cloned using procedures provided by the manufacturers. Tornado overexpression plasmid pAV-U6+27-Tornado-Broccoli[6] was acquired from addgene (#124360). Aptamers were inserted into the scaffold by restriction cloning using 5' NotI and 3' SacII sites. The plasmid encoding for mEGFP was acquired form addgene (#18696)[7]. SiRiuS synonymous repeats were subsequently inserted into two cassettes (Original and Syn 1 with AAA spacer; Syn 2 and Syn 3 with AAA spacer) using redundant

SalI and XhoI restriction sites for the 4x cassette. The 4x cassette was then cut out using XhoI and SalI, cutting the 4x plasmid with only SalI to yield 8 repeats and additionally inactivating the connecting restriction site. The same was done for 16 repeats. Cassettes were inserted into GAPDH and ActB-GFP plasmids (as described in Bühler *et al.*[8]). Cassettes were PCR amplified from mEGFP plasmid using overhang PCR to introduce necessary sites

#### Cell culture

Eukaryotic HEK293T (DSMZ, ACC 635), U2OS (DSMZ, ACC 785), HeLa (DSMZ, ACC 57), Cos7 (DSMZ, ACC 60) cells were cultured at 37°C under 5% $CO_2$ in Dulbecco's Modified Eagle's Medium (Thermo Fisher Scientific) supplemented with 10% FBS (Gibco) and 100 U/mL penicillin and 100 µg/mL of streptomycin. Cells were passaged when reaching ~75% confluency, using TrypLE Express (Gibco) for detachment, quenching after 1 min at 37°C with complete DMEM medium.

#### Confocal imaging of mammalian cells

For imaging, 8- or 18- glass coverslip well plates (ibidi) were coated with poly-D-lysine (PDL) and 20 x $10^5$ cells/cm$^2$ were seeded. After 24 h, cells were transfected with FuGENE®HD (Promega) according to the manufacturers' instructions. After 48 h, cells were washed with Leibovitz's L15 (Gibco) and imaged in L15 containing the desired concentration of SiR-R and Hoechst 33258. Confocal images were taken in the Nikon Imaging Center Heidelberg using the Nikon A1R confocal microscope (Objectives: Nikon N Apo 60x NA 1,4 deltas OI (WD 0,14mm, FOV 0.21 x 0.21 mm) / Nikon Plan Fluor 40x NA 1.3 OI (WD 0.2 mm, FOV 0.32 x 0.32 mm)). Laser power values were provided by the Imaging Facility.

Time trace experiments were conducted on the Nikon Ti2-Wi spinning disk microscope in the Nikon Imaging Center Heidelberg using the Nikon Plan Apo λD 100 x Oil NA 1.45 (WD 0.13 mm) objective. Transfected cells were placed on the microscope and FOVs selected. To induce stress, 1 µL of 50 mM $NaAsO_2$ was added to cells in 99 µL of L15 and measurement was started immediately, marking $t_0$. Images were taken in 30 s intervals.

Excitation lasers for SiRiuS:SiR-5 imaging were set to the fixed 640 nm excitation and detection using a 700/75 filter (wavelength [nm]/bandwidth [nm])

Images were handled with the FIJI software v1.54m. Fluorescent values were quantified using the Time Series Analyzer v3 plugin. Background subtraction was performed by measuring cell-free room on images in at least 3 points and subtracting it from the image.

#### STED microscopy

For STED, a Leica TCS SP8 STED 3X microscope with Falcon FLIM was used with Leica Application Suite X (LASX) software. It is equipped with a 470-670 nm pulsed white light laser for excitations and 592/660/775 nm STED lasers. The GFP channel was excited using 3% 488 nm laser, SpyRho with 20% 561 nm laser and SiR-5 using 50% of 631 nm laser. For GFP and TMR, 100% STED laser (1.5 W both 592 nm and 660 nm STED lasers respectively) were used, for SiR-5 80% of >1.5 W 775 nm laser was used. Images were deconvolved using Huygens Professional 25.04 software.

For the STED imaging procedure, we prepared samples similarly to those used for confocal imaging of stress granules, fixing the cells 15 min post-induction. The fixation involved 4% paraformaldehyde (PFA) for 10 min, followed by quenching with PBS containing 100 mM glycine for 20 min, and two washes with this solution. Following fixation, we added 100 nM SiR-5 in L15 medium and proceeded with imaging.

### Chemical synthesis

#### Synthetic overview of SiR-R

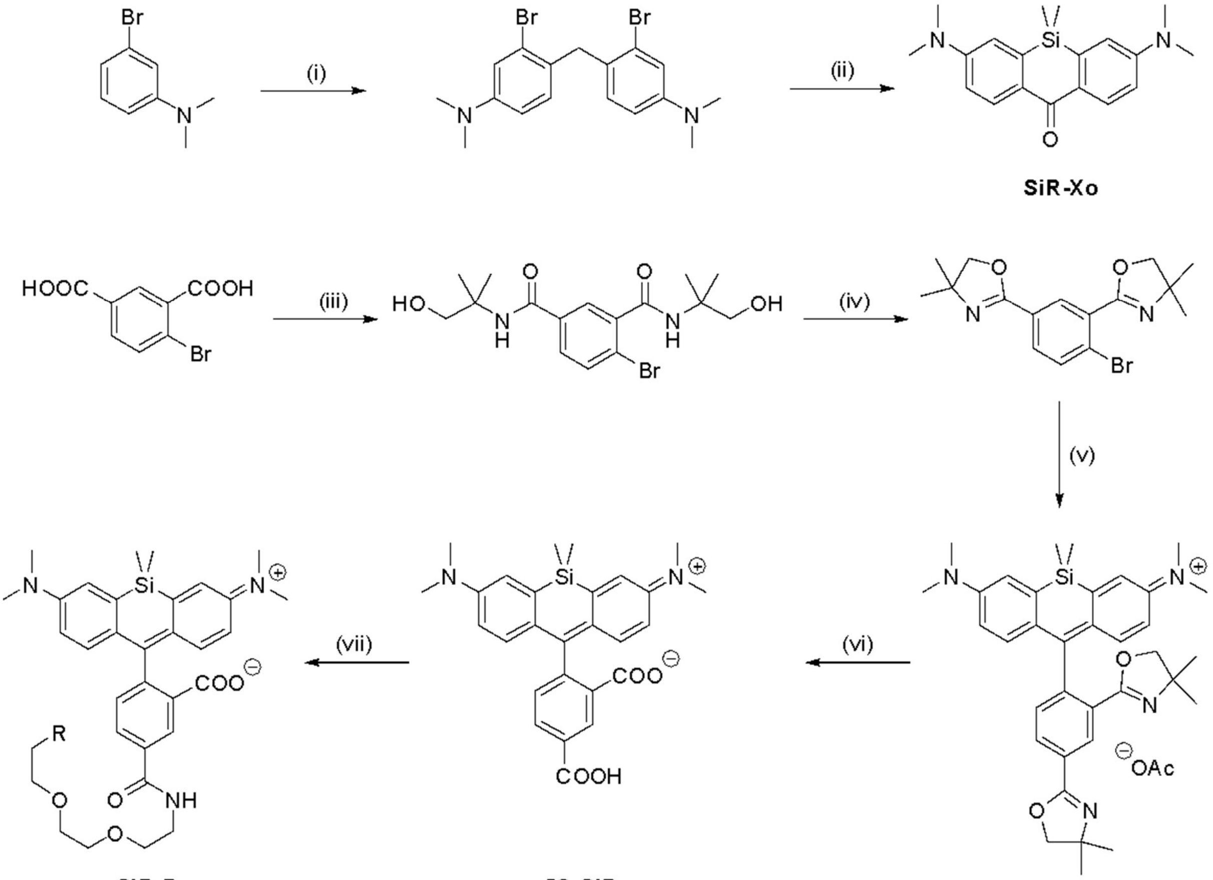

Scheme 1: Synthesis of SiR-R. (i) $CH_2O$, AcOH, 60 °C; (ii) 1) *sec*-BuLi, -78 °C, THF, 2) $Me_2SiCl_2$, -78 °C, THF, 3) $KMnO_4$, -15 °C, acetone; (iii) 1) $SOCl_2$, cat. DMF, reflux, 2) 2-amino-2-methylpropanol, DIPEA. DCM; (iv) $SOCl_2$; (v) 1) tert-BuLi, -78 °C, THF, 2) SiR-Xo, -78 °C, THF 3) AcOH; (vi) 6N HCl, 80 °C; (vii) 1) PyBOP, DIPEA, DMF, 2) $NH_2$-$(PEG)_2$-R, 0 °C, DMF.

#### General Procedures

General Procedure 1 (GP1): SiR-R

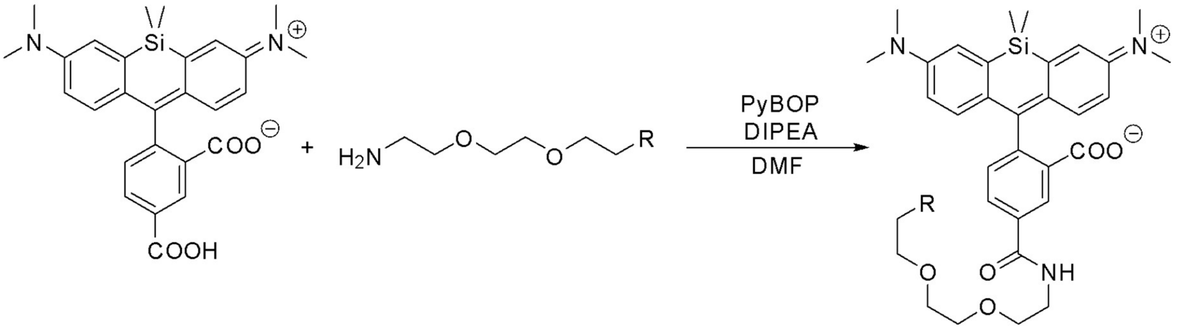

**5C-SiR** **$NH_2$-(PEG)$_2$-R** **SiR-R**

**5C-SiR** (known compound, prepared according to Wirth et al.[9]) (1.00 eq) and PyBOP (1.20 eq) were dissolved in dry DMF (500 µL) and DIPEA (2.40 eq) was added. The solution was placed in a thermo shaker at room temperature at 800 rpm for 2 h. In a flame dried Schlenk tube, the corresponding $NH_2$-(PEG)$_2$-R linker (1.20 eq) was dissolved in dry DMF (500 µL) and cooled to 0 °C. The solution containing the activated 5-Carboxy-SiR was added dropwise and the solution was stirred at room temperature for 20 h. The solvent was removed *in vacuo*, followed by purification by silica column chromatography and subsequent purification by reverse phase HPLC.

#### Syntheses and characterization

**4,4'-methylenebis(3-bromo-N,N-dimethylaniline)**

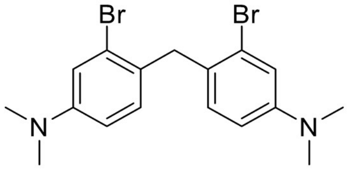

The title compound was prepared according to Wirth *et al.*[9]
3-bromo-*N,N*-dimethylanilie (3.0 g, 14.99 mmol, 2.14 mL, 2.00 eq) was dissolved in in glacial acetic acid (30 mL) and the solution was dropwise added to an aqueous formaldehyde solution (37%, 0.6 mL, 1.00 eq) at room temperature and the reaction mixture was stirred at 60 °C for 1 h. The mixture was cooled to room temperature and acetic acid was removed *in vacuo*. Saturated $NaHCO_3$ solution (70 mL) was slowly added to the residue and the solution was extracted with DCM (5x 30 mL). The combined organic phases were washed with water (3x 20 mL) and brine (1x 30 mL) and dried over $MgSO_4$. The solvent was removed *in vacuo* and the product was subsequently purified by silica gel column chromatography (cyclohexane:EtOAc, 50:1). 4,4'-methylenebis(3-bromo-*N,N*-dimethylaniline was obtained as a colorless solid (2.1 g, 5.09 mmol, 68%).

The recorded analytical data were in accordance with literature[9].

**$^{1}$H NMR (300 MHz, CDCl$_3$):** δ (ppm) = 6.95 (d, *J* = 2.7 Hz, 2H), 6.86 (d, *J* = 8.6 Hz, 2H), 6.59 (dd, *J* = 8.6 Hz,2.7 Hz, 2H), 4.01 (s, 2H), 2.92 (s, 12H).

**MS** (HR-ESI, pos): meas. m/z = 432.9868, calc. m/z = 432.9885 for $C_{17}H_{20}Br_2N_2Na$ $[M+Na]^+$.

**3,7-bis(dimethylamino)-5,5-dimethyldibenzo[b,e]silin-10(5H)-one**

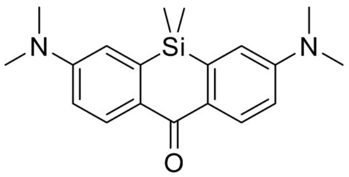

The title compound was prepared according to Wirth *et al.*[9]

In a flame-dried Schlenk flask, 4,4'-methylenebis(3-bromo-*N,N*-dimethylaniline (1.00 g, 2.43 mmol, 1.00 eq) were dissolved in dry and degassed THF (100 mL). The solution was cooled to -78 °C and *sec*-BuLi (1.4 M in *n*-hexane, 5.20 mL, 7.28 mmol, 3.00 eq) was added dropwise over 20 min. The orange solution was stirred for 2 h at -78 °C and $SiMe_2Cl_2$ (4.08 mL, 4.37 mmol, 1.80 eq) was slowly added over 15 min. The reaction mixture was slowly brought to room temperature and stirred for 18 h. Aqueous HCl (1 M, 10 mL) was added to the reaction mixture and THF was subsequently removed *in vacuo*. The resulting aqueous solution was neutralized by addition of conc. $NaHCO_3$ (5 mL) solution, and the solution was extracted with EtOAc (5x 20 mL). The combined organic phases were washed with water and brine and were dried over $MgSO_4$. The resulting solution was transferred in a flame-dried Schlenk flask, and the solvent was removed *in vacuo*. Due to its high sensitivity towards oxygen, the resulting crude was immediately used without further purification. The residue was dissolved in dry and degassed acetone (15 mL) and the solution was cooled to 0 °C. $KMnO_4$ (994 mg, 8.49 mmol, 3.50 eq) was added in small portions and the suspension was stirred for 6 h at 0 °C, followed by filtration through a plug of silica and rinsing with $CHCl_3$ (800 mL). The solvent was removed *in vacuo* and the product was subsequently purified by silica gel column chromatography (cyclohexane:$CHCl_3$:EtOAc 5:1:1). 3,7-bis(dimethylamino)-5,5-dimethyldibenzo[b,e]silin-10(5H)one was obtained as a colorless solid (350 mg, 1.08 mmol, 44%).

The recorded analytical data were in accordance with literature[9].

**$^{1}$H NMR (300 MHz, CDCl$_3$):** δ (ppm) = 8.40 (d. *J* = 9.0 Hz, 2H), 6.84 (dd, *J* = 9.0 Hz, 2.8 Hz, 2H), 6.79 (d, *J* = 2.8 Hz, 2H), 3.09 (s, 12H), 0.49 (s, 6H)

**MS** (HR-ESI, pos): meas. m/z = 325.1734, calc. m/z = 325.1731 for $C_{19}H_{25}N_2OSi$ $[M+H]^+$.

**4-bromo-*N*$^{1}$,*N*$^{3}$-bis(1-hydroxy-2-methylpropan-2-yl)isophthalamide**

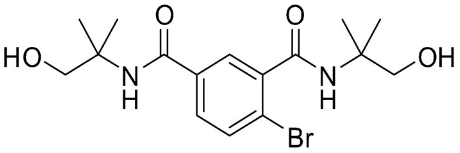

The title compound was prepared according to Wirth *et al.*[9]

4-Bromoisophthalic acid (5.00 g, 20.4 mmol, 1.00 eq) was suspended in freshly distilled $SOCl_2$ (30 mL)

and one drop of DMF was added as catalyst. The reaction mixture was refluxed for 4 h. After cooling to room temperature, $SOCl_2$ was removed *in vacuo*. The resulting residue was subsequently dissolved in dry and degassed DCM (60 mL) and was slowly added to a stirred solution of 2-amino-2-methylpropan-1-ol (5.00 g, 56.1 mmol, 2.75 eq) and DIPEA (10 mL) in dry DCM (60 mL) over 20 min. The reaction mixture was stirred for 18 h at room temperature, followed by the addition of saturated aqueous $NaHCO_3$ solution (60 mL) to neutralize the solution to pH 7. The mixture was extracted with EtOAc (5x 40 mL) and the combined organic phases were washed with water (80 mL) and brine (80 mL), dried over $MgSO_4$, filtered, and the solvent was removed *in vacuo*. The obtained white solid was used without further purification (4.25 g, 11.0 mmol, 54%).
The recorded analytical data were in accordance with literature[9].

**$^1$H NMR (300 MHz, DMSO-*d6*):** δ (ppm) = 7.84 (br s, 1H), 7.77 (d, *J* = 2.1 Hz, 1H), 7.74–7.64 (m, 3H), 4.86 (t, *J* = 6.0 Hz, 1H), 4.80 (t, *J* = 6.0 Hz, 1H), 3.51 (d, *J* = 3.2 Hz, 2H), 3.49 (d, *J* = 3.2 Hz, 2H), 1.31 (s, 6H), 1.30 (s, 6H).

**MS** (HR-ESI, pos): meas. m/z = 409.0717, calc. m/z = 409.0733 for $C_{16}H_{23}BrN_2NaO_4$ $[M+Na]^+$.

**2,2'-(4-bromo-1,3-phenylene)bis(4,4-dimethyl-4,5-dihydrooxazole)**

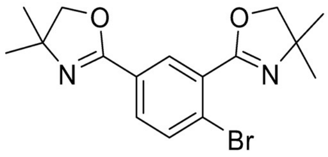

The title compound was prepared according to Wirth *et al.*[9]
In a flame-dried Schlenk flask, 4-bromo-*N[1],N[3]*-bis(1-hydroxy-2-methylpropan-2-yl)isophthalamide (1.00 g, 2.58 mmol, 1.00 eq) was dissolved in freshly distilled $SOCl_2$ (5.0 mL, 69.7 mmol, 27.0 eq) and stirred at room temperature for 2 h. $SOCl_2$ was removed *in vacuo* and the formed residue was dissolved in saturated aqueous $NaHCO_3$ (50 mL). The solution was extracted with EtOAc (5x 20 mL) and the combined organic phases were washed with water (50 mL) and brine (50 mL), dried over $MgSO_4$, filtered and the solvent was removed *in vacuo*. The product was subsequently purified by silica gel column chromatography (EtOAc, 100%). 2,2'-(4-bromo-1,3-phenylene)bis(4,4-dimethyl-4,5-dihydrooxazole) was obtained as a colorless oil (745.0 mg, 2.21 mmol, 82%).
The recorded analytical data were in accordance with literature[9].

**$^1$H NMR (300 MHz, $CDCl_3$):** δ (ppm) = 8.19 (d, *J* = 2.1 Hz, 1H), 7.80 (dd, *J* = 8.4 Hz, 2.1 Hz, 1H), 7.64 (d, *J* = 8.4 Hz, 1H), 4.11 (s, 2H), 4.09 (s, 2H), 1.40 (s, 6H), 1.36 (s, 6H).

**MS** (HR-ESI, pos): meas. m/z = 351.0703, calc. m/z = 351.0703 for $C_{16}H_{20}BrN_2O_2$ $[M+H]^+$.

**5-Carboxy-2-(7-(dimethylamino)-3-(dimethyliminio)-5,5-dimethyl-3,5-dihydrodibenzo[b,e]silin-10-yl)benzoate (5C-SiR)**

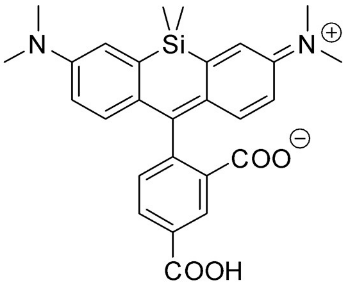

**5C-SiR**

The title compound was prepared according to Wirth *et al.*[9]

In a flame-dried Schlenk tube, 2,2'-(4-bromo-1,3-phenylene)bis(4,4-dimethyl-4,5-dihydrooxazole) (246.6 mg, 702.0 µmol, 1.70 eq) was dissolved in dry and degassed THF (8 mL) and the solution was cooled to -78 °C. *Tert*-BuLi (1.70 M in *n*-hexane, 461.5 µL. 784.6 µmol, 1.90 eq) was added dropwise over 20 min and the red solution was stirred for 2 h at -78 °C. In a second flame-dried Schlenk tube, 3,7-bis(dimethylamino)-5,5-dimethyldibenzo[b,e]silin-10(5H)one (134.0 mg, 412.9 µmol, 1.00 eq) was dissolved in dry and degassed THF (15 mL) and the resulting yellow solution was slowly added to the reaction mixture at -78 °C. The solution was brought to room temperature and stirred for 2 h. The reaction mixture was cooled to 0 °C and conc. acetic acid (2 mL) were added and the resulting blue solution was stirred at 0 °C for 45 min. The solvent was removed *in vacuo* and the blue residue was lyophilized to dryness. The crude was used for the next reaction step without further purification. The blue solid was dissolved in 6 M HCl (20 mL) and the yellow solution was stirred at 80 °C for 18 h. The reaction mixture was cooled to room temperature and the pH was adjusted to 2 through the addition of saturated aqueous $NaHCO_3$ solution. The mixture was extracted with DCM (5x 30 mL) and the combined organic phases were washed with HCl (0.1 M, 3x 20 mL) and brine (50 mL) and dried over $MgSO_4$. The solvent was removed *in vacuo* and the crude product was subsequently purified by silica gel column chromatography (DCM:MeOH 10:1). **5C-SiR** was obtained as a blue solid (153.0 mg, 323.7 µmol, 78%).

The recorded analytical data were in accordance with literature[9].

**$^1$H NMR (500 MHz, MeOH-d$_4$):** δ (ppm) = 8.35 (dd, *J* = 7.9, 1.7 Hz, 1H), 7.40 (d, *J* = 7.9 Hz, 1H), 7.30 (d, *J* = 2.8 Hz, 2H), 6.93 (d, *J* = 9.4 Hz, 2H), 6.74 (dd, *J* = 9.5, 2.9 Hz, 2H), 3.26 (s, 13H), 0.64 (s, 3H), 0.58 (s,3H).

**MS** (HR-ESI, pos): meas. m/z = 473.1884, calc. m/z = 473.1891 for $C_{27}H_{29}N_2O_4Si$ $[M+H]^+$.

***N*-(2-(2-(2-aminoethoxy)ethoxy)ethyl)acetamide (NH$_2$-(PEG)$_2$-NHAc)**

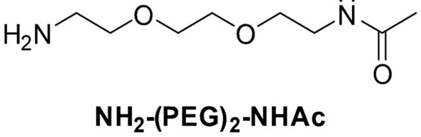

**NH$_2$-(PEG)$_2$-NHAc**

In a flame-dried Schlenk flask under argon atmosphere, 2,2'-(ethylenedioxy)bisethylenediamine (1.52 g, 10.2 mmol, 1.00 eq) was dissolved in anhydrous DCM (50 mL) and the solution was cooled to 0°C. A solution of acetic anhydride (208 mg, 193 µL, 2.04 mmol, 0.2 eq) in dry DCM (20 mL) was added dropwise over 15 min at 0 °C. The mixture was brought to room temperature and stirred for 8 h. The solvent was removed *in vacuo*, and the product was subsequently purified by silica gel column chromatography (3:1 EtOH:DCM + 5% $NH_4OH$ (aq)). *N*-(2-(2-(2-aminoethoxy)ethoxy)ethyl)acetamide (**$NH_2$-(PEG)$_2$-NHAc**) (227 mg, 1.19 mmol, 58%) was obtained as a colorless oil.

**$^1$H NMR (500 MHz, $CDCl_3$):** δ (ppm) = 6.59 (s, 1H), 3.63 (s, 4H), 3.57 (td, *J* = 5.1, 1.4 Hz, 4H), 3.47 – 3.42 (m, 2H), 2.92 (t, *J* = 5.1 Hz, 2H), 2.44 (s, 2H), 1.99 (s, 3H).

**$^{13}$C NMR (126 MHz, $CDCl_3$)** δ (ppm) = 170.6, 72.3, 70.4, 70.2, 70.1, 41.5, 39.5, 23.3.

**MS** (HR-ESI, pos): meas. m/z = 191.1394, calc. m/z = 191.1391 for $C_8H_{19}N_2O_3$ $[M+H]^+$.

##### *N*-(2-(2-(2-aminoethoxy)ethoxy)ethyl)methanesulfonamide ($NH_2$-(PEG)$_2$-$NHSO_2Me$)

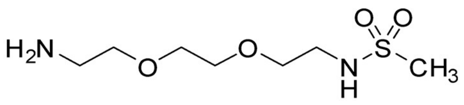

**$NH_2$-(PEG)$_2$-$NHSO_2Me$**

In a microwave tube under argon atmosphere, 2,2'-(ethylenedioxy)bisethylenediamine (150 mg, 1.01 mmol, 1.00 eq) was dissolved in dry DCM (3 mL) and the solution was cooled to -70 °C. A solution of methanesulfonic anhydride (88.15 mg, 506.05 µmol, 0.5 eq) in dry DCM (2 mL) was added dropwise over 15 min at -70 °C. The mixture was brought to room temperature and stirred for 8 h. The solvent was removed *in vacuo*, and the product was subsequently purified by silica gel column chromatography (3:1, EtOH:DCM + 5% $NH_4OH$ (aq)). *N*-(2-(2-(2-aminoethoxy)ethoxy)ethyl)methanesulfonamide (**$NH_2$-(PEG)$_2$-$NHSO_2Me$)** (88.0 mg, 3889 µmol, 77%) was obtained as a colorless oil.

**$^1$H NMR (500 MHz, $CDCl_3$)** δ (ppm) = 3.68 – 3.60 (m, 6H), 3.54 (t, *J* = 5.2 Hz, 2H), 3.33 – 3.28 (m, 2H), 2.88 (t, *J* = 5.1 Hz, 2H).

**$^{13}$C NMR (126 MHz, $CDCl_3$)** δ (ppm) = 73.3, 70.6, 70.1, 70.1, 43.2, 41.7, 40.5.

**MS** (HR-ESI, pos): meas. m/z = 227.1064; calc m/z = 227.1060 for $C_7H_{18}N_2O_4S$ $[M+H]^+$.

##### *N*-(2-(2-(2-aminoethoxy)ethoxy)ethyl)-1,1,1-trifluoromethanesulfonamide ($NH_2$-(PEG)$_2$-$NHSO_2CF_3$)

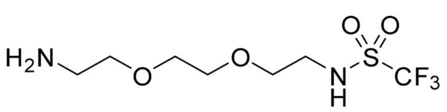

**$NH_2$-(PEG)$_2$-$NHSO_2CF_3$**

In a microwave tube under argon atmosphere, 2,2'-(ethylenedioxy)bisethylenediamine (200 mg, 1.35 mmol, 1.00 eq) was dissolved in dry DCM (5 mL) and the solution was cooled to -70 °C.

Trifluoromethanesulfonic anhydride (114.2 mg, 114.2 µL, 0.5eq) was added dropwise over 15 min at -70 °C and the solution was stirred for 1 h at -70 °C. The mixture was brought to room temperature and stirred for 3 h, resulting in a milky suspension. The solvent was removed *in vacuo*, and the product was subsequently purified by silica gel column chromatography (3:1, EtOH:DCM + 5% $NH_4OH$ (aq)). **$NH_2$-(PEG)$_2$-$NHSO_2CF_3$** was obtained as a colorless oil (164 mg, 585 µmol, 87 %).

**$^1$H NMR (500 MHz, MeOH-d4):** δ (ppm) = 3.73 – 3.69 (m, 2H), 3.66 (tdd, *J* = 7.3, 4.1, 2.0 Hz, 4H), 3.54 (t, *J* = 5.6 Hz, 2H), 3.26 (td, *J* = 5.6, 1.1 Hz, 2H), 3.04 – 3.00 (m, 2H).

**$^{13}$C NMR (126 MHz, MeOH-d4):** δ (ppm) = 123.0 (q, *J* = 326.3 Hz), 73.1, 71.3, 71.1, 69.5, 46.0, 40.9.

**$^{19}$F NMR (282 MHz, MeOH-d4):** δ (ppm)= -78.84 (s).

**MS** (HR-ESI, pos): meas. m/z = 303.062, calc. m/z = 303.0597 for $C_7H_{15}F_3N_2O_4SNa$ $[M+Na]^+$.

**2-(7-(dimethylamino)-3-(dimethyliminio)-5,5-dimethyl-3,5-dihydrodibenzo[b,e]silin-10-yl)-5-((2-(2-(2-methoxyethoxy)ethoxy)ethyl)carbamoyl)benzoate (SiR-1)**

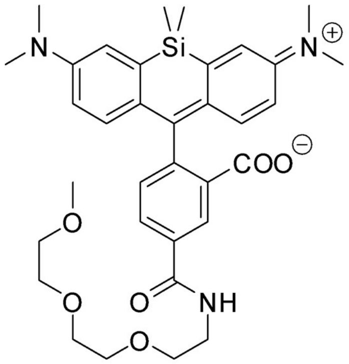

**SiR-1**

**SiR-1** was prepared according to **GP1.** The crude product was purified via normal phase silica column chromatography ($CHCl_3$:MeOH 50:1 to 20:1) and reverse phase HPLC (0.1 % TFA, $H_2O$/MeCN). **SiR-1** was obtained as a blue solid (7.05 mg, 11.4 µmol, 54%).

**$^1$H NMR (500 MHz, MeOH-d4):** δ (ppm) = 8.62 (s, 1H), 8.20 (dd, *J* = 8.0, 1.8 Hz, 1H), 7.38 (d, *J* = 7.9 Hz, 1H), 7.27 (d, *J* = 2.8 Hz, 2H), 6.91 (d, *J* = 9.4 Hz, 2H), 6.72 (dd, *J* = 9.4, 2.8 Hz, 2H), 3.71 (t, *J* = 5.1 Hz, 2H), 3.67 (q, *J* = 1.2 Hz, 4H), 3.66 – 3.63 (m, 4H), 3.55 – 3.51 (m, 2H), 3.32 (d, *J* = 1.0 Hz, 3H), 0.64 (s, 3H), 0.57 (s, 3H).

**$^{13}$C NMR (126 MHz, MeOH-d4):** δ (ppm) = 168.4, 154.1, 136.7, 132.7, 131.3, 130.6, 129.9, 129.4, 120.8, 115.1, 114.7,72.9, 71.6, 71.4, 71.3, 70.5, 59.1, 41.2, 40.9, –0.5, –1.7.

**MS** (HR-ESI, pos): meas. m/z = 618.3003, calc. m/z = 618.2994 for $C_{34}H_{44}N_3O_6Si$ $[M+H]^+$.

**2-(3-(dimethylamino)-7-(dimethyliminio)-5,5-dimethyl-2,3,5,7-tetrahydrodibenzo[b,e]silin10-yl)-5-((2-(2-(2hydroxyethoxy)ethoxy)ethyl)carbamoyl)benzoate (SiR-2)**

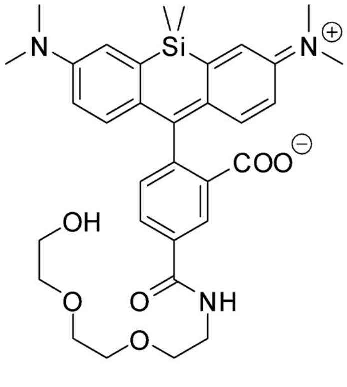

**SiR-2**

**SiR-2** was prepared according to **GP1.** The crude product was purified via normal phase silica column chromatography ($CHCl_3$:MeOH/$NEt_3$ 100:2:0.5) and reverse phase HPLC (0.1 % TFA, $H_2O$:MeCN). **SiR-2** was obtained as a blue solid (5.32 mg, 8.81 µmol, 42 %).

**$^1$H NMR (500 MHz, MeOH-d4):** δ (ppm) = 8.59 (s, 1H), 8.19 (dd, *J* = 8.0, 1.8 Hz, 1H), 7.41 – 7.35 (m, 1H), 7.23 (d, *J* = 2.8 Hz, 2H), 6.88 (d, *J* = 9.3 Hz, 2H), 6.71 (dd, *J* = 9.4, 2.8 Hz, 2H), 3.76 – 3.70 (m, 2H), 3.69 (dtq, *J* = 4.7, 3.3, 1.7 Hz, 3H), 3.67 – 3.63 (m, 4H), 3.58 (dd, *J* = 5.4, 3.9 Hz, 2H), 3.19 (s, 12H), 0.64 (s, 3H), 0.57 (s, 3H).

**$^{13}$C NMR (126 MHz, MeOH-d4):** δ (ppm) = 168.5, 153.8, 136.8, 132.9, 131.3, 120.3, 73.7, 71.5, 71.4, 70.5, 62.2, 41.1, 40.8, –0.4 –1.7.

**MS** (HR-ESI, pos): meas. m/z = 604.2837, calc. m/z = 604.2842 for $C_{33}H_{42}N_3O_6Si$ $[M+H]^+$.

**5-((2-(2-(carboxymethoxy)ethoxy)ethyl)carbamoyl)-2-(7-(dimethylamino)-3-(dimethyliminio)5,5-dimethyl-3,5-dihydrodibenzo[b,e]silin-10-yl)benzoate (SiR-3)**

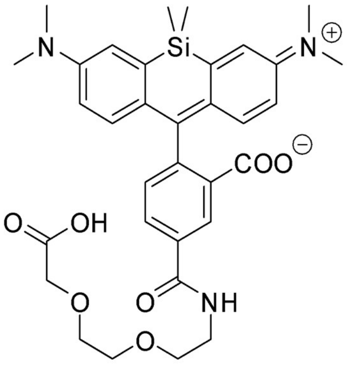

**SiR-3**

**SiR-3** was prepared according to **GP1.** The crude product was purified via normal phase silica column chromatography ($CHCl_3$:MeOH 30:1 to 2:1) and reverse phase HPLC (0.1 % TFA, $H_2O$/MeCN). **SiR-3** was obtained as a blue solid (3.3 mg, 5.3 µmol, 17%).

**$^{1}$H NMR (500 MHz, MeOH-d4):** δ (ppm) = 8.64 (s, 1H), 8.20 (dd, *J* = 8.0, 1.8 Hz, 1H), 7.39 (d, *J* = 7.9 Hz, 1H), 7.28 (d, *J* = 2.5 Hz, 2H), 6.93 (d, *J* = 9.5 Hz, 2H), 6.73 (dd, *J* = 9.5, 2.9 Hz, 2H), 4.14 (s, 2H), 3.77 – 3.70 (m, 6H), 3.66 (t, *J* = 5.2 Hz, 2H), 3.25 (s, 12H), 0.64 (s, 3H), 0.58 (s, 3H).
**$^{13}$C NMR (126 MHz, MeOH-d4):** δ (ppm) = 168.4, 139.1, 115.1, 105.5, 72.0, 71.7, 71.3, 71.3, 71.2, 70.5, 69.0, 41.1, 40.9, 39.8, -0.4, -1.8.

**MS** (HR-ESI, neg): meas. m/z = 616.2475, calc. m/z = 616.2485 for $C_{33}H_{38}N_3O_7Si$ $[M-H]^-$.

**5-((2-(2-(2-acetamidoethoxy)ethoxy)ethyl)carbamoyl)-2-(7-(dimethylamino)-3-(dimethyliminio)-5,5-dimethyl-3,5-dihydrodibenzo[b,e]silin-10-yl)benzoate SiR-4**

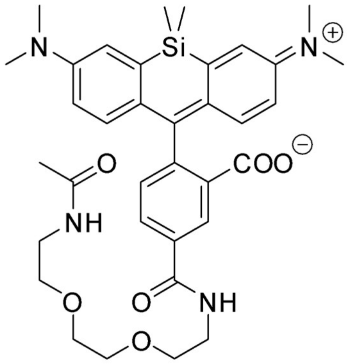

**SiR-4**

**SiR-4** was prepared according to **GP1.** The crude product was purified via normal phase silica column chromatography ($CHCl_3$:MeOH 50:1 to 30:1) and reverse phase HPLC (0.1 % TFA, $H_2O$/MeCN). **SiR-4** was obtained as a blue solid (6.45 mg, 10.0 µmol, 47 %).

**$^{1}$H NMR (500 MHz, MeOH-d4):** δ (ppm) = 8.70 – 8.61 (m, 1H), 8.19 (dd, *J* = 7.9, 1.8 Hz, 1H), 7.39 (d, *J* = 7.9 Hz, 1H), 7.29 (d, *J* = 2.8 Hz, 2H), 6.93 (d, *J* = 9.5 Hz, 2H), 6.73 (dd, *J* = 9.5, 2.9 Hz, 2H), 3.73 (t, *J* = 5.4 Hz, 2H), 3.71 – 3.68 (m, 2H), 3.66 (ddd, *J* = 5.8, 3.5, 2.2 Hz, 4H), 3.55 (t, *J* = 5.5 Hz, 2H), 3.33 (t, *J* = 5.6 Hz, 2H), 3.25 (s, 12H), 1.92 (s, 3H), 0.64 (s, 3H), 0.58 (s, 3H).

**$^{13}$C NMR (126 MHz, MeOH-d4):** δ (ppm) = 173.4, 168.5, 136.7, 132.6, 131.8, 130.8, 130.0,129.9, 121.0, 115.1, 71.4,70.6, 70.5, 41.1, 40.9, 40.4, 22.6, –0.6, –1.8.

**MS** (HR-ESI, pos): meas. m/z = 645.3116, calc. m/z = 645.3103 for $C_{35}H_{45}N_4O_6Si$ $[M+H]^+$.

**2-(7-(dimethylamino)-3-(dimethyliminio)-5,5-dimethyl-3,5-dihydrodibenzo[b,e]silin-10-yl)-5-((2-(2-(2-(methylsulfonamido)ethoxy)ethoxy)ethyl)carbamoyl)benzoate (SiR-5)**

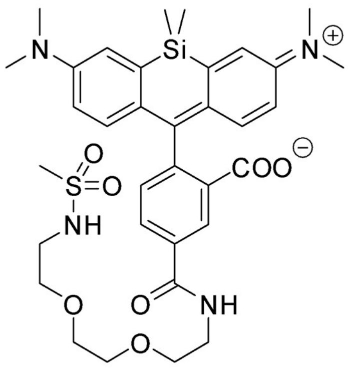

**SiR-5**

**SiR-5** was prepared according to **GP1.** The crude product was purified via normal phase silica column chromatography ($CHCl_3$:MeOH 50:1) and reverse phase HPLC (0.1 % TFA, $H_2O$/MeCN). **SiR-5** was obtained as a blue solid (7.3 mg, 10.7 µmol, 34%).

**$^1$H NMR (500 MHz, MeOH-d4):** δ (ppm) = 8.66 (s, 1H), 8.20 (dd, *J* = 7.9, 1.8 Hz, 1H), 7.40 (d, *J* = 7.9 Hz, 1H), 7.30 (d, *J* = 2.8 Hz, 2H), 6.95 (d, *J* = 9.5 Hz, 2H), 6.75 (dd, *J* = 9.5, 2.8 Hz, 2H), 3.73 (t, *J* = 5.4 Hz, 2H), 3.71 – 3.65 (m, 4H), 3.60 (t, *J* = 5.3 Hz, 2H), 3.27 (s, 12H), 3.23 (t, *J* = 5.3 Hz, 2H), 2.92 (s, 3H), 0.64 (s, 3H), 0.58 (s, 3H).

**$^{13}$C NMR (126 MHz, MeOH-d4):** δ (ppm) = 168.5, 136.6, 115.2, 71.4, 71.4, 71.4, 70.5, 44.0, 41.1, 40.9, 40.4, -0.6, -1.8.

**MS** (HR-ESI, pos): meas. m/z = 681.2793, calc. m/z = 681.2773 for $C_{34}H_{45}N_4O_7SSi$ $[M+H]^+$.

**2-(7-(dimethylamino)-3-(dimethyliminio)-5,5-dimethyl-3,5-dihydrodibenzo[b,e]silin-10-yl)-5-((2-(2-(2-((trifluoromethyl)sulfonamido)ethoxy)ethoxy)ethyl)carbamoyl)benzoate (SiR-6)**

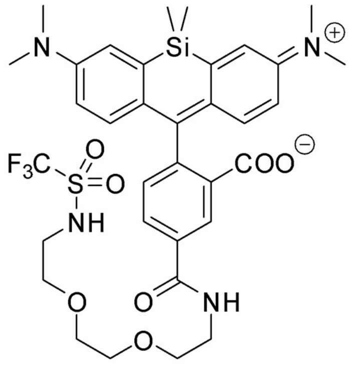

**SiR-6**

**SiR-6** was prepared according to **GP1.** The crude product was purified via normal phase silica column chromatography ($CHCl_3$:MeOH 50:1) and reverse phase HPLC (0.1 % TFA, $H_2O$/MeCN). **SiR-6** was obtained as a blue solid (8.7 mg, 11.8 µmol, 37 %).

**$^1$H NMR (500 MHz, MeOH-d4):** δ (ppm) = 8.68 (d, *J* = 1.8 Hz, 1H), 8.19 (dd, *J* = 7.9, 1.8 Hz, 1H), 7.39 (d, *J* = 7.9 Hz, 1H), 7.32 (d, *J* = 2.8 Hz, 2H), 6.96 (d, *J* = 9.5 Hz, 2H), 6.74 (dd, *J* = 9.5, 2.8 Hz, 2H), 3.74 (t, *J* = 5.4 Hz, 2H), 3.73 – 3.69 (m, 2H), 3.69 – 3.65 (m, 4H), 3.60 (t, *J* = 5.3 Hz, 2H), 3.36 (t, *J* = 5.3 Hz, 2H), 3.29 (s, 12H), 0.64 (s, 3H), 0.58 (s, 3H).

**$^{13}$C NMR (126 MHz, MeOH-d4):** δ (ppm) = 168.5, 136.6, 121.4, 115.1, 71.4, 71.4, 71.0, 70.5, 44.8, 41.0, 40.9, -0.7, -1.8.

**$^{19}$F NMR (282 MHz, MeOH-d4):** δ (ppm)= -79.43 (s).

**MS** (HR-ESI, pos): meas. m/z =735.2484, calc. m/z = 735.2490 for $C_{34}H_{42}F_3N_4O_7SSi$ $[M+H]^+$.

### NMR spectra

1H-NMR (500 MHz, CDCl3)

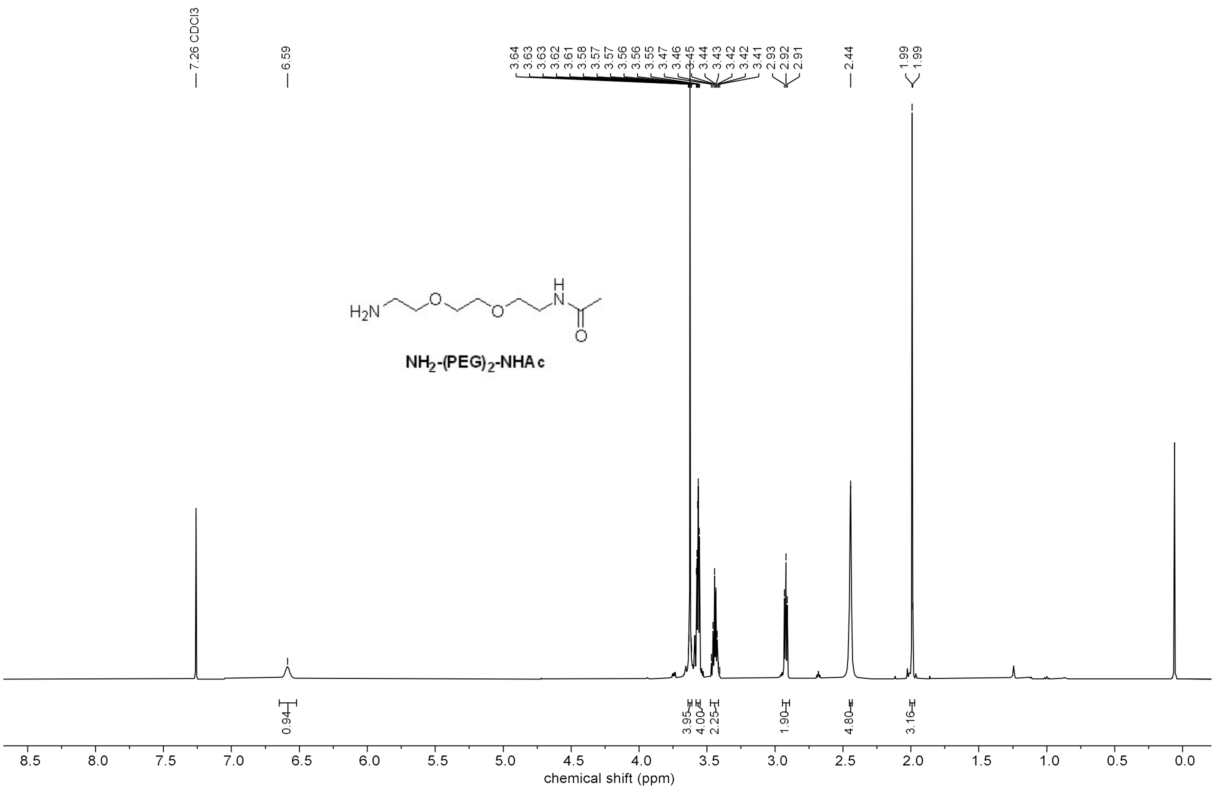

13C-NMR (126 MHz, CDCl3)

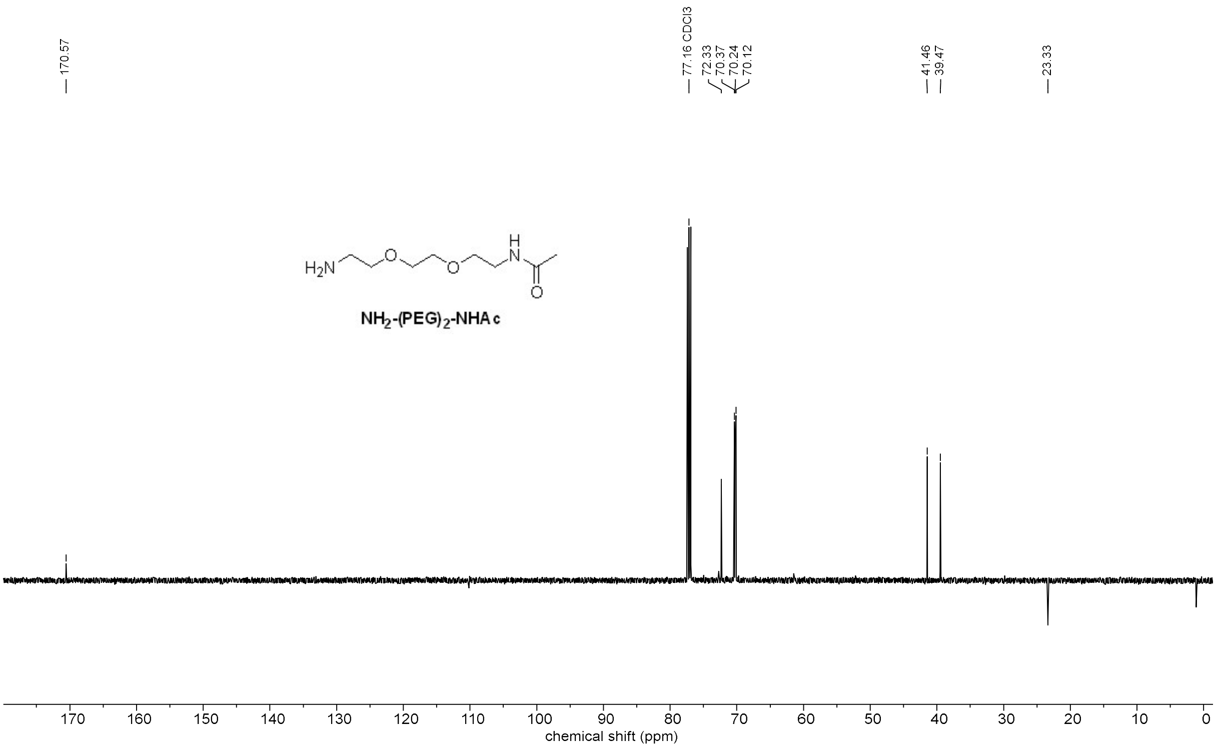

1H-NMR (500 MHz, CDCl3)

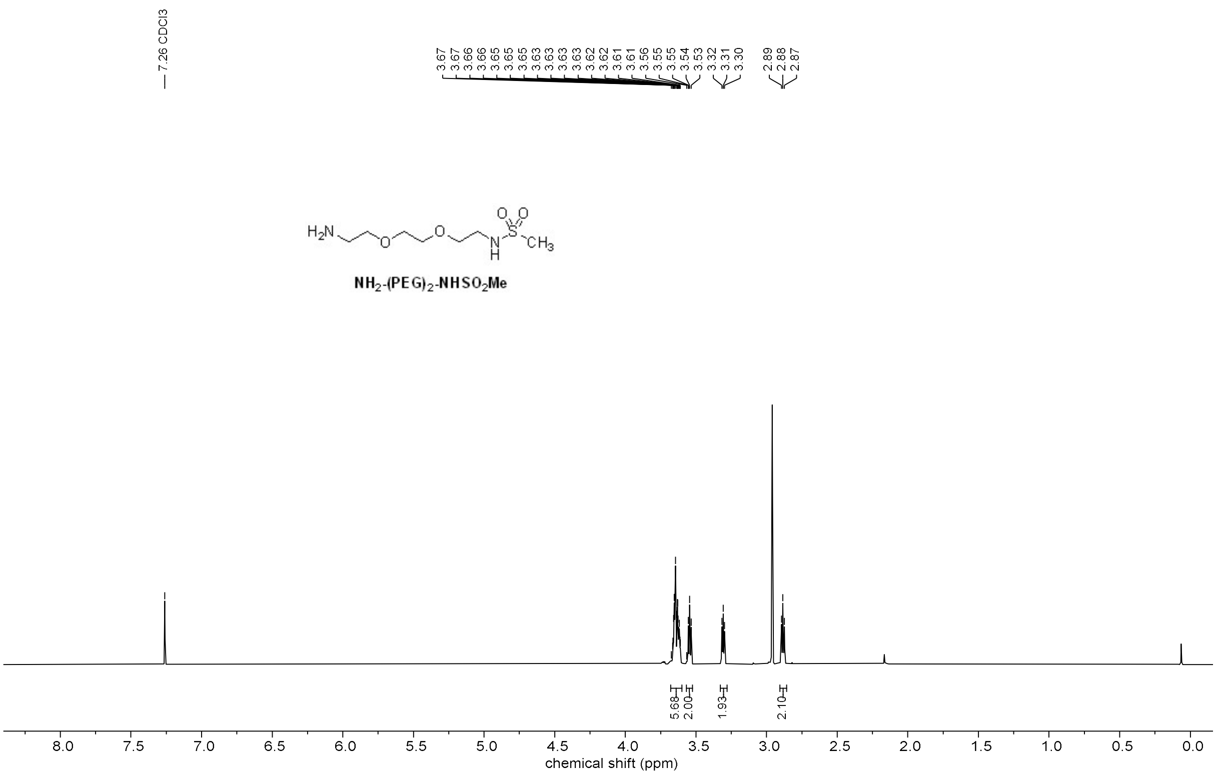

13C-NMR (126 MHz, CDCl3)

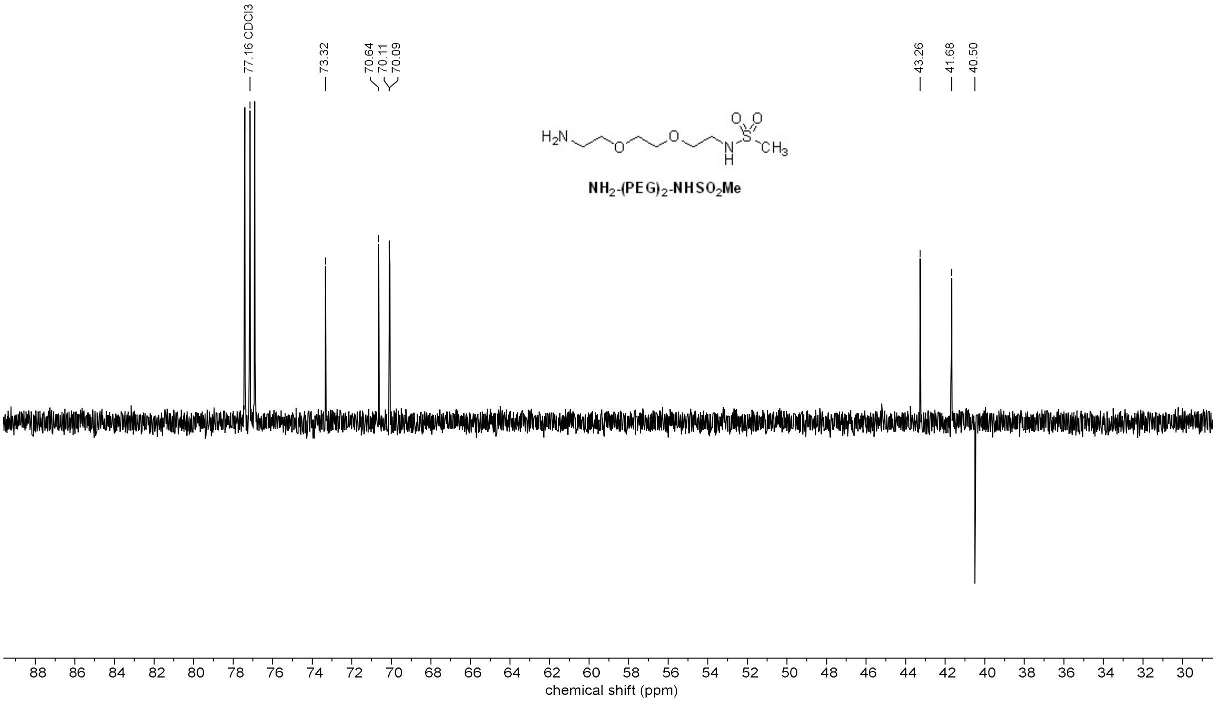

1H-NMR (500 MHz, MeOH-d4)

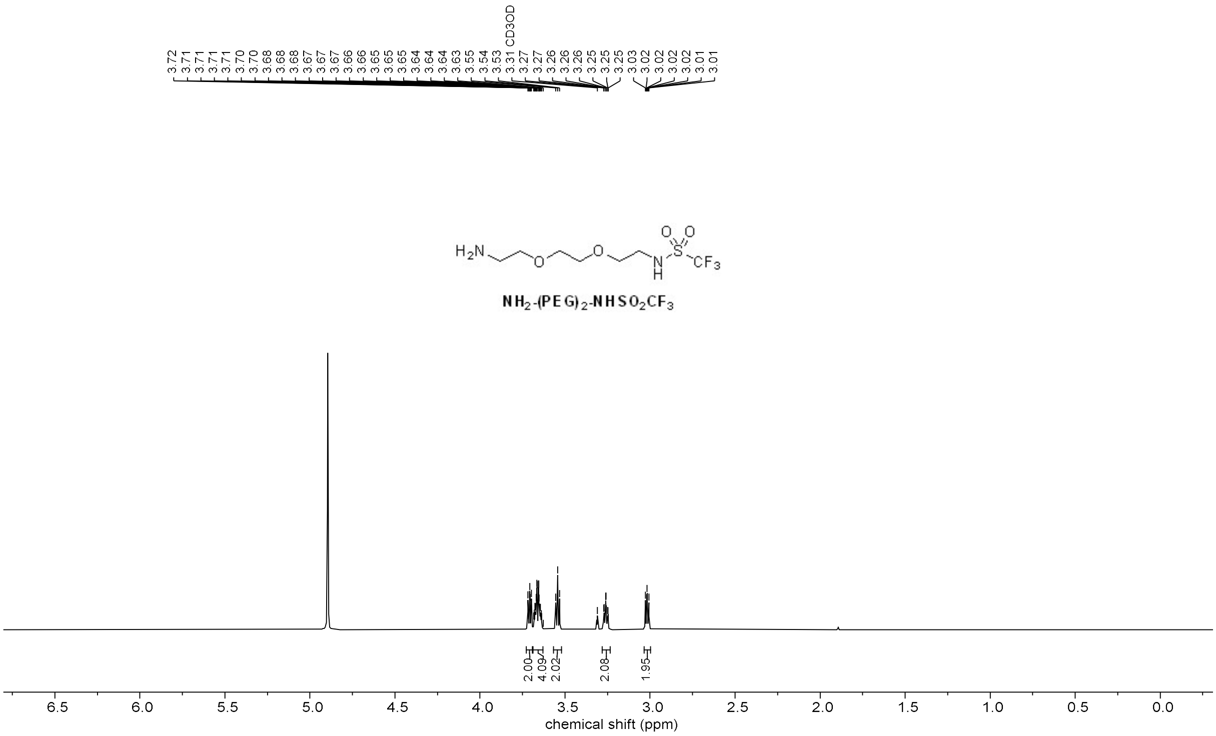

13C-NMR (126 MHz, MeOH-d4)

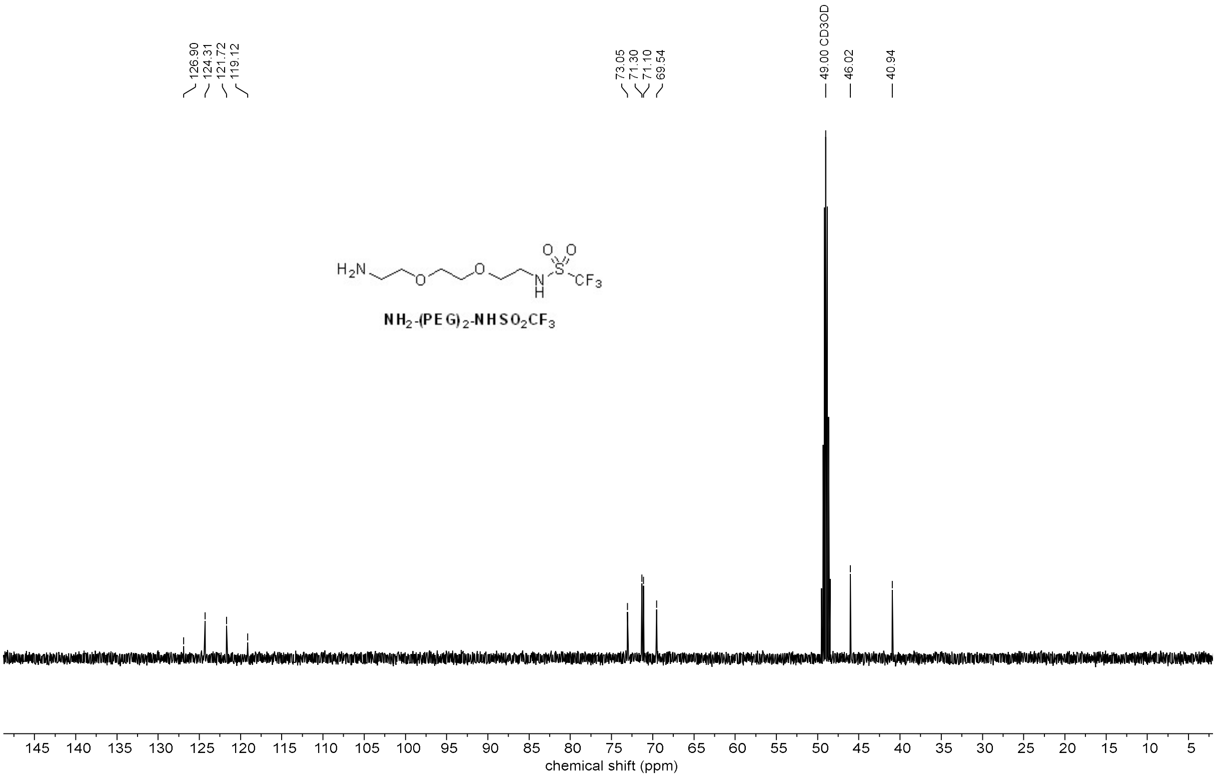

19F-NMR (282 MHz, MeOH-d4)

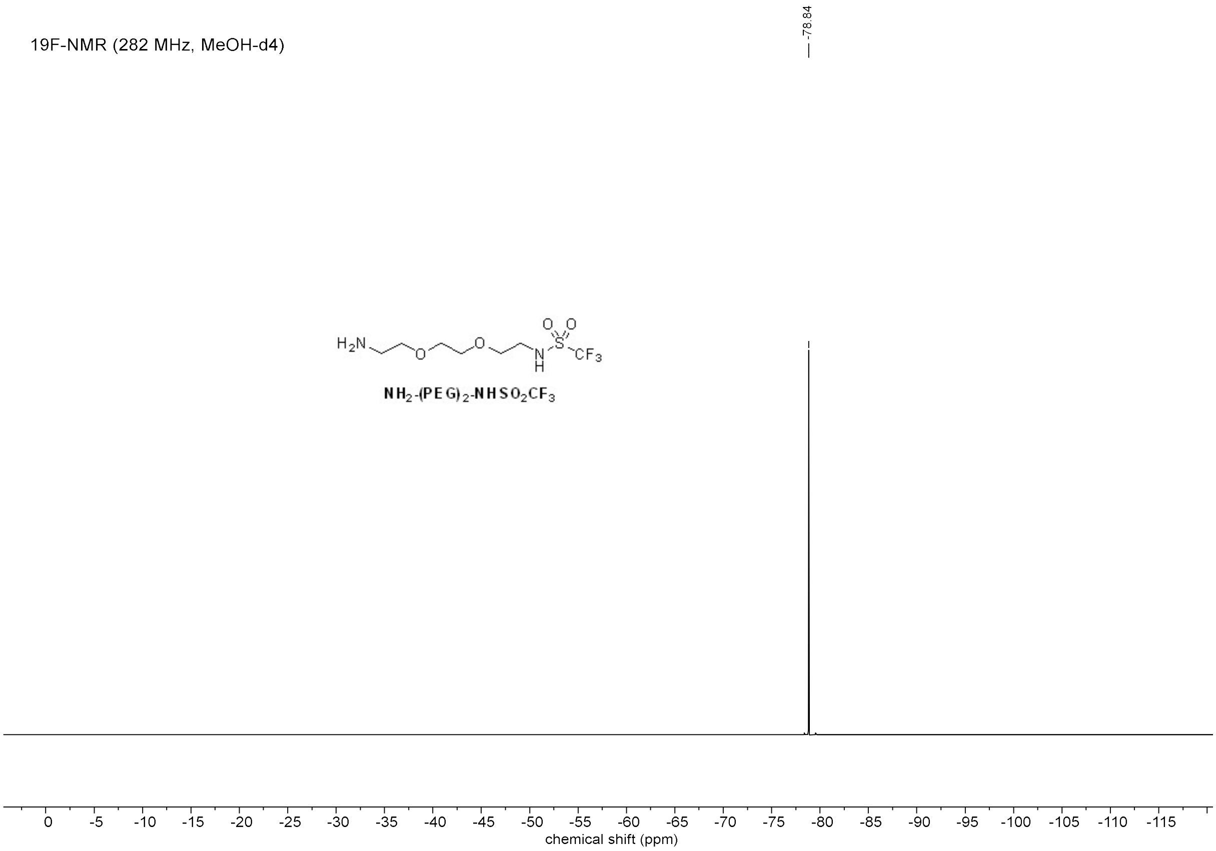

1H-NMR (500 MHz, MeOH-d4)

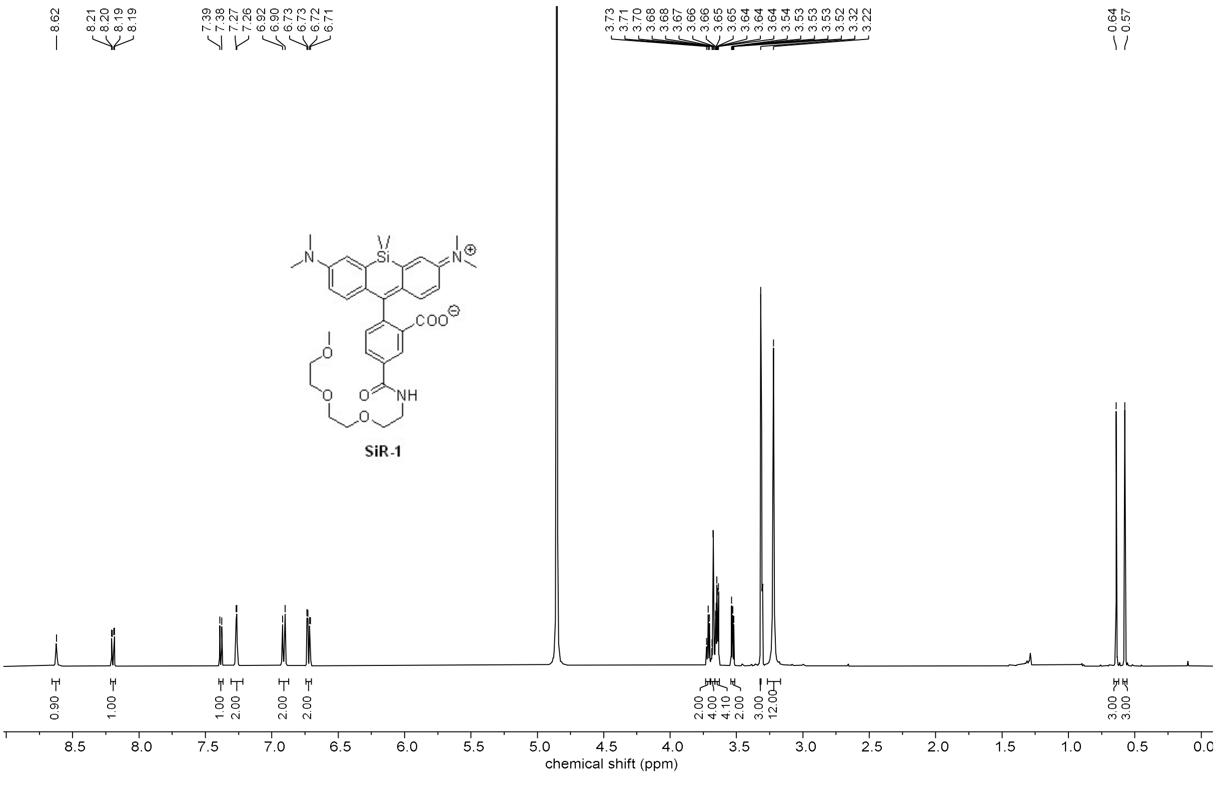

13C-NMR (126 MHz, MeOH-d4)

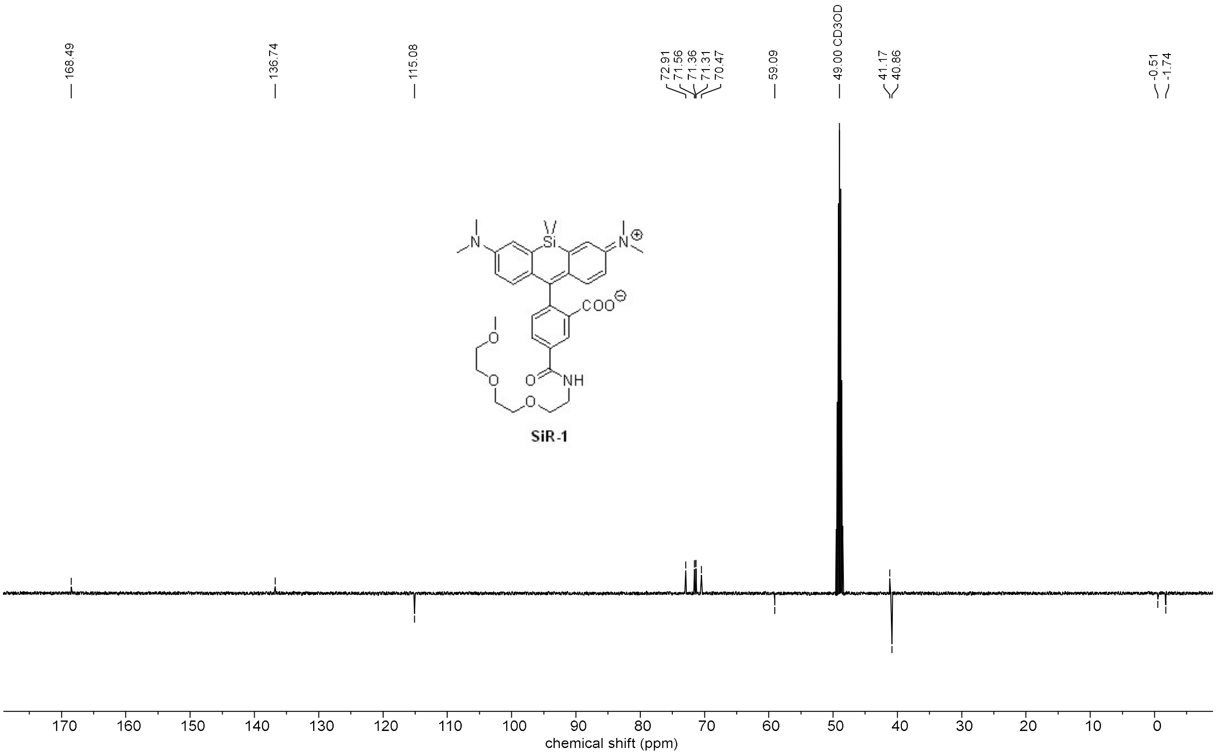

1H-NMR (500 MHz, MeOH-d4)

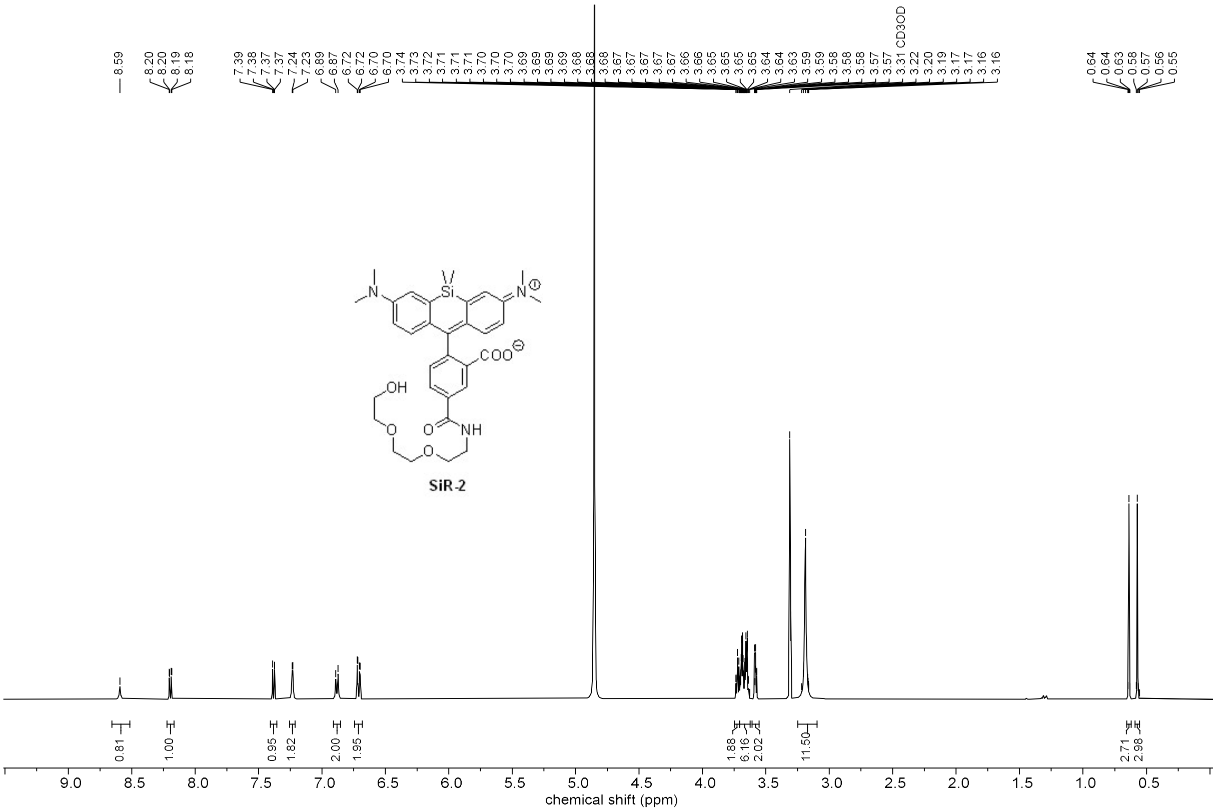

13C-NMR (126 MHz, MeOH-d4)

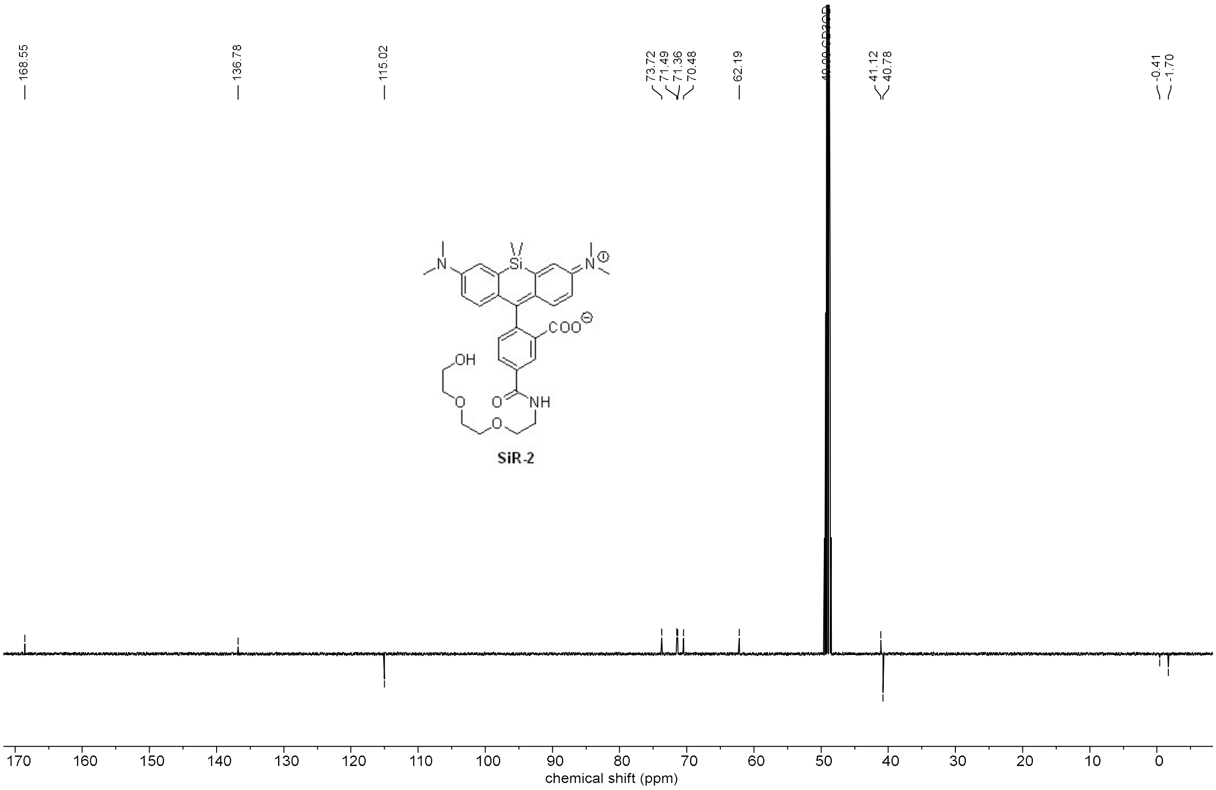

1H-NMR (500 MHz, MeOH-d4)

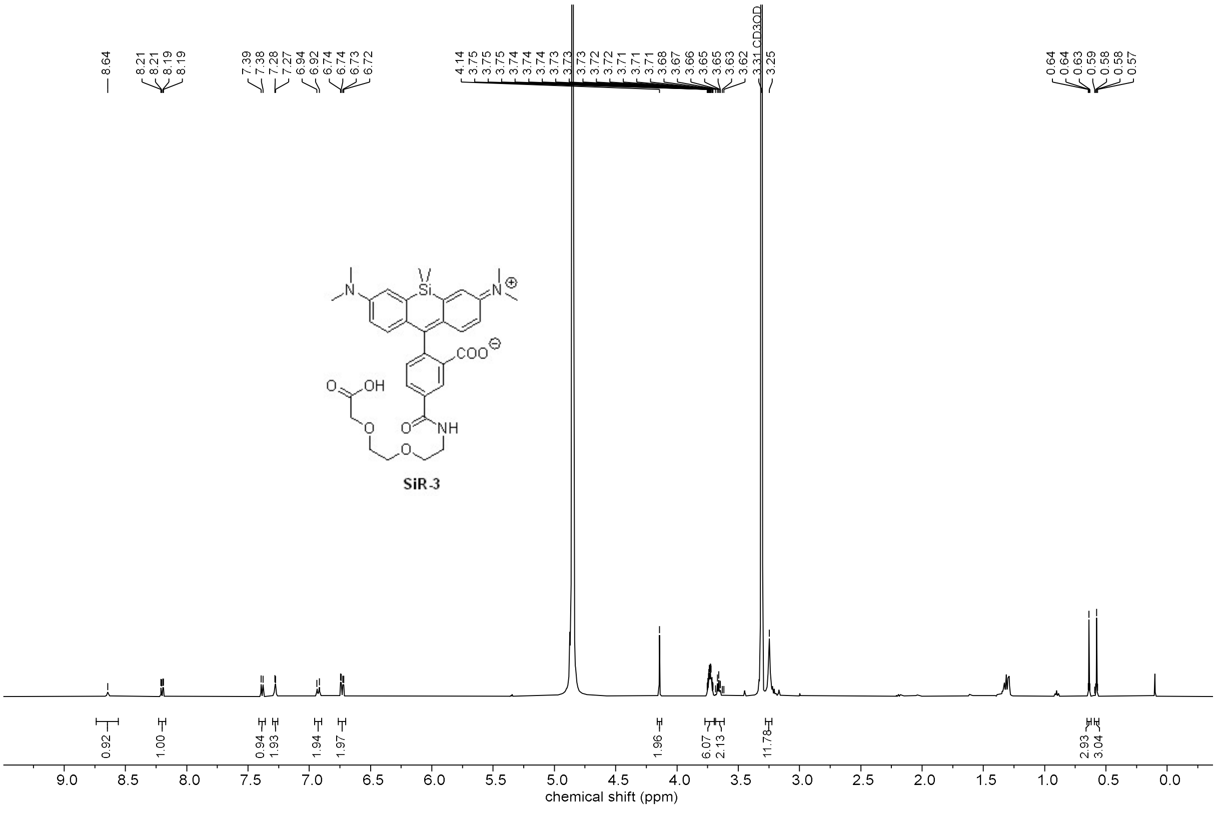

13C-NMR (126 MHz, MeOH-d4)

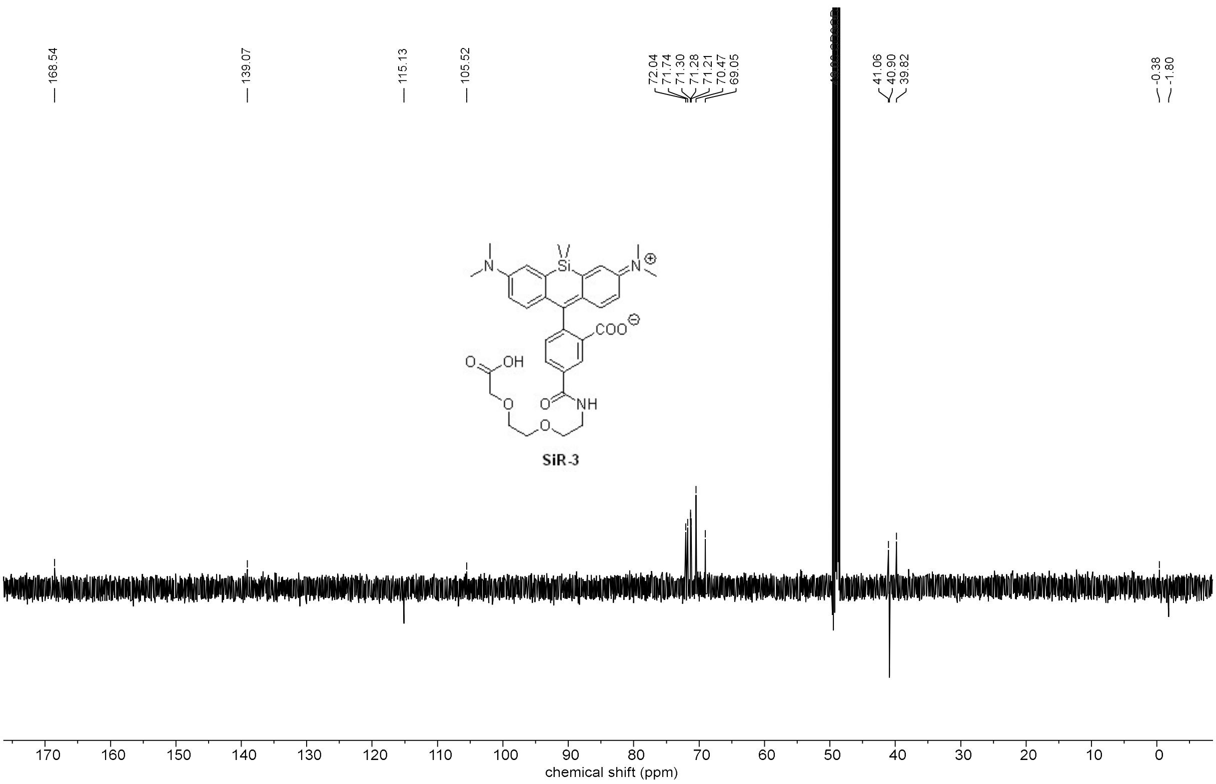

1H-NMR (500 MHz, MeOH-d4)

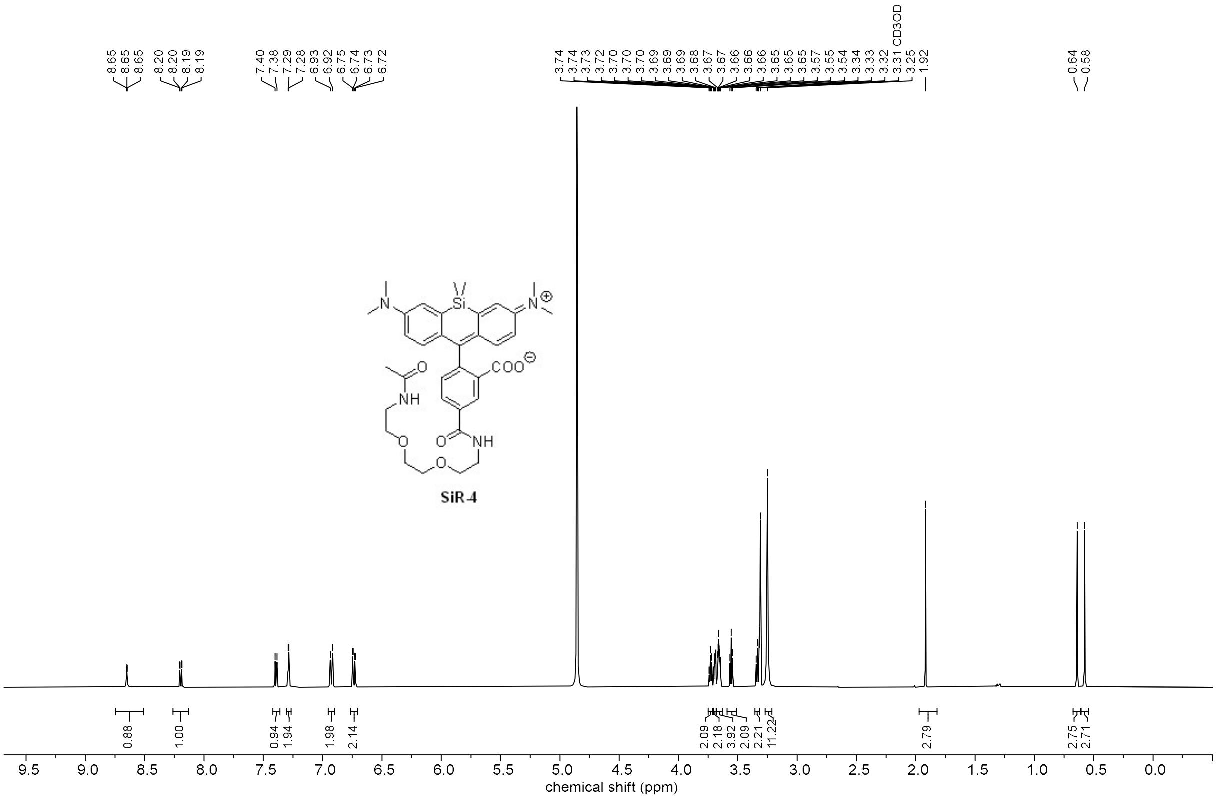

13C-NMR (126 MHz, MeOH-d4)

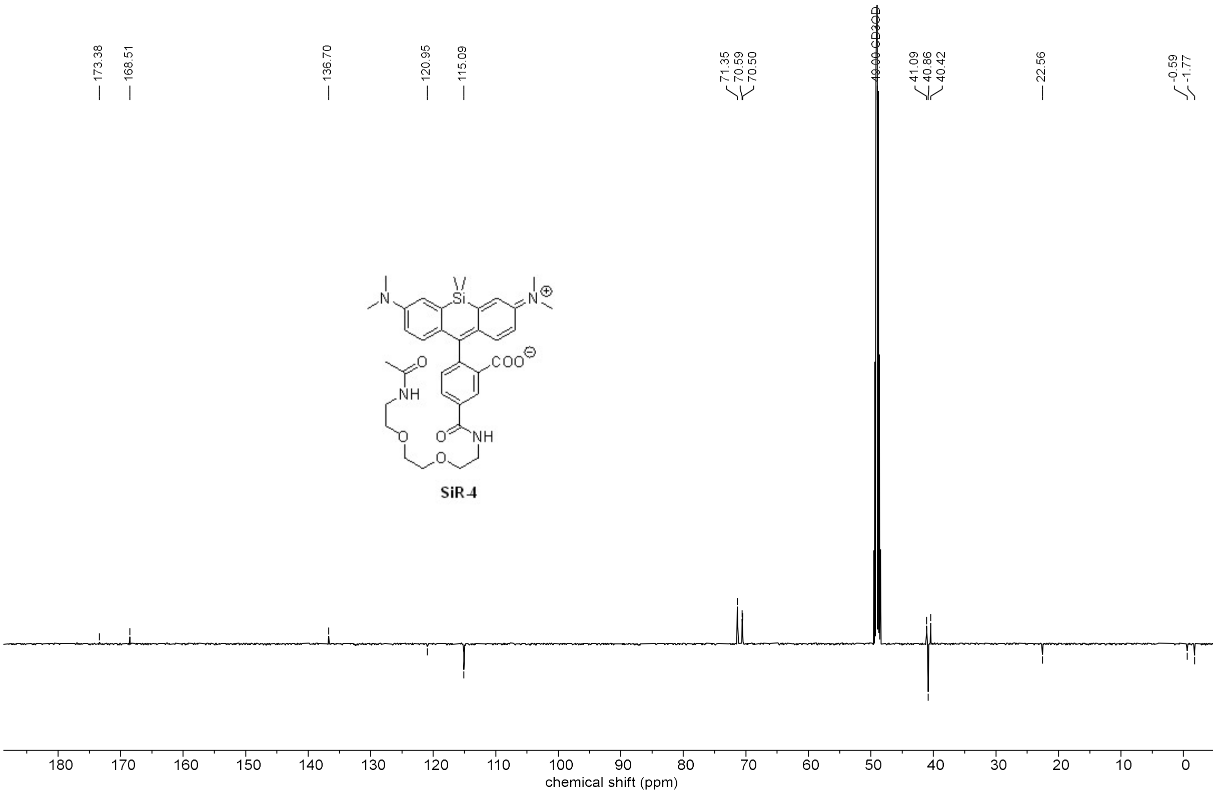

1H-NMR (500 MHz, MeOH-d4)

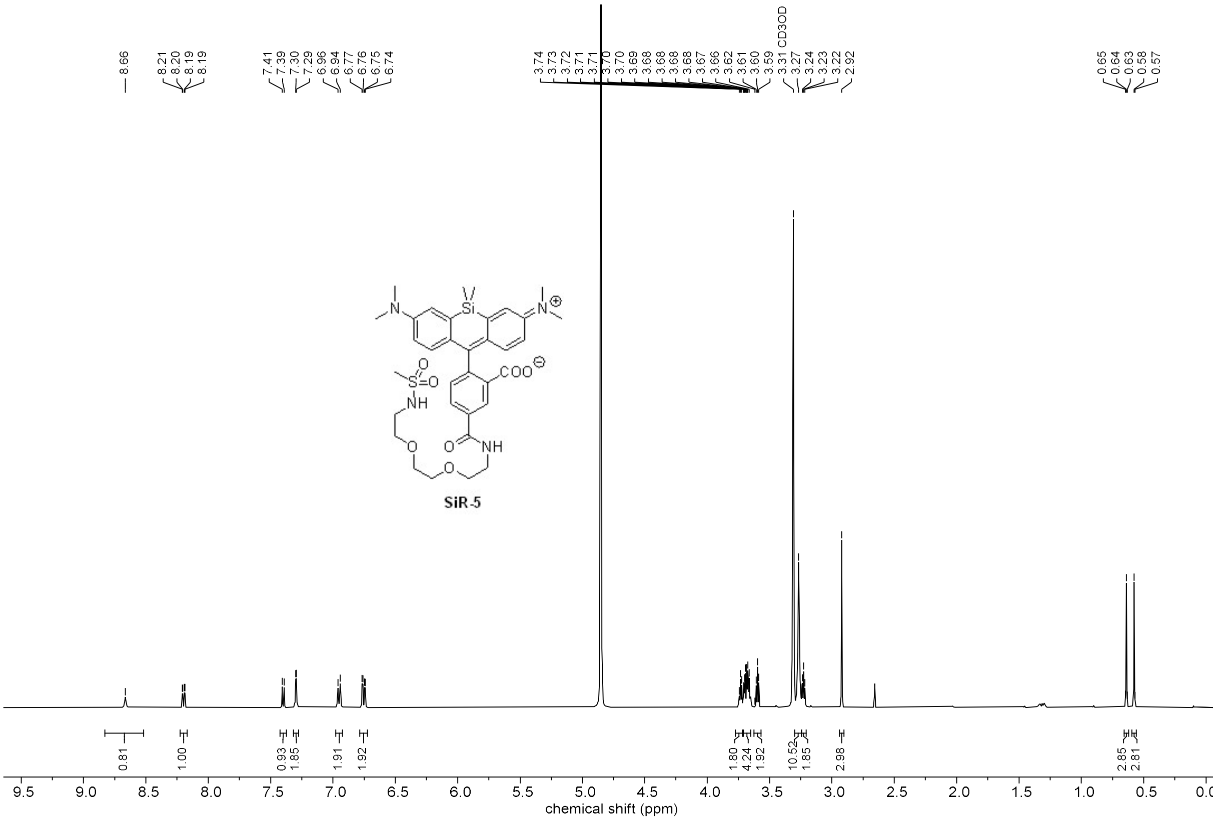

13C-NMR (126 MHz, MeOH-d4)

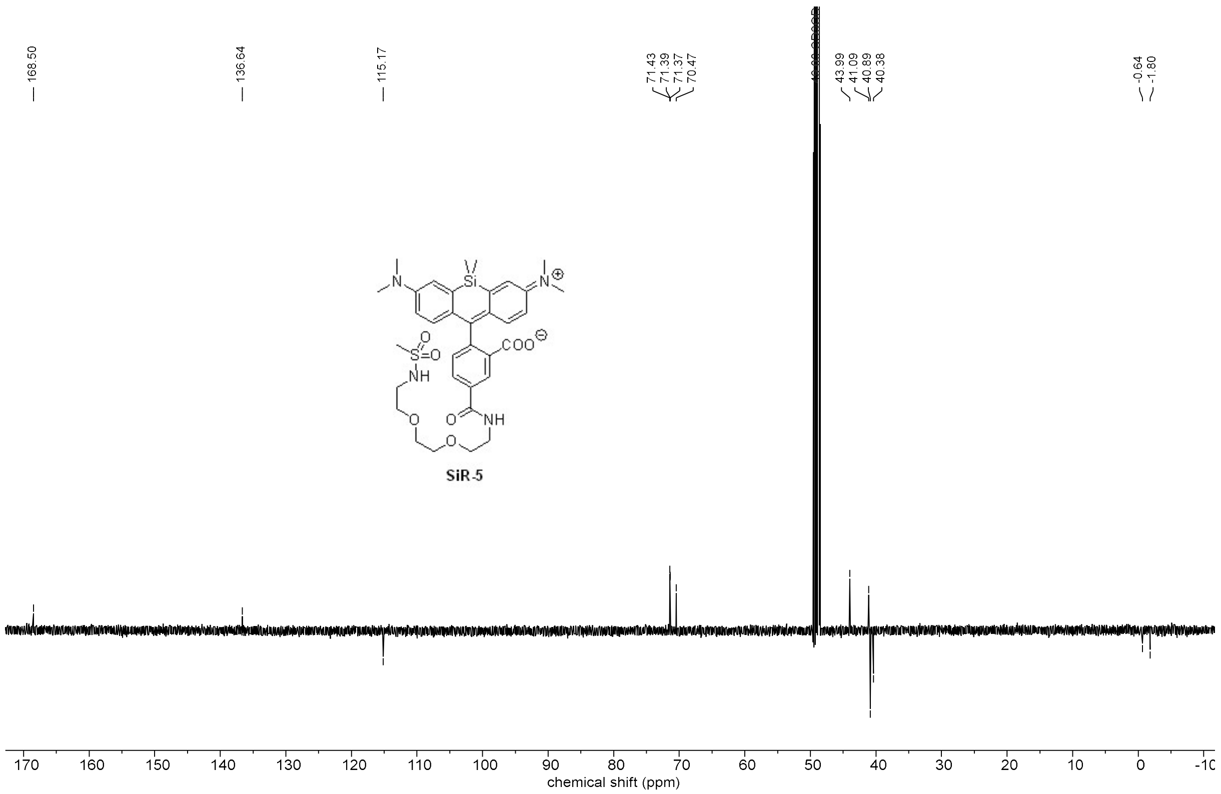

1H-NMR (500 MHz, MeOH-d4)

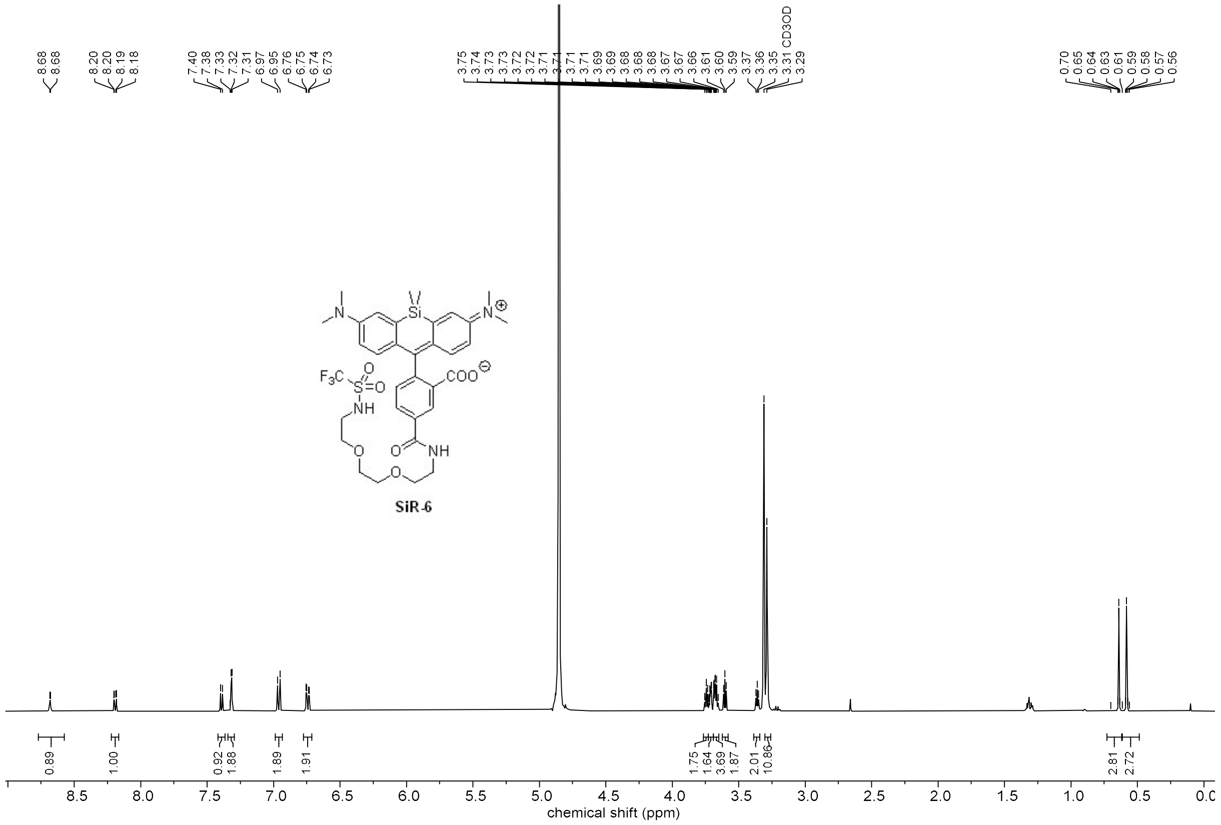

13C-NMR (126 MHz, MeOH-d4)

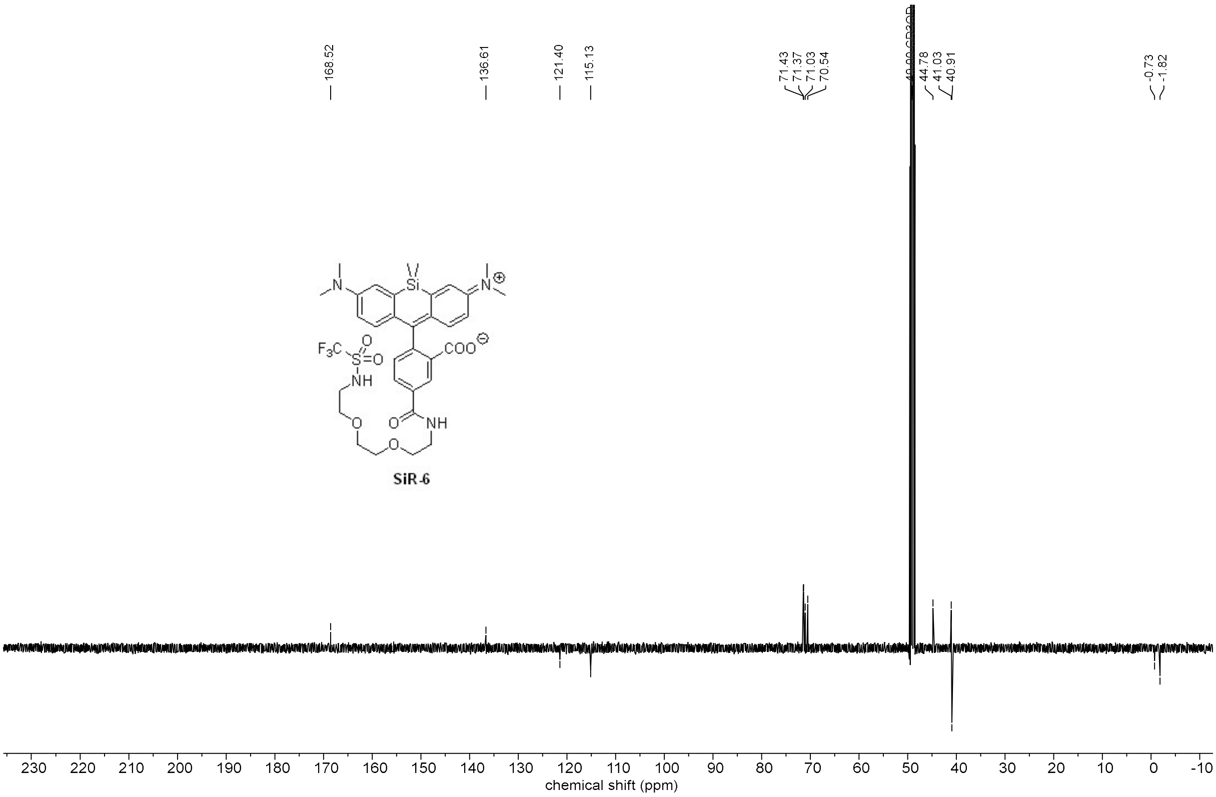

19F-NMR (282 MHz, MeOH-d4)

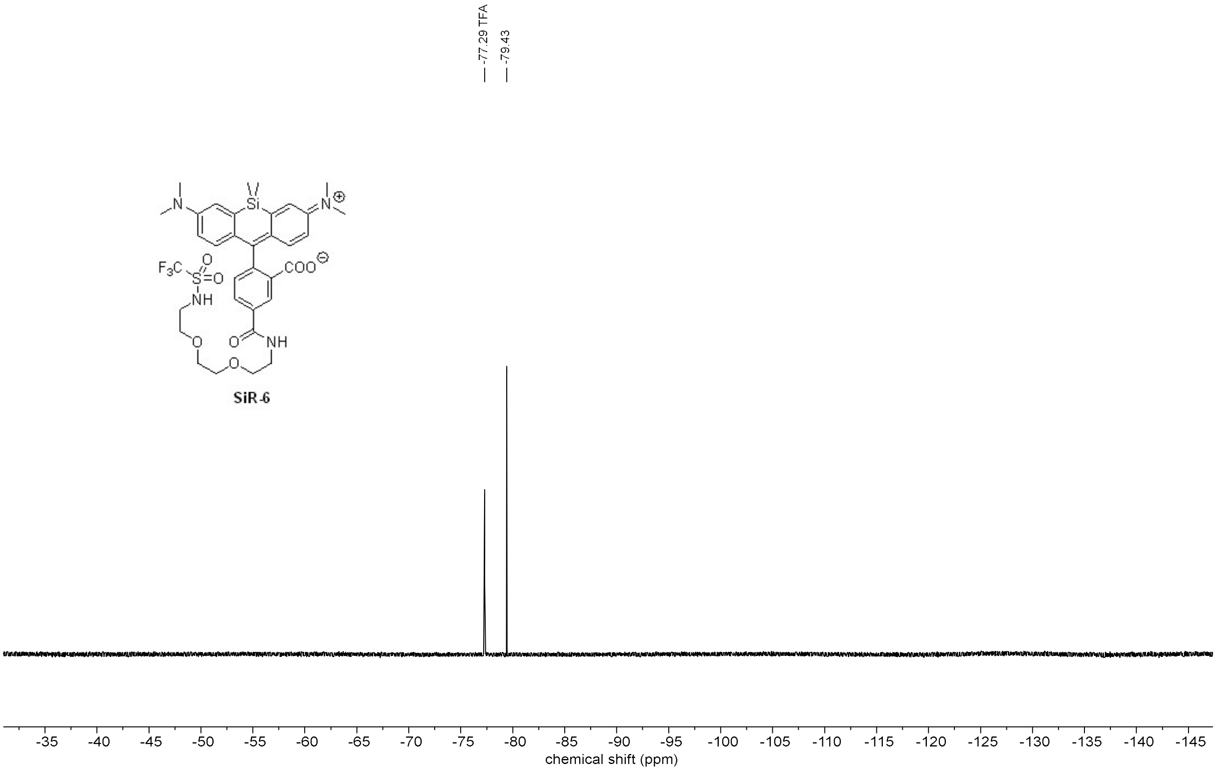

### Supplementary data

#### Supplementary figures

##### Quantification of SiRA:SiR-NH2 in mammalian live cells

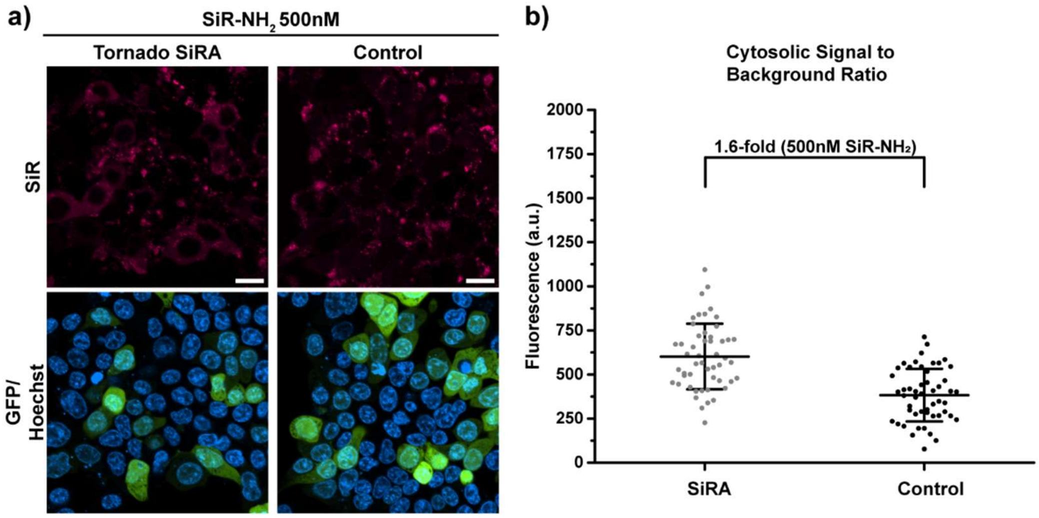

**Figure S1:** Quantification of the fluorescence intensity of SiRA:SiR-$NH_2$ *in cellulo*. a) Confocal imaging of live HEK293T cells expressing circular SiRA or RhoBAST[4](control) with the Tornado expression system[6], incubated with SiR-$NH_2$ (500 nM), showing unspecific accumulation independent of the presence of SiRA. Cells were cotransfected with mEGFP as transfection control. Scalebars, 20 µm. b) Quantification of the cytosolic fluorescence signal in the SiR-channel of individual transfected cells (*n* = 50, repeated in triplicates).

##### IPTG-viability

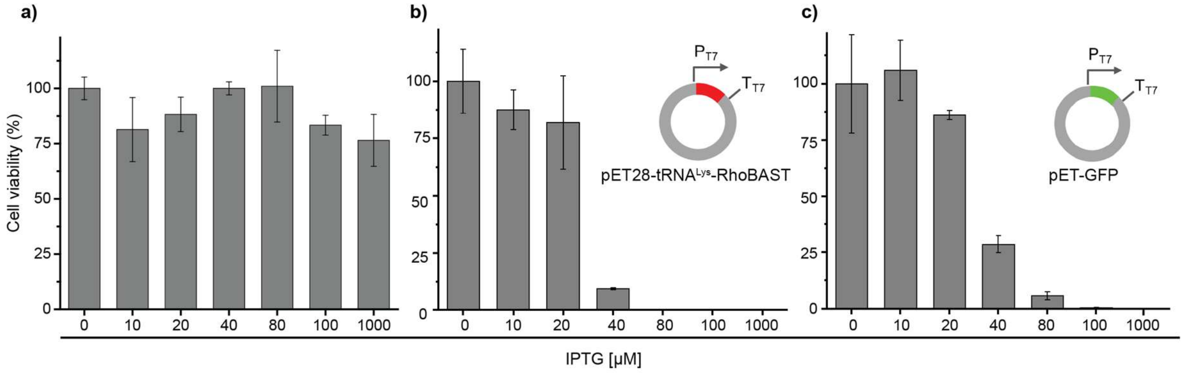

**Figure S2.** Effect of varying IPTG concentrations on the viability of a) untransfected bacteria, b) bacteria transfected with pET28-tRNA$^{Lys}$-RhoBAST (used as a negative control in FACS selection), and c) bacteria transfected with pET-GFP.

#### FACS SELEX with SiR-$NH_2$

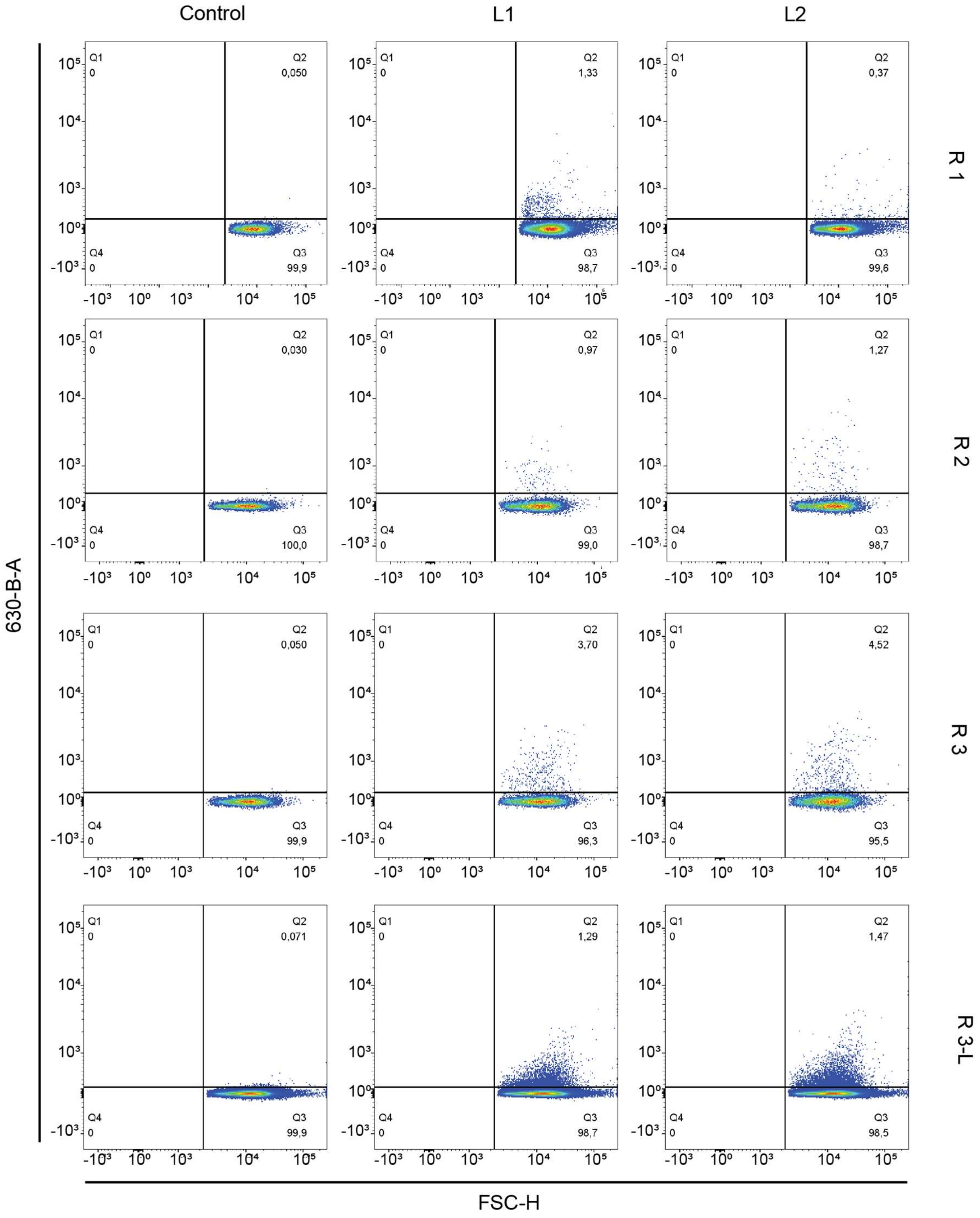

**Figure S3**. Results of FACS SELEX throughout the rounds. Sorting gate was set so <0.1% of cells were sorted for the negative control. Scatter plot y axis shows measured fluorescence; x axis shows front scatter. Logicle -0.2% scale.

#### Candidate evolution throughout FACS SELEX

a)

| | library | R1 | R2 | R3-L |
|---|---|---|---|---|
| L2-1 | 23 | 3 | 1 | 1 |
| L2-2 | 41 | 5 | 4 | 2 |
| L2-3 | -- | 39 | 5 | 3 |
| L2-4 | 47 | 9 | 8 | 4 |
| L2-5 | -- | 32 | 6 | 5 |
| L2-6 | 39 | 2 | 3 | 6 |
| L2-7 | -- | -- | 42 | 7 |
| L2-8 | 39 | -- | 19 | 8 |
| L2-9 | -- | 8 | 14 | 9 |
| L2-10 | -- | 12 | 12 | 10 |

b)

| | library | R1 | R2 | R3-L |
|---|---|---|---|---|
| L1-1 | -- | 6 | 5 | 1 |
| L1-2 | 16 | 4 | 3 | 2 |
| L1-3 | -- | 10 | 7 | 3 |
| L1-4 | 12 | 1 | 2 | 4 |
| L1-5 | -- | 13 | 8 | 5 |
| L1-6 | -- | -- | -- | 6 |
| L1-7 | -- | 36 | 9 | 7 |
| L1-8 | 24 | 3 | 6 | 8 |
| L1-9 | -- | 12 | 1 | 9 |
| L1-10 | -- | 2 | 4 | 10 |

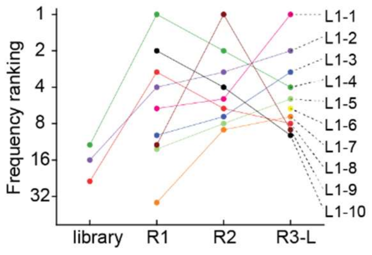

**Figure S4.** Selection results of L1 and data for L2 a) Table of data corresponding to Fig. 2 b). b) Table of data for ranking of candidates throughout the FACS SELEX L1 and graphical representation.

#### Candidate performance in HEK293T cells

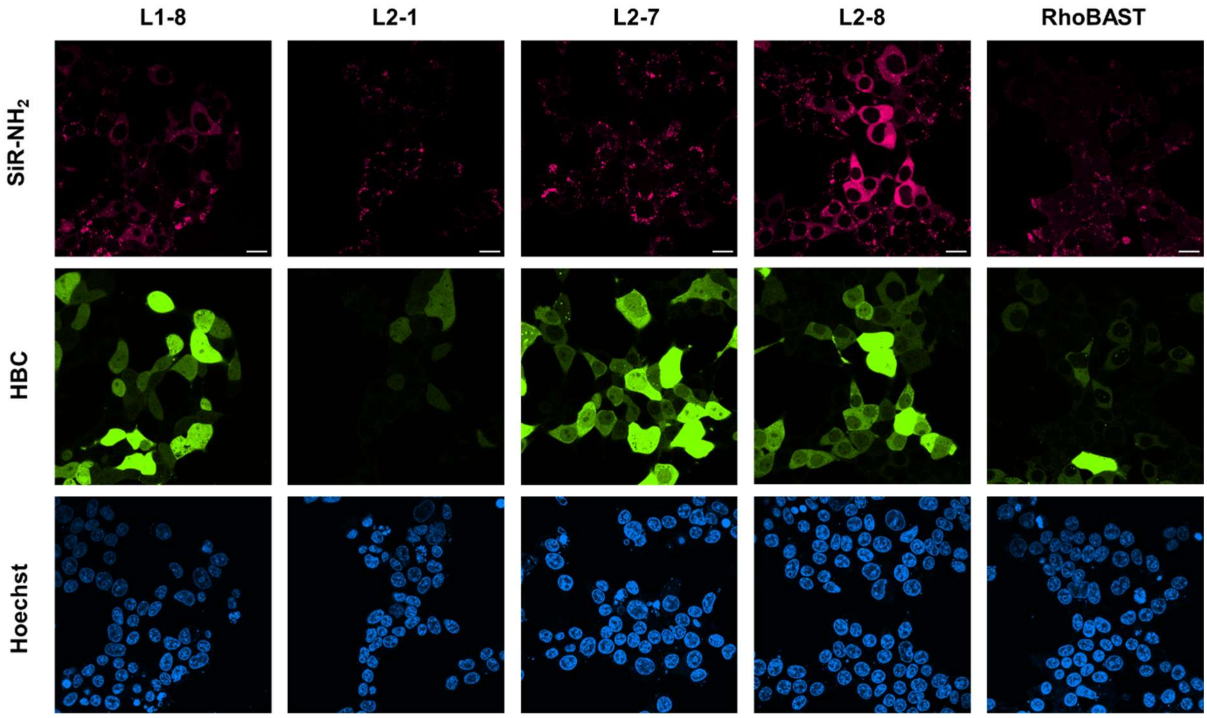

**Figure S5.** Comparison of the highest affinity binders identified in FACS SELEX. Aptamers were introduced into the tetraloop of the pepper aptamer[10] and the construct subsequently embedded into the Tornado overexpression scaffold for RNA expression. Cells were transfected as described in the methods. 1 µM SiR-$NH_2$ and 500 nM HBC530 were added 60 minutes prior to imaging. Experiments were repeated two times with similar results. Scale bars 20 µm.

#### SHAPE probing of aptamer with SiR-$NH_2$

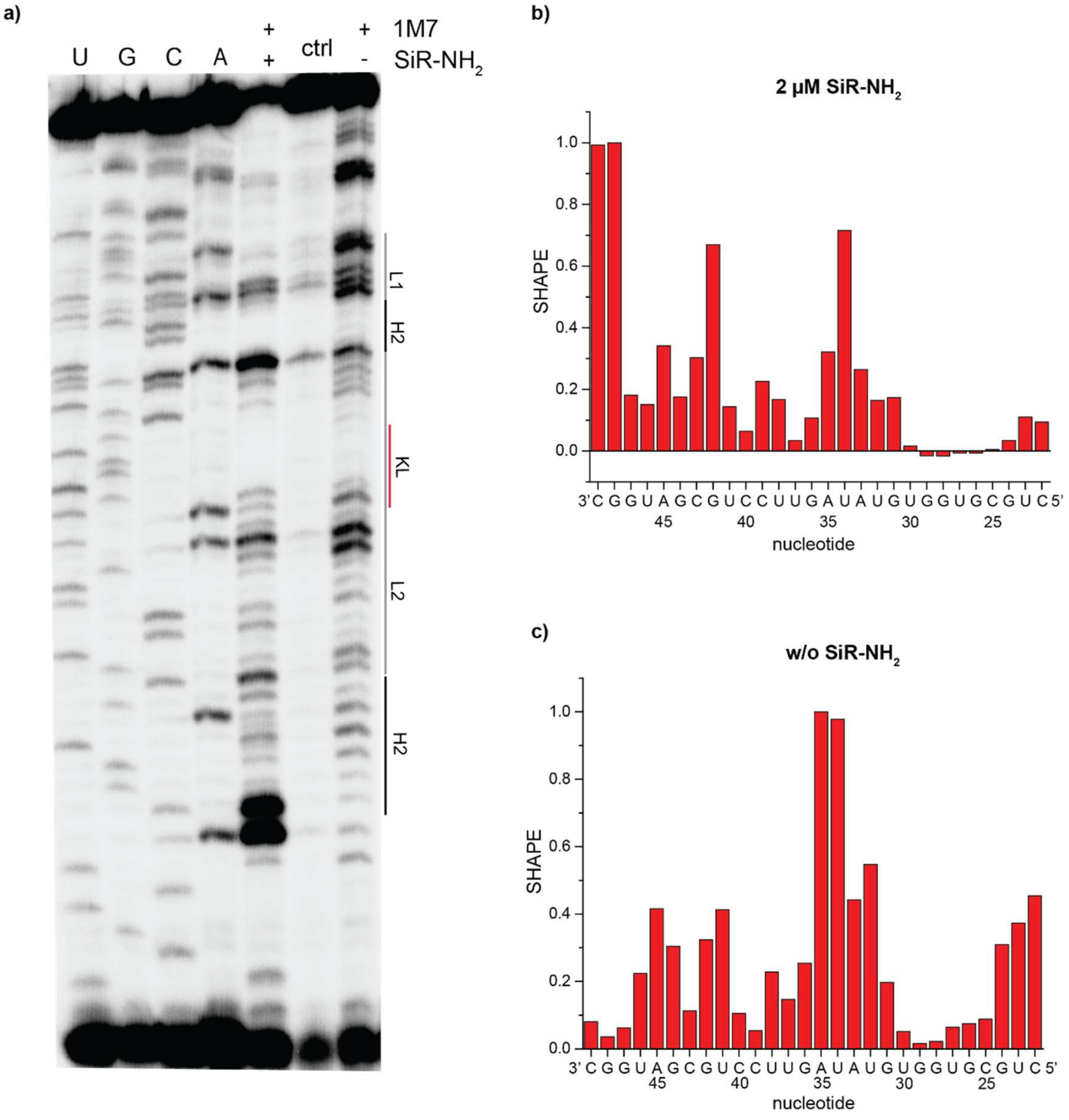

**Figure S6.** SHAPE probing in the presence and absence of SiR-$NH_2$. a) Sequencing gel with sequencing ladder and SHAPE-probed RNA ladder and unmodified control. b) Reactivities of nucleotides when probed in the presence of 2 µM of SiR-$NH_2$. c) Reactivities of nucleotides in the absence of SiR-$NH_2$.

#### $K_D$ curve of v2 with SiR-$NH_2$

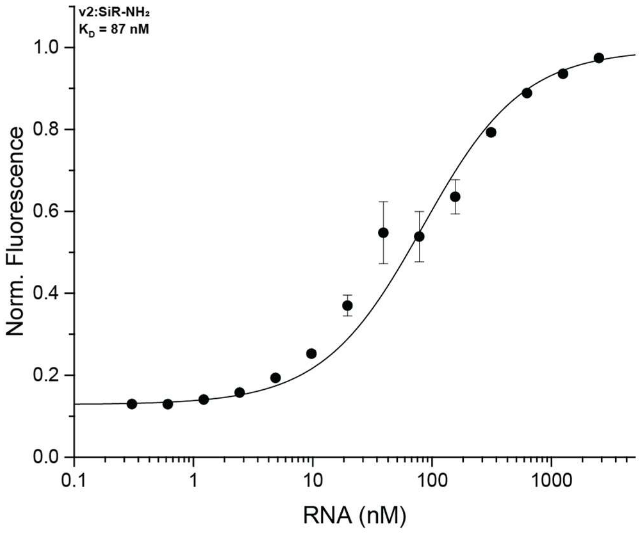

**Figure S7.** Affinity curve of truncated aptamer v2 to SiR-$NH_2$.(10 nM). Measured in triplicates. Measurements were performed in ASB supplemented with 0.05% Tween 20 at 25 °C.

#### Sequences and structures of synonymous repeats

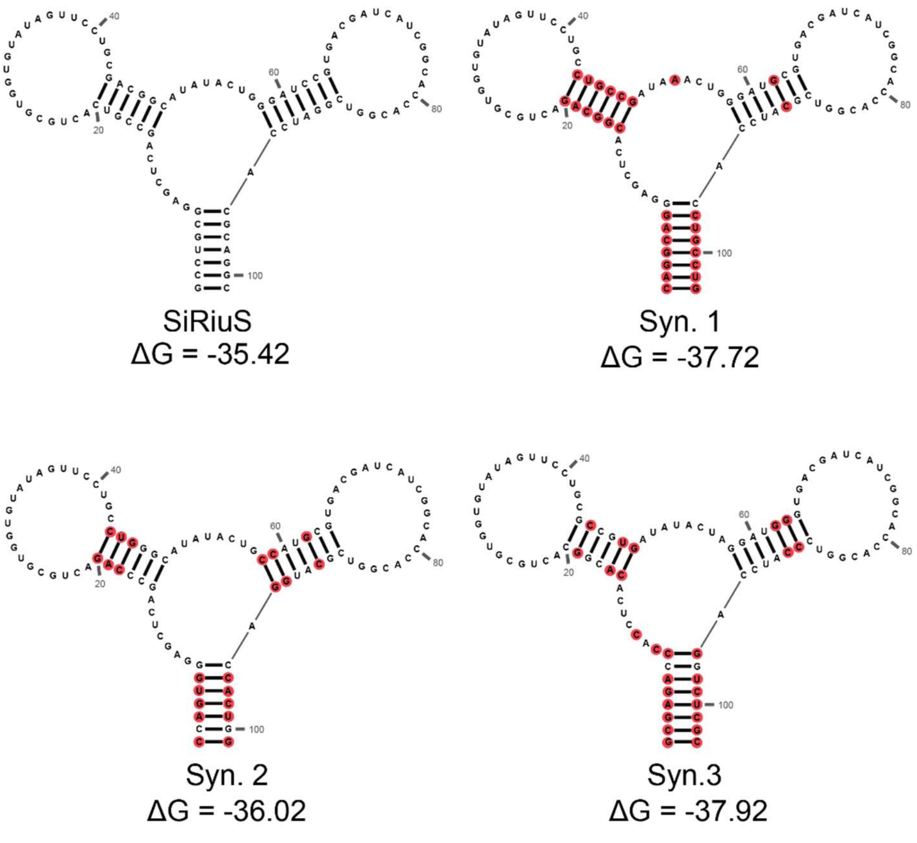

**Figure S8.** Predicted folding and free Gibbs energy of the SiRiuS synonymous repeats as predicted by the Zuker algorithm[11]. Changes to the sequence are marked in red.

#### $K_D$ curves of SiR-R with SiRA and SiRiuS

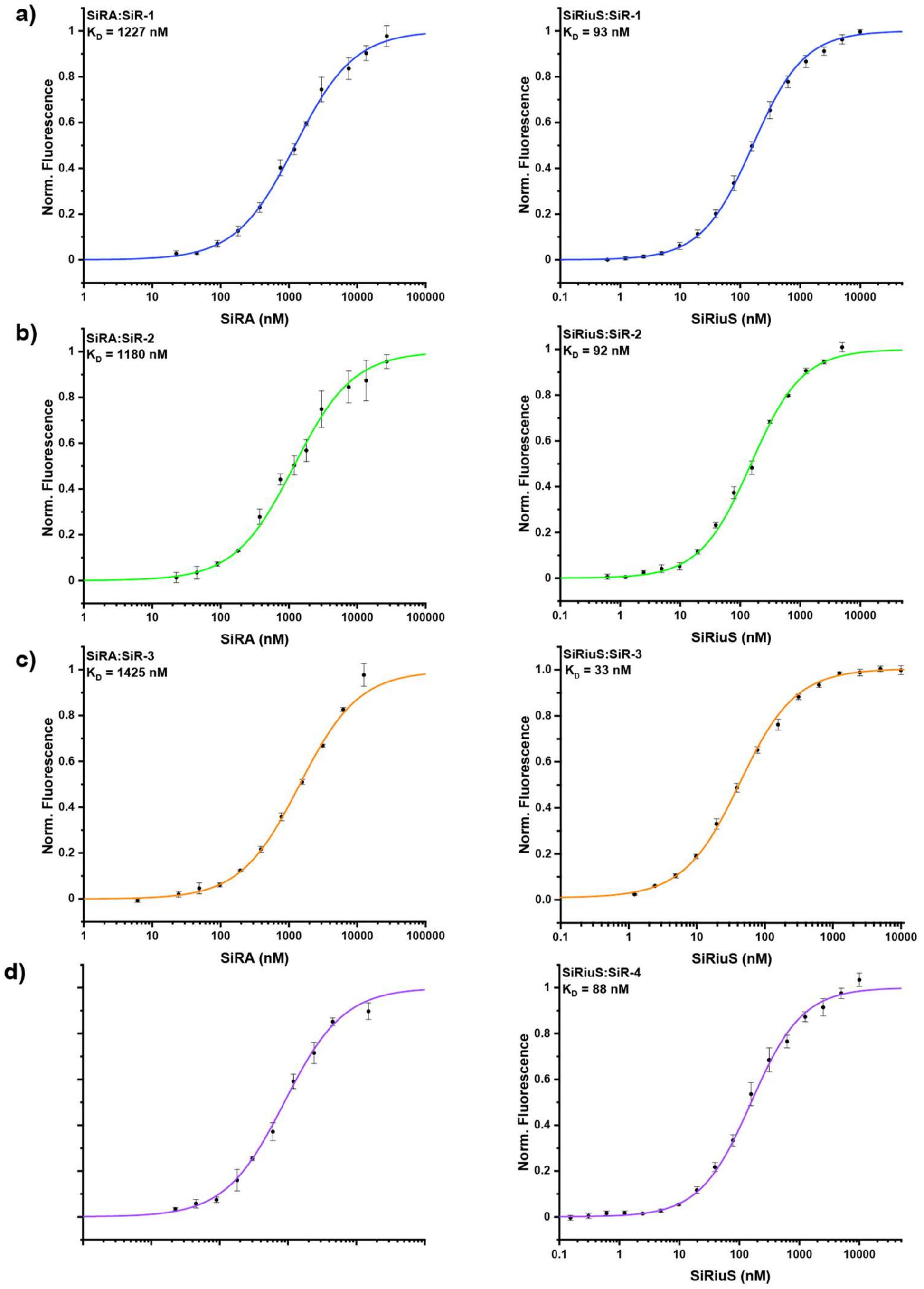

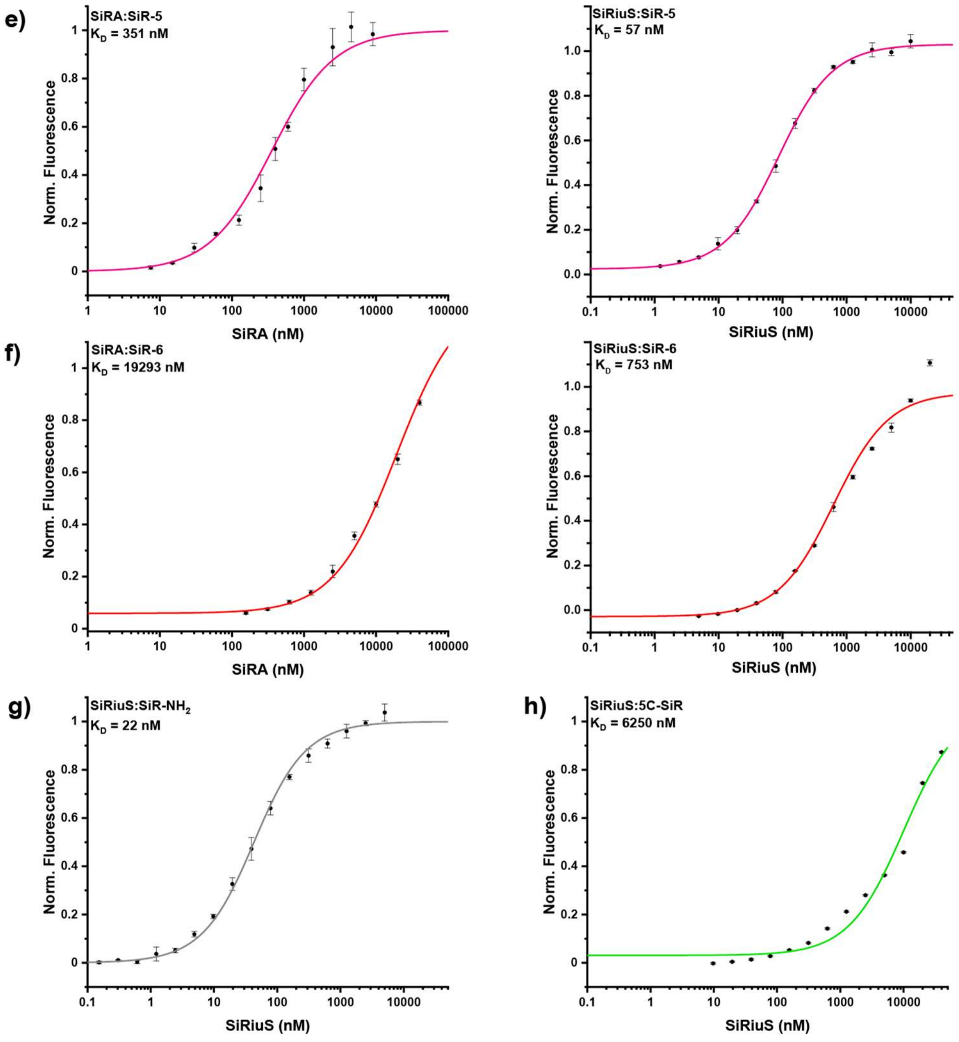

**Figure S9.** $K_D$ curves or SiR-R binding to SiRA or SiRiuS.a)-f) SiR-R (10 nM) binding to SiRA (left) or SiRiuS (right). g) SiR-$NH_2$ (10 nM) binding to SiRiuS. h) 5C-SiR (10 nM) binding to SiRiuS. Measurements were performed in ASB supplemented with 0.05% Tween 20 at 25 °C.

#### Excitation and emission spectra of SiR-R

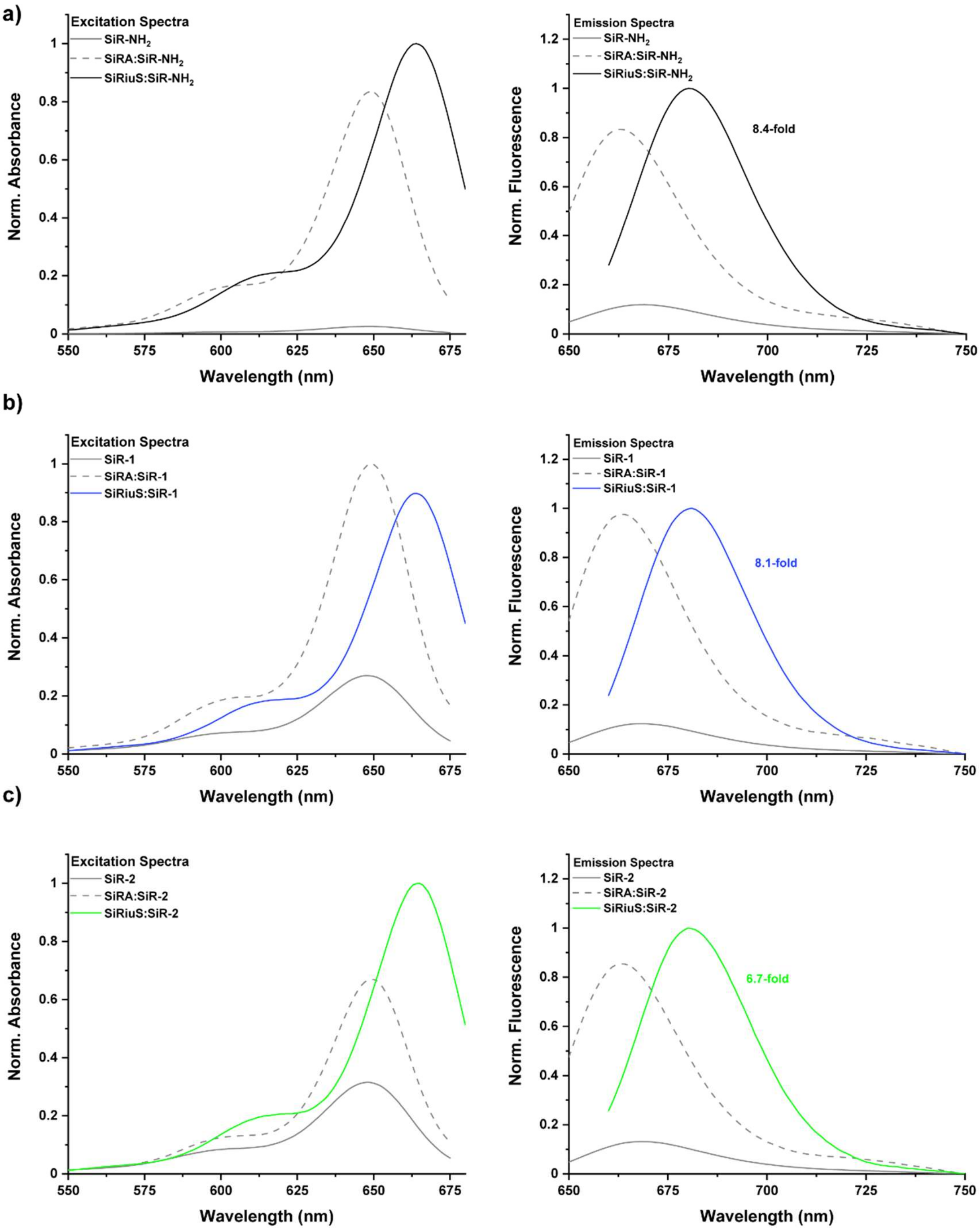

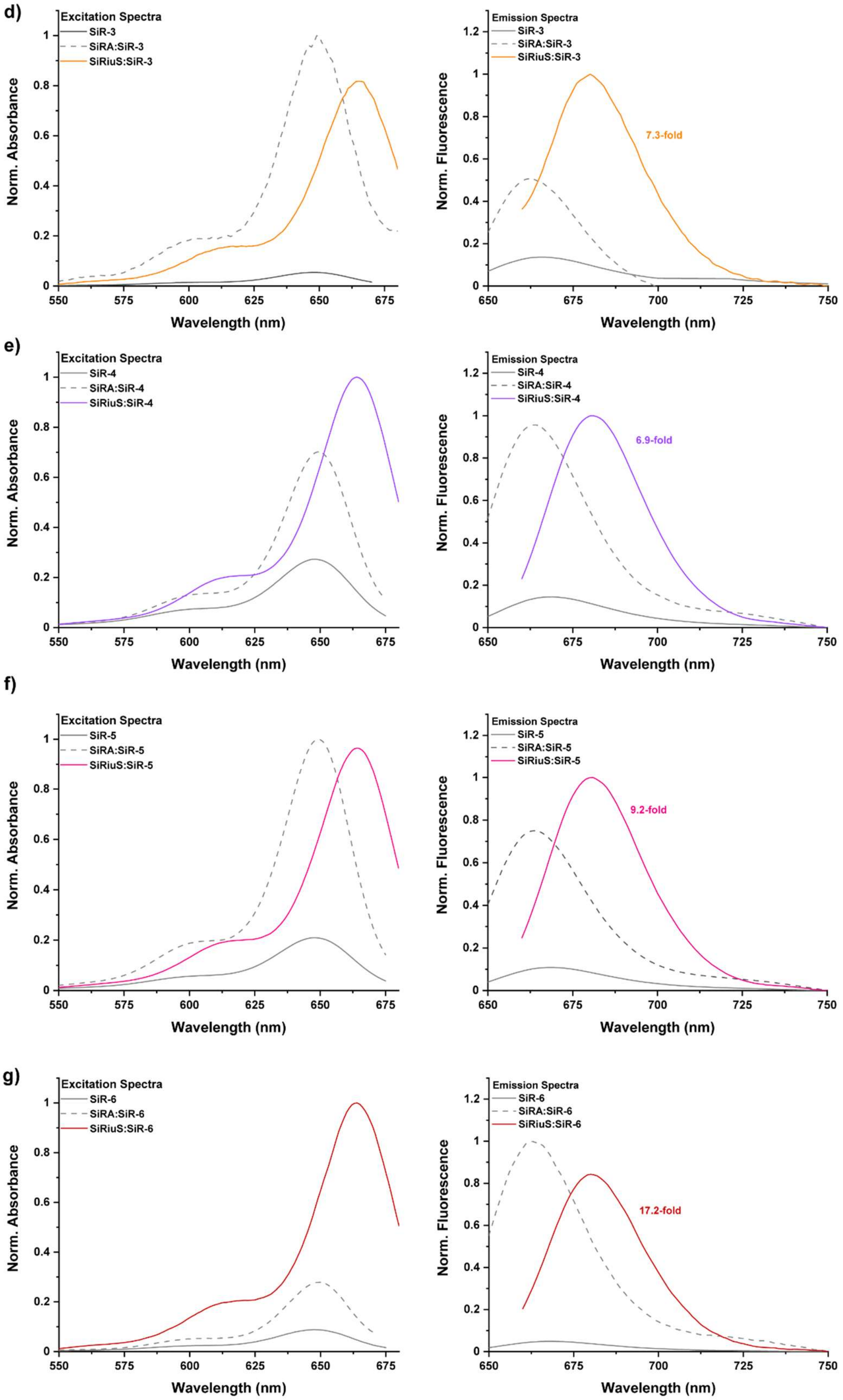

**Figure S10.** Normalized absorbance (left) and emission (right) spectra of free SiR-R dyes (grey solid line), in presence of SiRA (dashed grey line) or SiRiuS (solid colored line). Measurements were performed in ASB buffer at 25 °C using 200 nM of SiR-R and 20 µM of the respective aptamer.

#### $D_{50}$ values of SiR-R

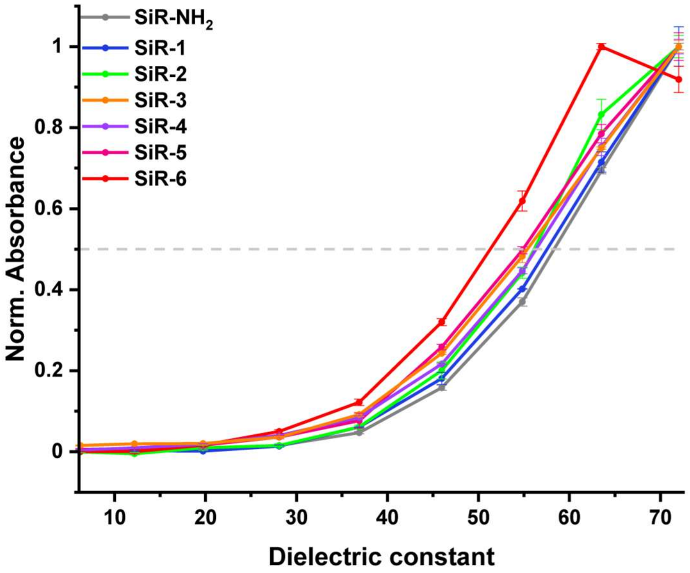

| Dye | $D_{50}$ |
|---|---|
| $SiR\text{-}NH_2$ | 58 |
| SiR-1 | 58 |
| SiR-2 | 56 |
| SiR-3 | 55 |
| SiR-4 | 56 |
| SiR-5 | 55 |
| SiR-6 | 51 |

**Figure S11.** Determination of $D_{50}$ values for the SiR-R dyes by measuring the absorbance in different dioxane:water ratios. The grey dashed line indicates the half-maximum absorbance at which $D_{50}$ is determined.

#### Colocalization experiments with LysoTracker

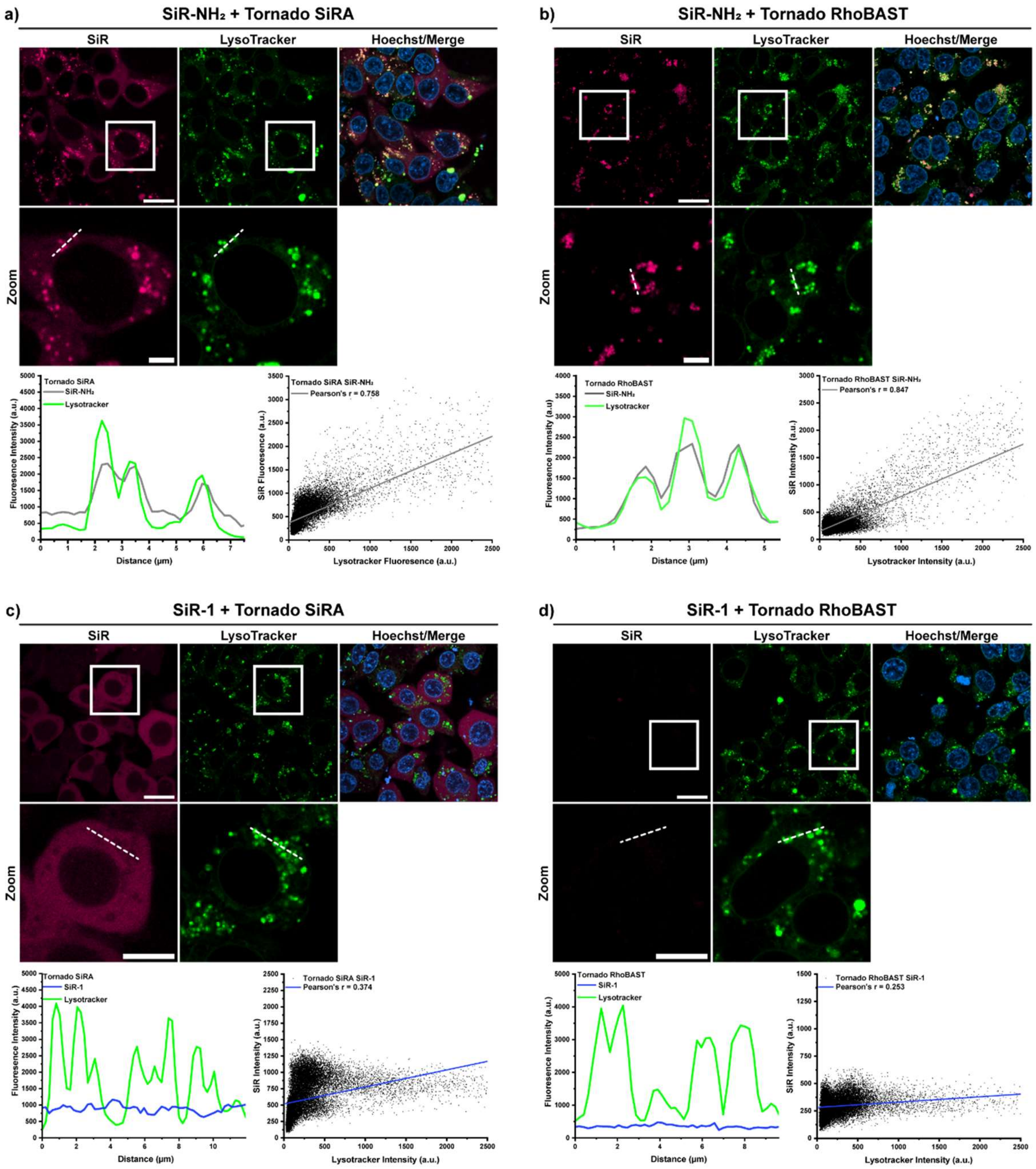

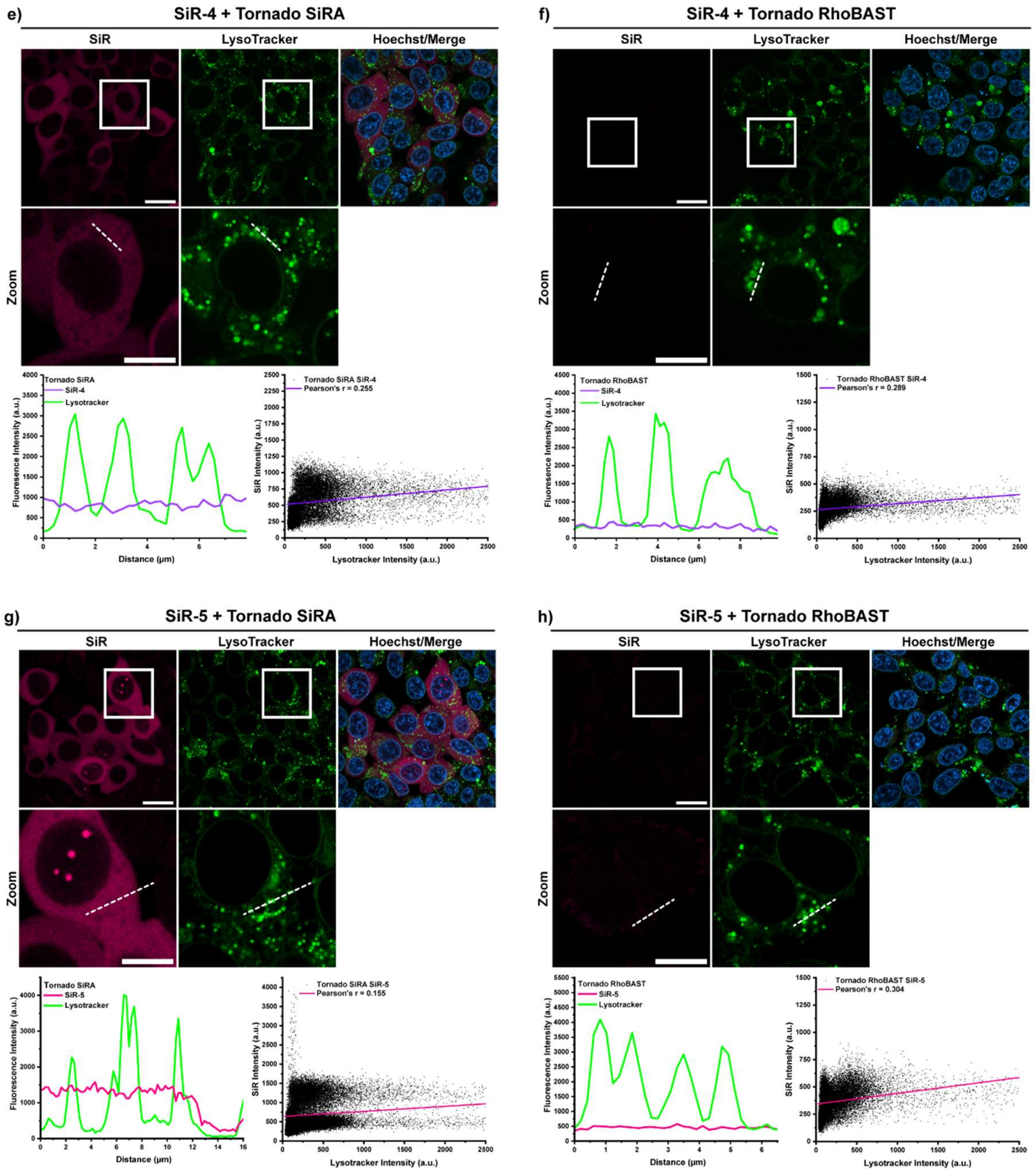

e)
SiR-4 + Tornado SiRA
SiR
LysoTracker
Hoechst/Merge
Zoom
Tornado SiRA
SiR-4
Lysotracker
Fluoresence Intensity (a.u.)
Distance (µm)
Tornado SiRA SiR-4
Pearson's r = 0.255
SiR Intensity (a.u.)
Lysotracker Intensity (a.u.)
f)
SiR-4 + Tornado RhoBAST
SiR
LysoTracker
Hoechst/Merge
Zoom
Tornado RhoBAST
SiR-4
Lysotracker
Fluoresence Intensity (a.u.)
Distance (µm)
Tornado RhoBAST SiR-4
Pearson's r = 0.289
SiR Intensity (a.u.)
Lysotracker Intensity (a.u.)
g)
SiR-5 + Tornado SiRA
SiR
LysoTracker
Hoechst/Merge
Zoom
Tornado SiRA
SiR-5
Lysotracker
Fluorescence Intensity (a.u.)
Distance (µm)
Tornado SiRA SiR-5
Pearson's r = 0.155
SiR Intensity (a.u.)
Lysotracker Intensity (a.u.)
h)
SiR-5 + Tornado RhoBAST
SiR
LysoTracker
Hoechst/Merge
Zoom
Tornado RhoBAST
SiR-5
Lysotracker
Fluoresence Intensity (a.u.)
Distance (µm)
Tornado RhoBAST SiR-5
Pearson's r = 0.304
SiR Intensity (a.u.)
Lysotracker Intensity (a.u.)

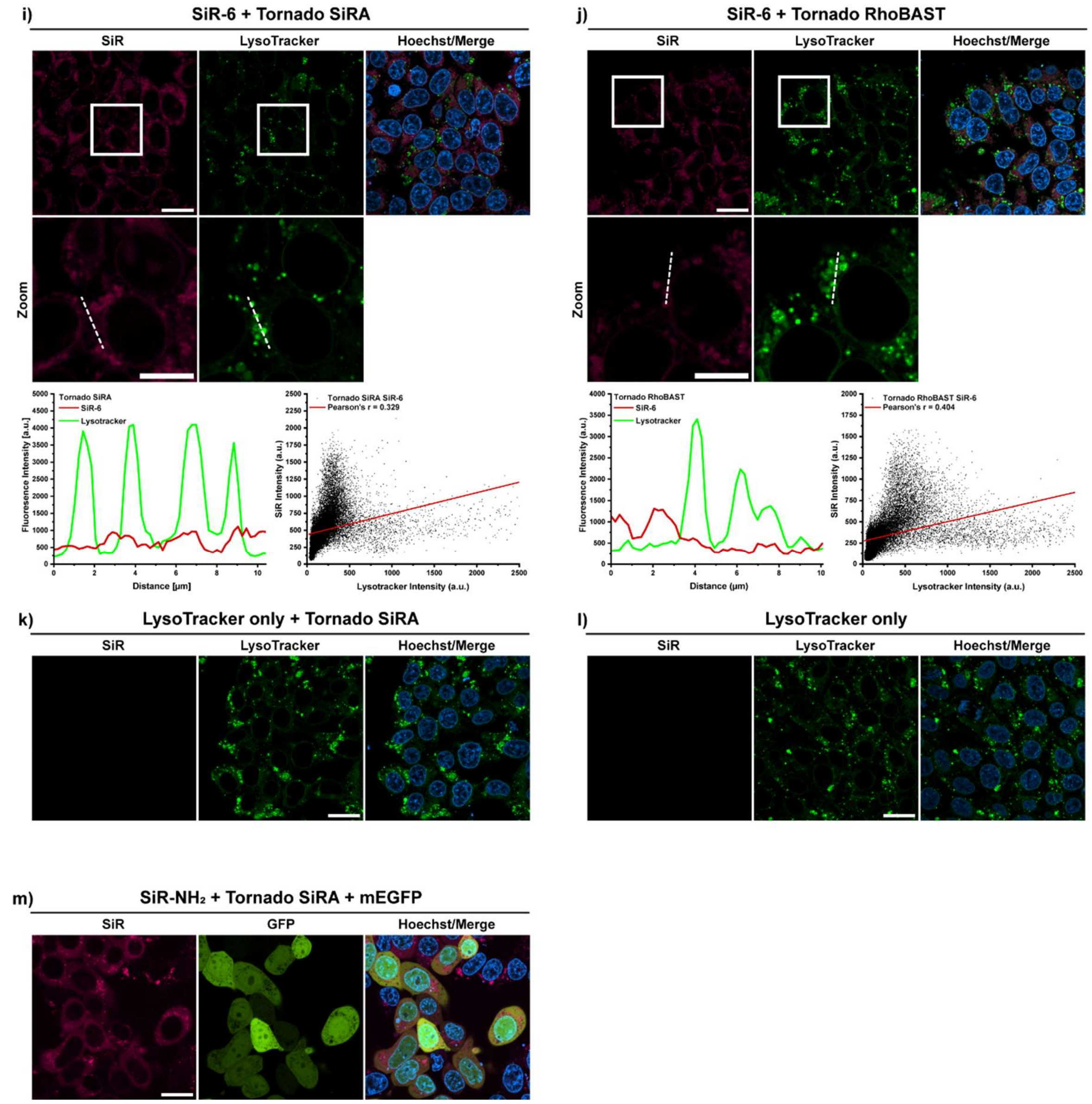

**Figure S12.** Colocalization experiments of SiR-R (500 nM) dyes in live HEK293T cells with LysoTracker Green (60 nM), showing line profiles and Pearson correlation. Cells were transfected with Tornado-SiRA (positive samples) and Tornado-Rhobast (negative control). a) Cells transfected with Tornado SiRA, incubated with SiR-$NH_2$; r = 0.785. b) Cells transfected with Tornado RhoBAST, incubated with SiR-$NH_2$, r = 0.847. c) Cells transfected with Tornado SiRA, incubated with SiR-1; r = 0.374. d) Cells transfected with Tornado RhoBAST, incubated with SiR-1; r = 0.253. e) Cells transfected with Tornado SiRA, incubated with SiR-4; r = 0.255. f) Cells transfected with Tornado RhoBAST, incubated with SiR-4; r = 0.289. g) Cells transfected with Tornado SiRA, incubated with SiR-5; r = 0.155. h) Cells transfected with Tornado RhoBAST, incubated with SiR-5; r = 0.304. i) Cells transfected with Tornado SiRA, incubated with SiR-6; r = 0.329. j) Cells transfected with Tornado RhoBAST, incubated with SiR-6; r = 0.404. k) Cells transfected with Tornado SiRA, incubated with LysoTracker Green (60 nM). l) Untransfected cells, incubated with LysoTracker Green (60 nM). m) Cells transfected with Tornado SiRA and mEGFP as transfection control, incubated with SiR-$NH_2$ (500 nM). Scalebars: 20 µm and 10 µm (zoom).

#### Colocalization experiments with SiR-6 and MitoTracker

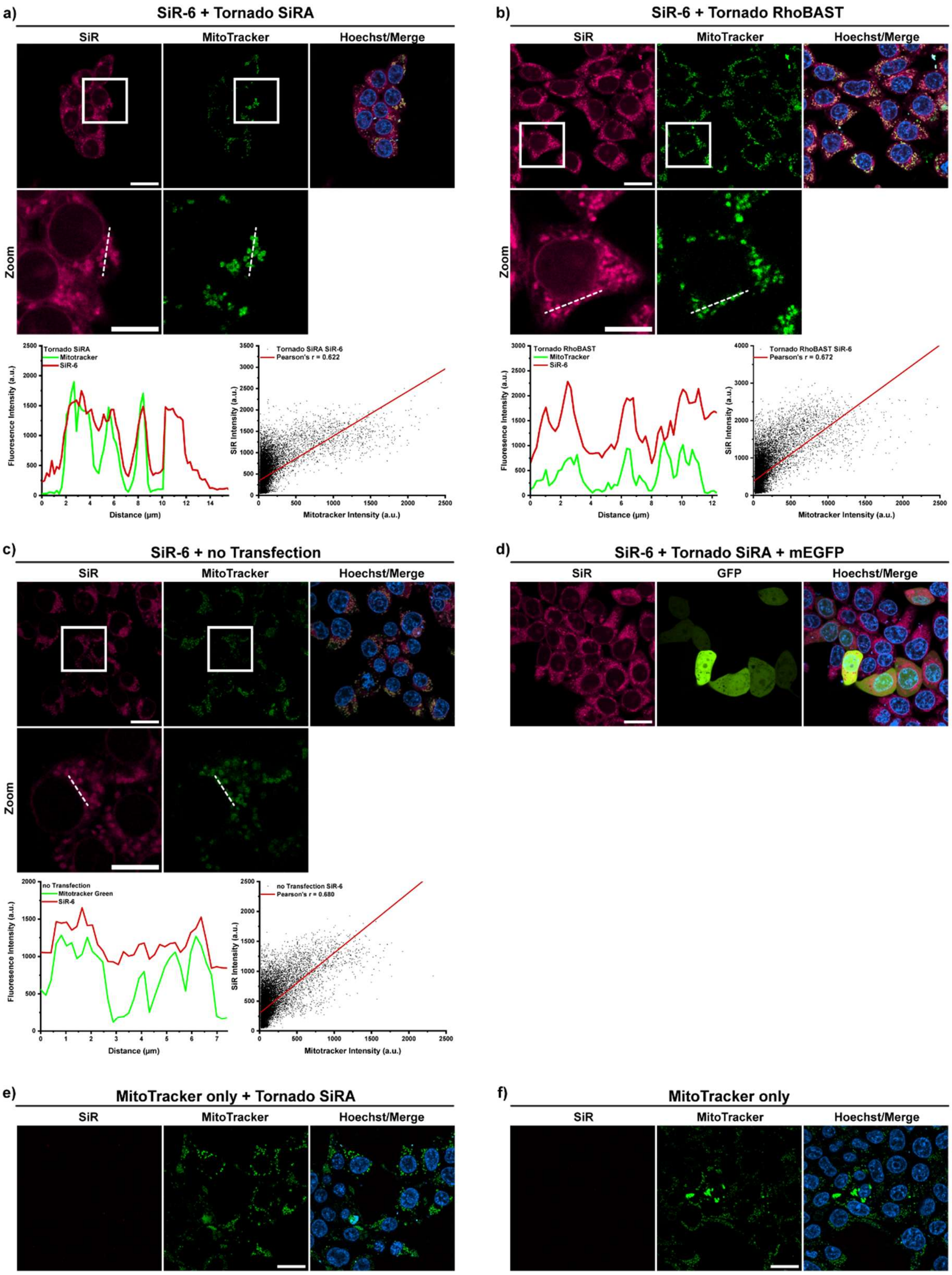

**Figure S13.** Colocalization experiments of SiR-6 (500 nM) in live HEK293T cells with MitoTracker Green (60 nM), showing line profiles and Pearson correlation. Cells were transfected with Tornado SiRA (positive samples) and Tornado RhoBAST (negative control). a) Cells transfected with Tornado SiRA, incubated with SiR-6; r = 0.622. b) Cells transfected with Tornado RhoBAST, incubated with SiR-6; r = 0.672. c) Untransfected cells incubated with SiR-6 and MitoTracker Green; r = 0.680. d) Cells transfected with Tornado SiRA and mEGFP as transfection control, incubated with SiR-6 (500 nM). e) Cells transfected with Tornado SiRA, incubated with MitoTracker Green (60 nM) only. l) Untransfected cells, incubated with MitoTracker Green (60 nM). Scale bars: 20 µm and 10 µm (zoom).

##### *In cellulo* dye titrations of SiR-R and SiRiuS in live HEK293T cells

a)

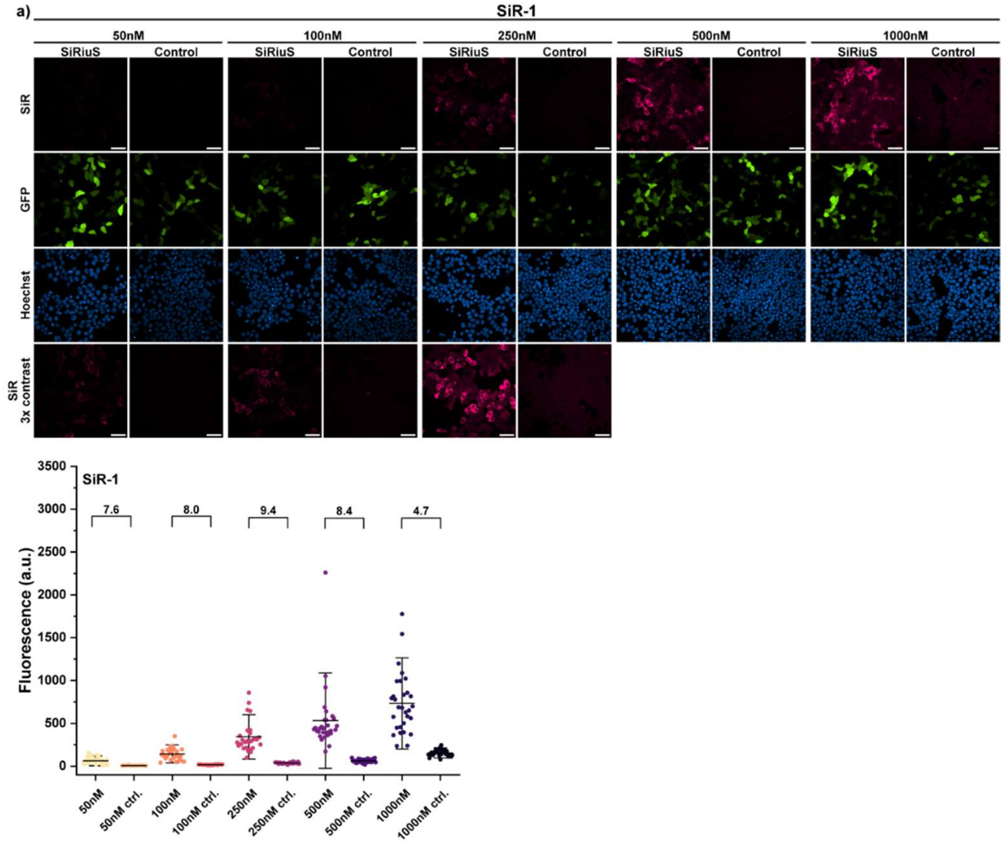

b)

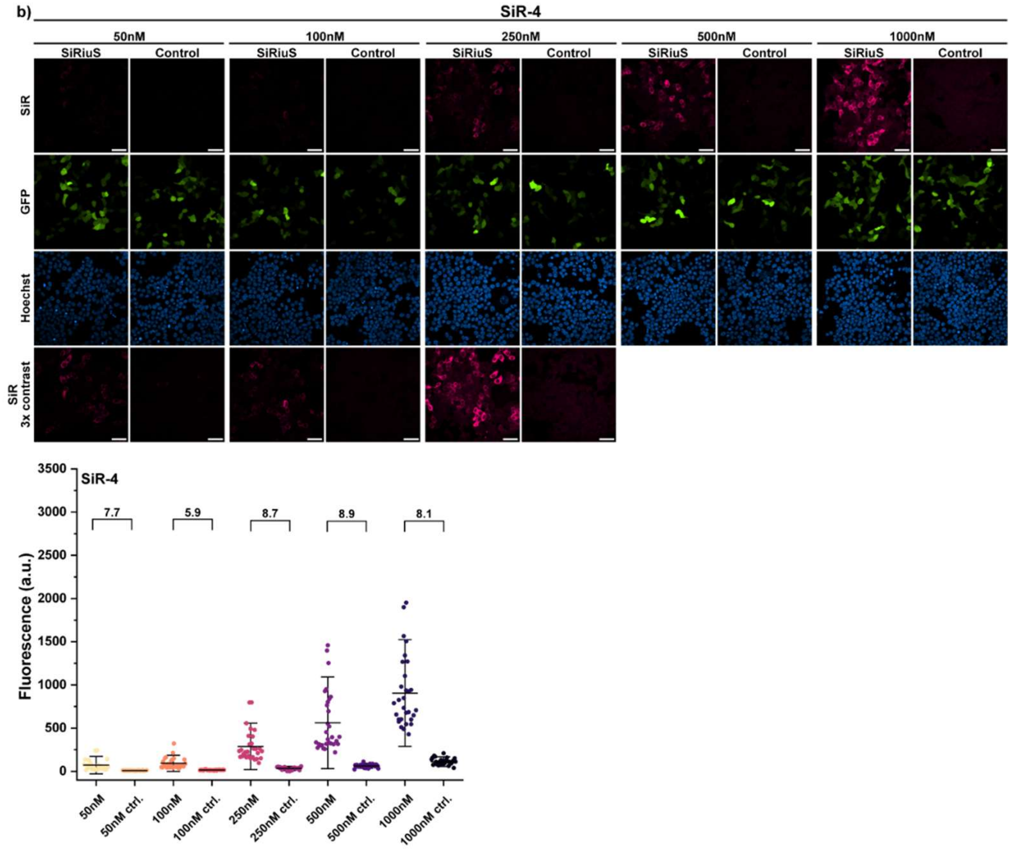

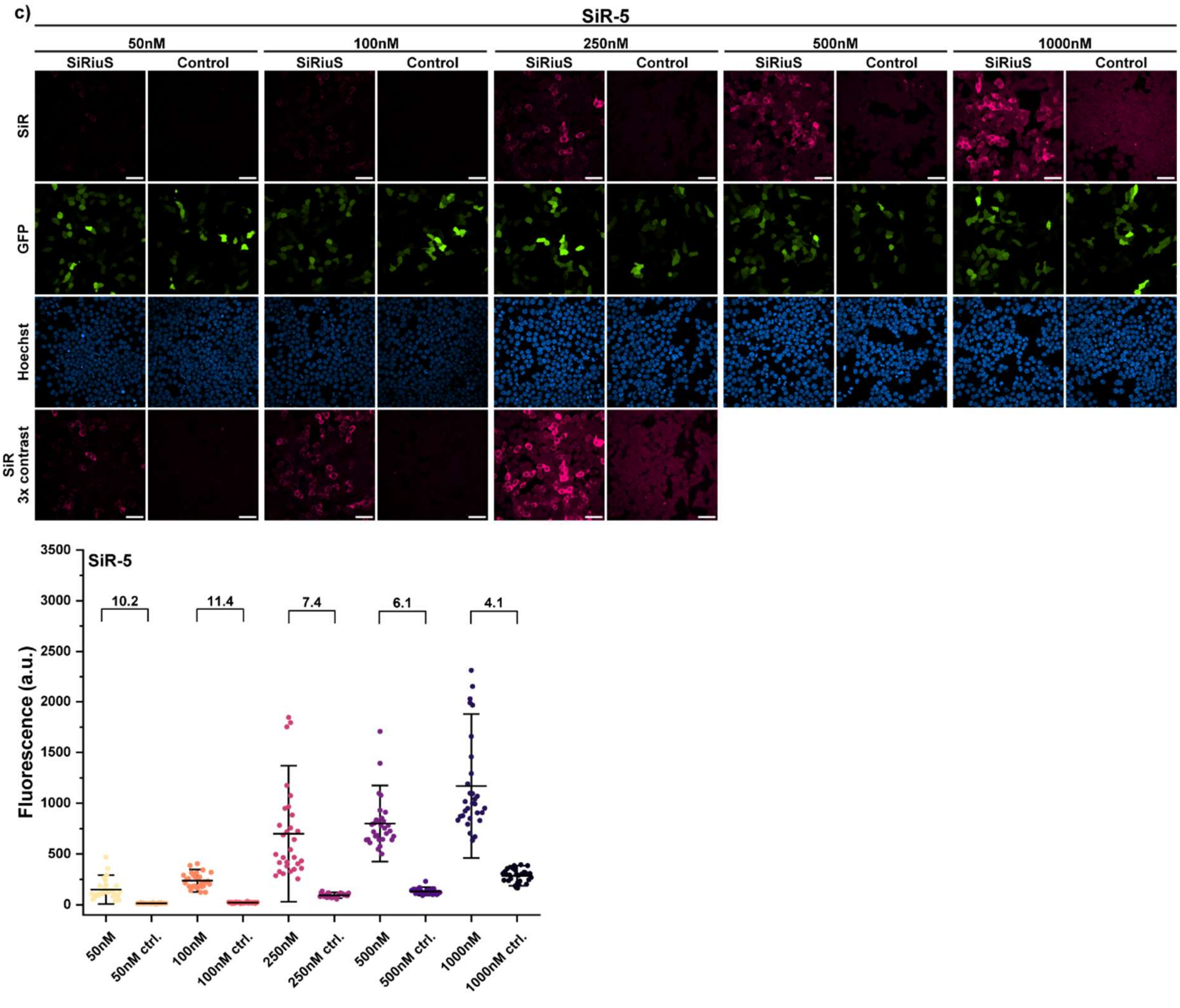

**Figure S14.** Dye titrations and quantification of the fluorescence turn-on of SiR-1, SiR-4 and SiR-5 in live HEK293T cells (*n* =30 cells) expressing either Tornado SiRiuS or Tornado RhoBAST (control). Cells were co-transfected with mEGFP. Imaging parameters and microscope settings were kept constant throughout the entire measurement series. Scale bars, 20 µm. a) Titration of SiR-1. Highest turn-on at 250 nM (9.4-fold). b) Titration of SiR-4. Highest turn-on at 500 nM (8.9-fold). c) Titration of SiR-5. Highest turn-on at 100 nM (11.4-fold).

##### *In cellulo* comparison of SiRA and SiRiuS with SiR-R in live HEK293T cells

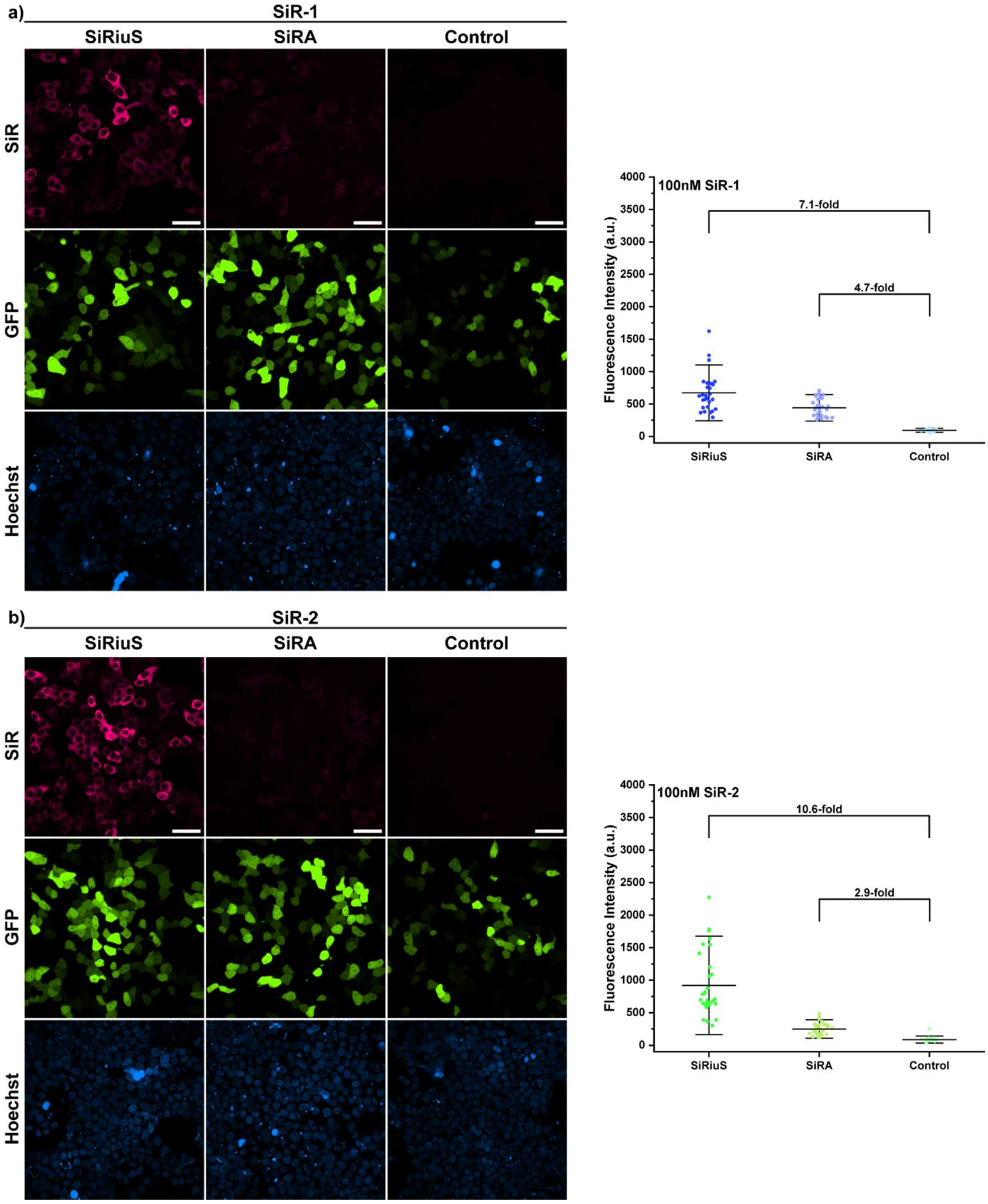

c)

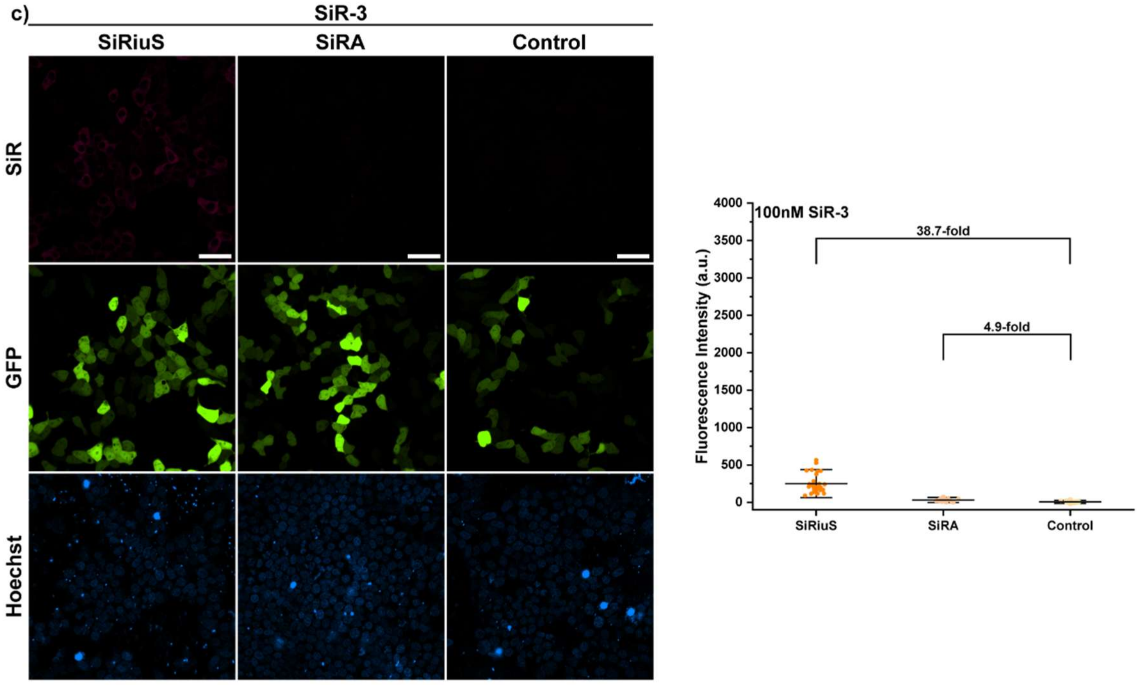
SiR-3
SiRiuS
SiRA
Control
SiR
GFP
Hoechst
100nM SiR-3
38.7-fold
4.9-fold
Fluorescence Intensity (a.u.)
4000
3500
3000
2500
2000
1500
1000
500
0
SiRiuS
SiRA
Control

d)

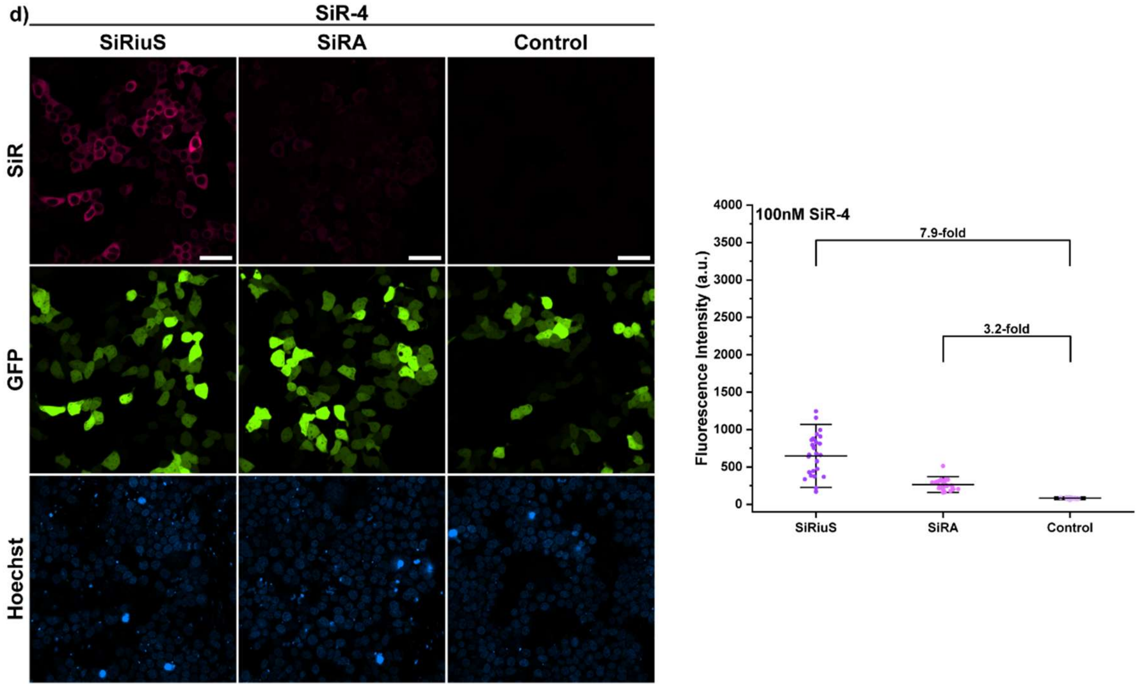
SiR-4
SiRiuS
SiRA
Control
SiR
GFP
Hoechst
100nM SiR-4
7.9-fold
3.2-fold
Fluorescence Intensity (a.u.)
4000
3500
3000
2500
2000
1500
1000
500
0
SiRiuS
SiRA
Control

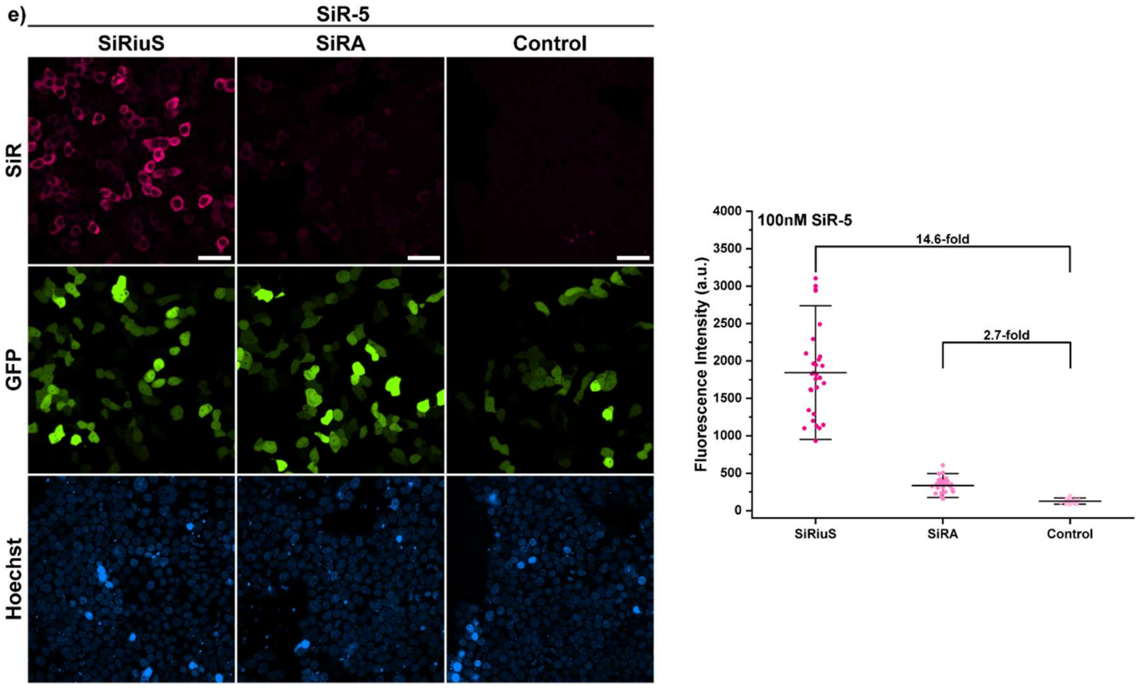

**Figure S15:** Comparison of live cell performance of SiRiuS and SiRA in HEK293T cells. Cells are expressing Tornado SiRiuS or Tornado SiRA. Cells expressing Tornado RhoBAST were used as negative control. All cells were co-transfected with mEGFP as transfection control and incubated with each dye (100 nM) for 45 min before the imaging session. Microscope settings were kept constant throughout the entire measurement series. To determine the fluorescence turn-on, $n$ =10 transfected cells were quantified in three independent experiments. Scale bars: 20 µm.

#### Live-cell imaging of SiRiuS:SiR-5 in different cell lines

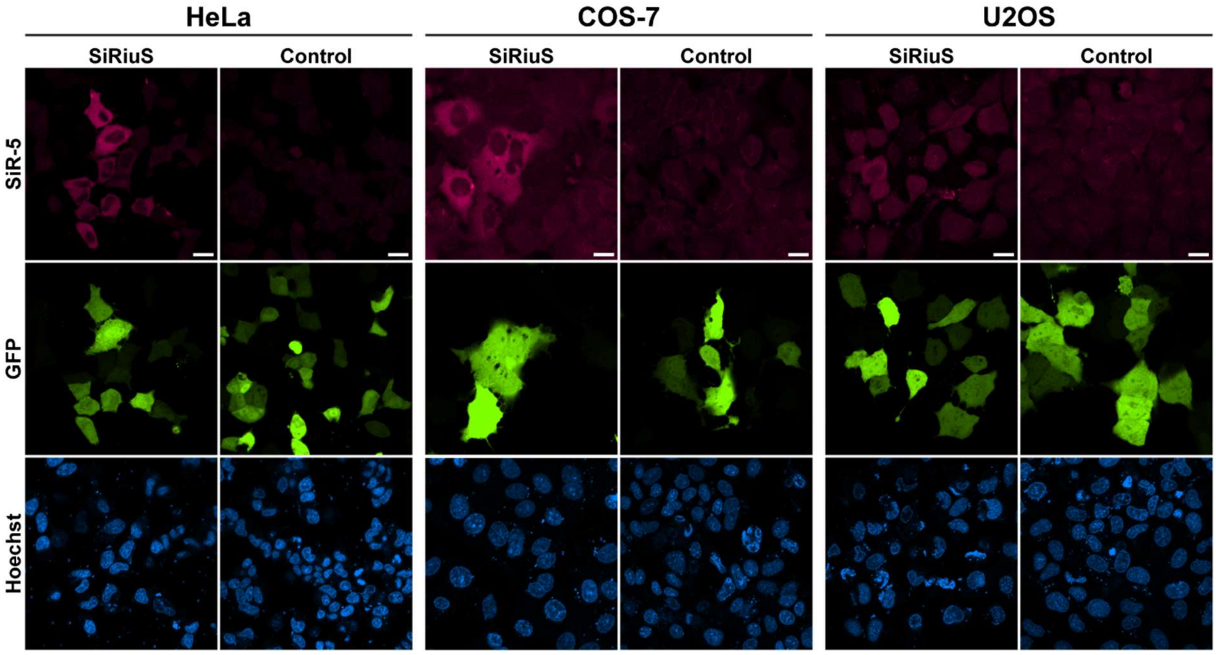

**Figure S16.** Confocal microscope images of live HeLa, COS-7 and U2OS cells expressing circular SiRiuS or RhoBAST (negative control) aptamer an incubated with SiR-5 (100 nM) for 1 h. All cell lines showed a specific fluorescent signal in the SiR-channel in positively transfected cells (mEGFP was co-transfected and used as transfection control). This experiment was performed in independent triplicates. Scale bars, 20 µm.

#### *In vitro* characteristics of SiRiuS:SiR-5

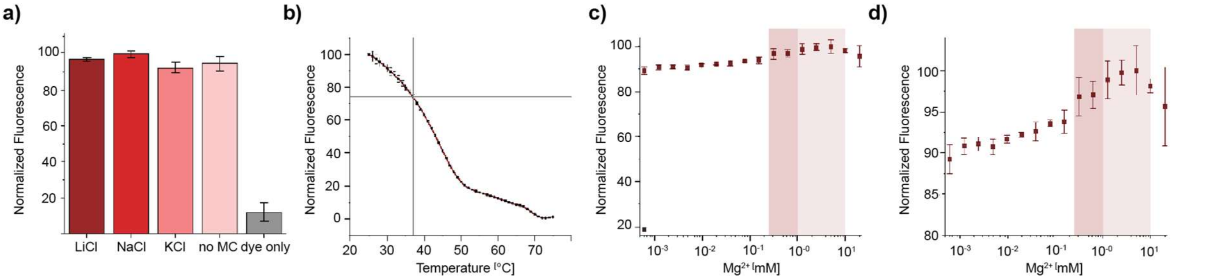

**Figure S17.** *In vitro* characterization of the SiRiuS:SiR-5 system. Dependence of the SiRiuS:SiR-5 complex fluorescence on a) monovalent cations; b) Temperature; c and d) Magnesium concentration. Panel d shows a zoom in of upper area of the graph in panel c).

#### Dye-uptake and bleaching of SiR-5 in live HEK293T cells

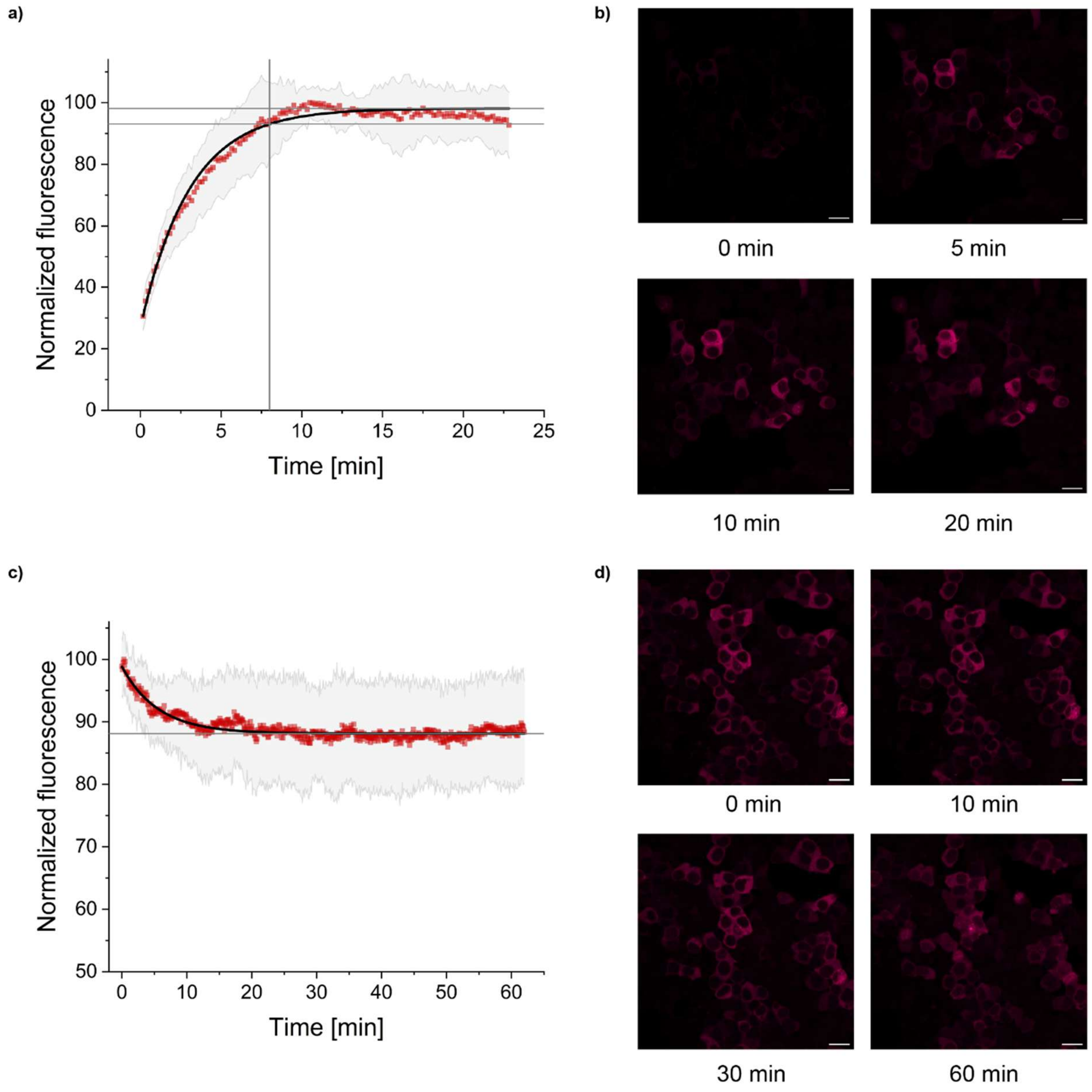

**Figure S18.** Dye uptake and photobleaching measurements. a) Dye uptake quantified from $n$ = 50 cells. 95% of the maximum intensity was reached at 8 minutes. b) 100 nM of SiR-5 was added and recording started immediately at 0.33 frames/s. c)

Bleaching curve quantifying n=50 cells. The curve asymptotically moves towards 87% maximum intensity. d) Bleaching of SiR-5 over 60 min with 0.33 frames/s and 200 µW. Scale bars, 10 µm.

#### Dual-color imaging of SiRiuS:SiR-5 and RhoBAST:SpyRho

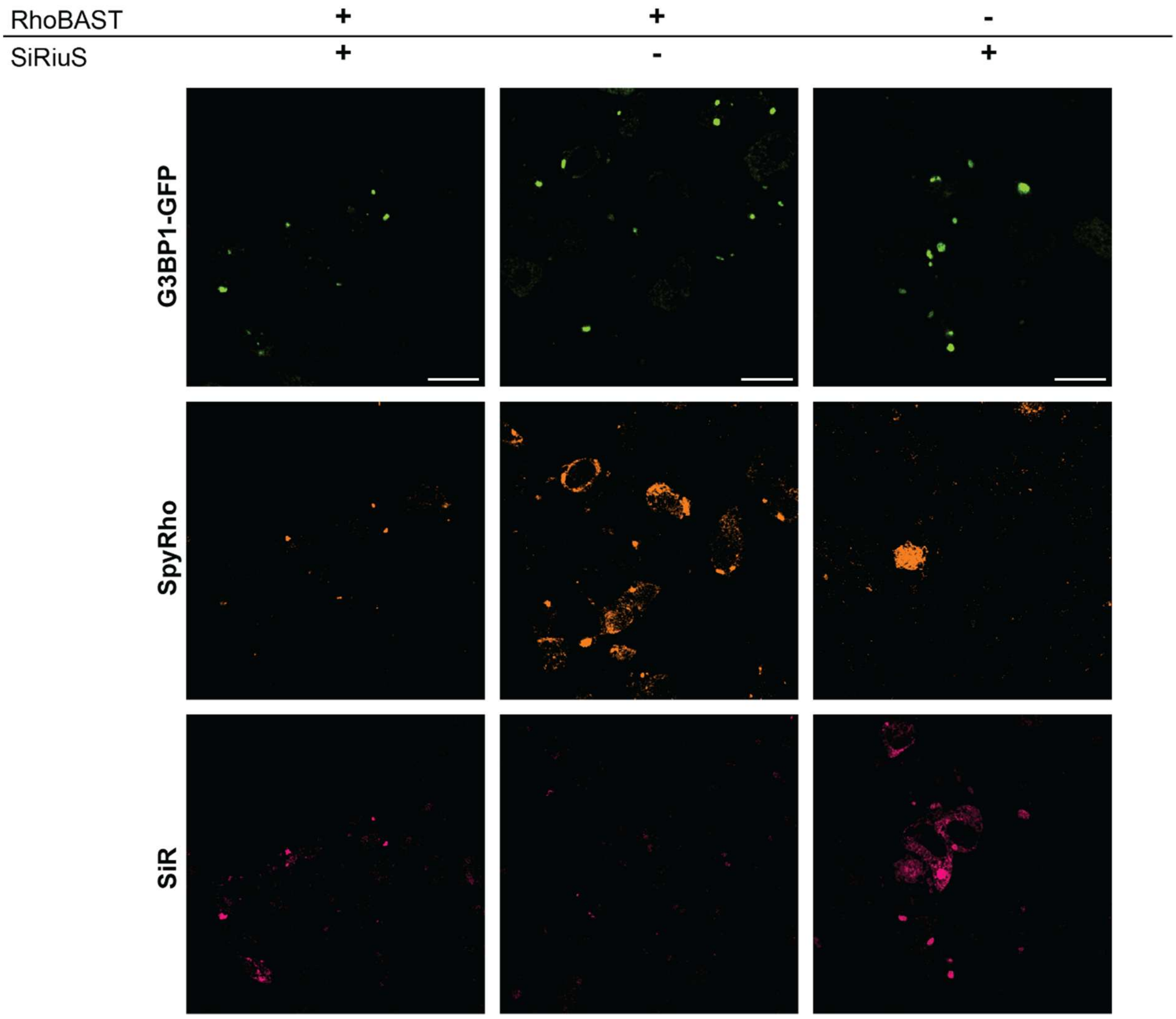

**Figure S19.** Orthogonality of SiRiuS and RhoBAST. Cells were transfected with either tagged or untagged versions of GAPDH-8xSiRius or mAzurite-8xRhoBAST or only one of the two, respectively, and G3BP1-GFP. Cells were fixed as indicated in method section 24 h after transfection and after 15 min incubation with 500 µM $NaAsO_2$. For each condition, 100 nM of both SpyRho and SiR-5 were added 30 min prior to imaging. Scale bars, 10 µm.

### Supplementary tables

#### RNA and DNA sequences

**Table S1.** Annotated sequences used in this work.

| Construct | Sequence 5’□ 3’ |
| --- | --- |
| **L2-1** | GGAGCUCAGCCUUCACUGCAAGAUGUGGACCAUUUAACUUGUAGACUGCUUCGGCAGGCGGCUGUUCCCUCAAGGGAACGCUUGGCACCACGGUCGGAUCCAC |
| **L2-2** | GGAGCUCAGCCUUCACUGCGACGUUUGAAAACGUCUAACACGAGUCUGCUUCGGCAGAGUCUGACGGUAUCCCGGCGGAUGUUGGCACCACGGUCGGAUCCAC |
| **L2-3** | GGAGCUCAGCCUUCACUGCAGCUGGCGUGGUGUAUAGUCUCCUGGCUGCUUCGACAGCUGUUUAAAUCGAUCUGGCGGACAUUGGCACCACGGUCGGAUCCAC |
| **L2-4** | GGAGCUCAGCCUUCACUGCACACUUUGAAAAGUGUCACUGCUAUGCUGCUUCGGCAGCAUCUCACGGUAUCCCGGCGGACGAGGCACCACGGUCGGAUCCAC |
| **L2-5** | GGAGCUCAGCCUUCACUGCCGCCCCCACCGGGUUUGAAAACCUGGCUGCUUCGGCAGUUGUAUCCUUUGGGGCUCGGCAAUUCGGCACCACGGUCGGAUCCAC |
| **L2-6** | GGAGCUCAGCCUUCACUGCACUCAGAUGGUUUUGAAAAGCCAUGACUGCUUCGGCAGUUGAUAUCUCCUGAACUCCUGAUGGAGGCACCACGGUCGGAUCCAC |
| **L2-7** | GGAGCUCAGCCUUCACUGCAGCGUCGGUGGUGUAUAGUCGGCGUCCUGCAGCAGUAUUGGCUCGGUUCGGCCCCCUCAUCGGCACCACGGUCGGAUCCAC |
| **L2-8** | GGAGCUCAGCCUUCACUGCGUGGUGUAUAGUUCCUGCGAUGGCAUCUGCUUCGGCAGAUAUACUGGGAUCCGUGACGAUCAUCGGCACCACGGUCGGAUCCAC |
| **L2-9** | GGAGCUCAGCCUUCACUGCUACGUACGGACUGAGCAAAGAGCUAACUACUUCGGCAGACCCUGCCGGACGGCGACCCCGGUCUGGCACCACGGUCGGAUCCAC |
| **L2-10** | GGAGCUCAGCCUUCACUGCUUCUCGUGAGUGCCCGCGCUCGCUGGUGUCGCGGAUCAGUUUUUGAAUCCACUCAAGGUUAACGAUAGGCACCACGGUCGGAUCCAC |
| **L1-1** | GGAGCUCAGCCUUCACUGCAGUAGGAUCAAGGUGCUGUAGGUGUUCUGCUUCGGCAGAGGUGUCCUACUGAUAGCUUCCGCCCGGCACCACGGUCGGAUCCAC |
| **L1-2** | GGAGCUCAGCCUUCACUGCAGUCGCCCGGGUGUAUAGGUAUCGACUGCUUCGGCAGAUAGCUCAAGAUGGACCGGUACCAUUGGCACCACGGUCGGAUCCAC |
| **L1-8** | GGAGCUCAGCCUUCACUGCGUGAUGUAUAGCCCCAGUGAACUAUCCUGCUUCGGCAGACAUAUGCUCCGGUCCGCCGGGCAUCGGCACCACGGUCGGAUCCAC |
| **L1-10** | GUUGCUGUUUGCGGUACACGCCGUGCUGGCGGCCUCAAUGUCGACCGUAAUGUAUCUCUUUGGUGUGGUG |
| **T7 Φ6.5 promoter** | TAATACGACTCACTATA’G |
| **L2-8+stem** | GGCCUGCGGAGCUCAGCCUUCACUGCGUGGUGUAUAGUUCCUGCGAUGGCAUCUGCUUCGGCAGAUAUACUGGGAUCCGUGACGAUCAUCGGCACCACGGUCGGAUCCACGCAGGC |
| **SiRiuS** | GCCUGCGGAGCUCAGCCGUCACUGCGUGGUGUAUAGUUCCUGCGACGGCAUAUACUGGGAUCCGUGACGAUCAUCGGCACCACGGUCGGAUCCACGCAGGC |
| **Syn. 1** | CAGGCAGGGAGCUCACGGCAGACUGCGUGGUGUAUAGUUCCUGCCUGCCGAUAAACUGGGAUGCGUGACGAUCAUCGGCACCACGGUCGCAUCCACCUGCCUG |
| **Syn. 2** | CCAGUGGGAGCUCAGCCCAGACUGCGUGGUGUAUAGUUCCUGCCUGGGCAUAUACUGCCAUGCGUGACGAUCAUCGGCACCACGGUCGCAUGGACCACUGGU |

| | |
|---|---|
| **Syn. 3** | GCGAGACCCACCUCACACGGCACUGCGUGGUGUAUAGUUCCUGCGCCGUGAUA UACUAGGAUGGGUGACGAUCAUCGGCACCACGGUCCCAUCCAGGUCUCGC |
| **L2-8 SHAPE** | GGCCTTCGGGCCAAGGCCTGCGACGAACGGAGCTCAGCCTTCACTGCGTGGTG TATAGTTCCTGCGATGGCATCTGCTTCGGCAGATATACTGGGATCCGTGACGA TCATCGGCACCACCGTTCGTCGGTCGGATCCACGCAGGCTCGATCCGGTTCGC CGGATCCAAATCGGGCTTCGGTCCGGTTC |
| **RT Primer 3'** | GATCCGGCGAACCGGATC |
| **RT Primer 5'** | CACGGATCCCAGTATATCTGCCG |
| **Primer A** | GGAGCUCAGCCUUCACUGC |
| **Primer B** | GGUCUGGCACCACGGUCGGAUCCAC |
| **SELEX pools with SalI and AatII** | AAAAAAGTCGACGGAGCTCAGCCTTCACTGCN26CTGCTTCGGCAG N26 GGCACCACGGTCGGATCCACGACGTCTTTTTT |
| **tRNA$_{Lys}$ with SalI and AatII sites** | GCCCGGATAGCTCAGTCGGTAGAGCAGCGTCGACGGTTTTGACGTCGA TGCGGGTCCAGGGTTCAAGTCCCTGTTCGGGCGCCA |
| **Primer A with SalI** | AAAAAAGTCGACGGAGCTCAGCCTTCACTGC |
| **Primer B with AatII** | GGCACCACGGTCGGATCCACGACGTCTTTTTT |

##### Candidate performance *in vitro*

**Table S2.** Results of the top ten candidates found for both performed FACS SELEX experiments. $K_D$ curves that did not reach the upper plateau or showed no turn on were deemed not determinable. Screening of $K_D$ values were done once.

| Candidate | $K_D$ | Turn-on |
|---|---|---|
| **L2-1** | 25 | 4.4 |
| **L2-2** | 208 | 5.3 |
| **L2-3** | 317 | 7.1 |
| **L2-4** | 274 | 5.7 |
| **L2-5** | 498 | 5.4 |
| **L2-6** | 782 | 5.1 |
| **L2-7** | 85 | 6.1 |
| **L2-8** | 53 | 5.6 |
| **L2-9** | n.d. | 1.1 |
| **L2-10** | n.d. | 1.9 |
| **L1-1** | 141 | 8.2 |
| **L1-2** | 354 | 7.6 |
| **L1-8** | 105 | 7.0 |
| **L1-10** | n.d. | 1.0 |

#### In vitro characteristics of SiR-R

**Table S3.** Photophysical properties of free SiR-R dyes in aqueous solution (1x ASB-T buffer).

| | $\lambda_{Ex}^{max}$ [nm] | $\lambda_{Em}^{max}$ [nm] | ε [$M^{-1}cm^{-1}$] | Φ | Brightness | $D_{50}$ |
|---|---|---|---|---|---|---|
| **$SiR-NH_2$[8]** | 649 | 666 | 27000 | 0.37 | 9940 | 58 |
| **SiR-1** | 647 | 668 | 23500 ± 3200 | 0.38 | 9000 | 58 |
| **SiR-2** | 647 | 668 | 23450 ± 2730 | 0.37 | 8650 | 56 |
| **SiR-3** | 648 | 668 | 28000 ± 4000 | 0.28 | 7840 | 55 |
| **SiR-4** | 648 | 669 | 24400 ± 4180 | 0.38 | 9200 | 56 |
| **SiR-5** | 648 | 669 | 24590 ± 3980 | 0.39 | 9565 | 55 |
| **SiR-6** | 648 | 668 | 3000 ± 150 | 0.15 | 462 | 51 |

**Table S4.** Photophysical properties of SiR-R dyes in presence of SiRA *in vitro*.

| | $\lambda_{Ex}^{max}$ [nm] | $\lambda_{Em}^{max}$ [nm] | ε [$M^{-1}cm^{-1}$] | Φ | Brightness | Turn-On | $K_D$ [nM] |
|---|---|---|---|---|---|---|---|
| **$SiR-NH_2$[8]** | 649 | 663 | 86000 | 0.59 | 50740 | 7 | 430 |
| **SiR-1** | 649 | 663 | 87050 ± 13000 | 0.61 | 53100 | 7.9 | 1227 ± 52 |
| **SiR-2** | 649 | 663 | 49740 ± 9200 | n.d. | n.d. | 6.5 | 1180 ± 79 |
| **SiR-3** | 649 | 663 | 51360 ± 4100 | n.d. | n.d. | 3.7 | 1425 ± 124 |
| **SiR-4** | 649 | 664 | 62710 ± 7200 | 0.61 | 38250 | 6.6 | 877 ± 60 |
| **SiR-5** | 649 | 664 | 125400 ± 6770 | 0.59 | 73990 | 6.9 | 351 ± 40 |
| **SiR-6** | 650 | 662 | 9460 ± 610 | n.d. | n.d. | 20.4 | 19293 ± 1580 |

**Table S5.** Photophysical properties of SiR-R dyes in presence of SiRiuS *in vitro*.

| | $\lambda_{Ex}^{max}$ [nm] | $\lambda_{Em}^{max}$ [nm] | ε [$M^{-1}cm^{-1}$] | Φ | Brightness | Turn-On | $K_D$ [nM] |
|---|---|---|---|---|---|---|---|
| **$SiR-NH_2$[8]** | 664 | 680 | 102990 ± 1440 | 0.63 | 64880 | 8.4 | 22 ± 2 |
| **5C-SiR** | 665 | 678 | n.d. | n.d. | n.d. | 14.6 | 6250 ±1290 |
| **SiR-1** | 664 | 681 | 78100 ± 3440 | 0.61 | 47640 | 8.1 | 93 ± 6 |
| **SiR-2** | 665 | 680 | 74300 ± 3200 | n.d. | n.d. | 6.7 | 92 ± 6 |
| **SiR-3** | 665 | 680 | 42000 ± 2800 | n.d. | n.d. | 7.3 | 33 ± 2 |
| **SiR-4** | 664 | 681 | 89400 ± 17150 | 0.60 | 53640 | 6.9 | 88 ± 3 |
| **SiR-5** | 664 | 681 | 120970 ± 13100 | 0.57 | 68950 | 9.2 | 57 ± 12 |
| **SiR-6** | 664 | 680 | 33880 ± 2760 | n.d. | n.d. | 17.2 | 753 ± 65 |
